# Collateral effects of bacteriophage resistance on antibiotic sensitivity in *Pseudomonas aeruginosa*

**DOI:** 10.64898/2026.10.10.758180

**Authors:** Laura R. Baars, Karsten Nagel, Laura B. Zwep, Tingjie Guo, J. G. Coen van Hasselt

**Affiliations:** Division of Systems Pharmacology and Pharmacy, Leiden Academic Centre for Drug Research, Leiden, The Netherlands

## Abstract

Collateral effect (CEs) of phage resistance can alter antibiotic sensitivity, leading to collateral sensitivity (CS). CS of phage resistance can potentially be used to design more effective phage–antibiotic combination treatments, yet insufficient knowledge on the occurrence of such phage resistance associated CS is currently available. Here, we characterized CEs across diverse *Pseudomonas aeruginosa* strains, antibiotics, and phages. For each of eight strains (seven clinical isolates and PAO1), we generated mutants resistant to phage PAO1-1 and, separately, mutants resistant to each of six antibiotics. Phage-resistant mutants were also generated using four additional phages (M32, NP1, NP3, and PNM), and double phage- and antibiotic-resistant mutants from one strain. We measured growth curves across antibiotic concentration ranges after which the area-under-the-curve (AUC) was quantified as a measure of bacterial burden. Dose-response curves for AUC against antibiotic concentrations were constructed to estimate a fold change in EC_50_ of mutants relative to their parental strains. PAO1-1-resistant mutants showed significant CS to colistin and polymyxin B. For other antibiotics, CEs varied by phage and parental strain: PAO1-1-resistant mutants showed CS to tobramycin, whereas mutants resistant to the other four phages tested were associated with CR. For double phage- and antibiotic resistant mutants we found CS to colistin and tobramycin relative to the parental strain. Resistance mutations occurred predominantly in genes involved in metabolism, virulence, and DNA replication. Antibiotic-resistant mutants also showed increased phage sensitivity, with the strongest effect in colistin-resistant mutants. In conclusion this study demonstrates collateral effects of phage resistance are most specific to both strain- and antibiotic-specific.

## 1 Introduction

Bacteriophage (phage) therapy is increasingly considered as an alternative to antibiotics given the global health crisis caused by antimicrobial resistance. This is particularly relevant for *Pseudomonas aeruginosa* (PA), a WHO priority pathogen and a major cause of chronic respiratory and wound infections, in which multidrug resistance is common and treatment options are increasingly limited.^1^ Despite its promise, one major concern regarding the application of phage therapy is the development of phage resistance.^2,3^ Phage resistance can arise quickly, often appearing within 6 to 48 hours.^4–7^

Development of resistance to an antimicrobial agent may lead to enhanced sensitivity towards a second antimicrobial agent, as an evolutionary trade-off. Such collateral sensitivity (CS) has been extensively studied as a consequence of resistance to antibacterial agents.^8–10^ CS effects arising from phage resistance that lead to enhanced antibiotic sensitivity have also been reported. For example in PA, CS in phage-resistant mutants has been described for polymyxins^11,12^, aminoglycosides^12–14^, fluoroquinolones^5,15,16^, tetracyclines^15,16^, cephalosporins^15–17^, and phosphonic acid antibiotics.^16,17^ In contrast, collateral resistance (CR), where antibiotic sensitivity is reduced has been reported for piperacillin, tobramycin, and meropenem.^14^ Illustrating the clinical relevance, emergence of phage resistance in patients has coincided with increased antibiotic sensitivity towards fluoroquinolones and aminoglycosides.^18,19^ These studies typically examine a single phage in a single strain background. As such, the variability and predictability of collateral effects (CEs) across PA strains, phages and antibiotic classes remain poorly characterised, which limits the generalisation of CS-based strategies for therapeutic use. Three aspects relevant to clinical application are largely under examined. First, it is unclear whether CEs are consistent across different phages, or whether they depend on the specific resistance mechanism selected by each phage. Second, phages are typically applied to strains that are already antibiotic-resistant, whereas CEs have mostly been characterised in antibiotic-sensitive backgrounds. Third, reciprocality of CEs, i.e., whether antibiotic resistance alters phage sensitivity, has received little attention despite its relevance for the design of sequential and combination regimens exploiting CS.

The current study sought to characterize phage-resistance associated CEs across multiple PA strains, phages, and antibiotics. Specifically, we aimed (i) to quantify CEs of phage resistance and identify patterns associated with antibiotic and parental strain background, (ii) to assess whether these effects are phage-specific, (iii) to determine whether phage resistance alters the antibiotic sensitivity of strains that are already antibiotic-resistant, and (iv) to assess whether antibiotic resistance can reciprocally alter phage sensitivity.

## 2 Materials and Methods

### 2.1 Bacterial strains, antibiotics and phages

Eight *Pseudomonas aeruginosa* (PA) strains were selected to represent a selection of strains with diverse antibiotic- and phage-sensitivity profiles, including one reference strain (DSM19880; PAO1) and seven clinical isolates: PA1947 (University Medical Center Groningen) and PA315, PA552, PA1583, PA1613, PA1948, and PA2108.^20^

Stock solutions for the antibiotics studied (aztreonam, ceftazidime, ciprofloxacin, colistin, polymyxin B, meropenem, and tobramycin) were prepared according to the dilutions and solvents described in Supplementary Table S1. Luria Broth (LB) medium was used for the antibiotic dilutions and the inoculum in the antibiotic sensitivity analysis. For tobramycin, we used Mueller-Hinton broth (MHB) supplemented with Mg^2+^ and Ca^2+^ (final concentrations: 10 *µ*g/mL and 20 *µ*g/mL, respectively; MHB+).

The lytic PA phages PAO1-1 (DSM112169; with recommended propagation host DSM19880), NP1 (GenBank accession number KX129925; Prof. Bruce Levin’s laboratory, Emory University, Atlanta, GA, USA), NP3 (GenBank accession number KU198331; Prof. Bruce Levin’s laboratory, Emory University, Atlanta, GA, USA), M32 (GenBank accession number KX711710.1; Phage Therapy Center in Tbilisi, Georgia), and PNM (Genbank accession number OP292288; Eliava Institute, Tbilisi, Georgia) were used in this study.

### 2.2 Mutant generation

#### 2.2.1 Phage-resistant mutants

All eight PA strains were sensitive to the PAO1-1 phage and were used to generate single phage-resistant mutants for this phage. A selection of these eight strains were used to generate single phage-resistant mutants against four other phages depending on their sensitivity profile (NP1 and PNM, n = 1; PAO1-1, n = 8; NP3 and M32, n = 5).

PA strains were co-cultured with the selected phage by mixing 200 *µ*L of overnight pre-culture (OD_600_ = 0.1) with 100 *µ*L of phage solution (10^8^ PFU/mL in LB) and 5 mL of molten soft LB agar (0.5%), which was then poured onto LB agar plates (1.5%). After gentle swirling and solidification, the plates were incubated overnight at 37 °C. Colonies that grew in the presence of phages were randomly collected and transferred, along with the parental strains, to 12 cm *×* 12 cm square LB agar (1.5%) plates containing 500 *µ*L phage stock (10^8^ PFU/mL) and 10 mL molten soft LB agar (0.5%).^21^ After a second overnight incubation at 37 °C, colonies that grew without visible lysis were considered candidate mutants. Approximately twelve colonies per strain–phage combination were selected and preserved in cryovials containing 10% glycerol at -80 °C for further testing. A bacterial strain was defined as phage-resistant when there was no lysis visible over all phage concentrations tested (10–10^8^ PFU/mL) or when the phage sensitivity was *>* 4-fold decreased. Phage titres for each phage specific to each parental strain were determined by plating PFUs. In total, 200 *µ*L of bacterial preculture (OD_600_ = 0.1), 100 *µ*L of phage stock (10^2^–10^3^ PFU/mL), and 5 mL of molten LB soft agar (0.5%) were mixed on LB agar, incubated overnight at 37°C, and PFUs manually counted to back-calculate the phage concentration.

#### 2.2.2 Antibiotic-resistant mutants

To investigate the collateral effect of antibiotic resistance on phage susceptibility, single antibiotic-resistant mutants were generated for six antibiotics. The bacterial strain PA1947 was arbitrarily chosen and evolved using a fluctuation assay. The overnight pre-culture was first diluted to a final cell density of approximately 5000 CFU/mL and distributed into 60 individual Eppendorf tubes containing 1 mL each. These cultures were plated onto selective agar containing one of the following antibiotics: aztreonam (AZT, 16 mg/L; 4*×*MIC), ceftazidime (CFT, 32 mg/L; 32*×*MIC), ciprofloxacin (CIP, 0.5 mg/L; 8*×*MIC), colistin (COL, 8 mg/L; 16*×*MIC), meropenem (MER, 8 mg/L; 32*×*MIC), or tobramycin (TOB, 16 mg/L; 32*×*MIC). Colonies that emerged on the selective plates were randomly picked as candidate mutants, and their minimum inhibitory concentrations (MICs) were subsequently determined (Supplementary table S2).

#### 2.2.3 Double antibiotic- and phage-resistant mutants

To assess whether pre-existing antibiotic resistance influences the collateral effects of phage resistance, antibiotic-resistant mutants (Section 2.2.2) were used to generate double antibiotic- and phage-resistant mutants. Three individual antibiotic-resistant mutants for each antibiotic were randomly selected to evolve additional phage resistance against PAO1-1, leading to double antibiotic- and phage-resistant mutants.

### 2.3 Whole Genome Sequencing

DNA is isolated from three PAO1-1 resistant mutants of each parental strain, bacterial cells were cultured in 5 ml of LB medium and incubated overnight at 37 °C with shaking at 150 rpm. Cell pellets were then harvested by centrifugation and DNA was extracted using the DNeasy Blood & Tissue Kit (Qiagen, Hilden, Germany) according to the manufacturer’s protocol. DNA concentrations were assessed using a NanoDrop spectrophotometer, ensuring that each sample contained at least 10 ng/*µ*L.

Whole-genome sequencing was performed by SeqCoast Genomics. For the eight parental strains, cell pellets were preserved in DNA/RNA Shield Stabilization Solution (Zymo Research) prior to extraction and hybrid sequencing (Oxford Nanopore and Illumina). For phage-resistant mutants, whole-genome sequencing was restricted to short-read sequencing (Illumina). Raw read quality was verified using a threshold of Q30 *≥* 90% for both long- and short reads (Supplementary Figures S15, S16, and S17). In addition to Q-score distributions, quality metrics were evaluated for read uniqueness, per-base quality as a function of read length, undetermined base calls (N content), and adapter contamination.

For the parental strains, short-read Illumina and long-read Nanopore data were combined using Unicycler in hybrid assembly mode, producing complete or near-complete chromosomal assemblies. Assembly quality control was performed for both long- and short-read data. For the Nanopore long reads, assembly metrics of the parental strains, including number of contigs, coding gene content, rRNA and tRNA counts, coding density, GC content, assembly length, and hypothetical proteins, were all within expected ranges, and no contamination from foreign DNA was detected (Data not shown). One exception was PA1948, which showed fragmented contig assembly (Supplementary Figure S18), however, as the longest contig reached 6,741,859 bp, this strain was retained for downstream analysis.

For the Illumina short reads, mapping of mutant reads to their respective parental reference genomes was satisfactory across all samples (Supplementary Figure S19), with the exception of two PA1613-derived mutants, where additional phage-derived reads were detected. As these phage sequences do not align to the parental reference genome, they were excluded from variant calling, and both mutants were therefore retained in the analysis.

Genome annotation was performed using Bakta (v1.12.0) with the full database (v6.0, February 2025). Mutations in phage-resistant mutants relative to their parental strains were identified using breseq (v0.39.0). Mutant assemblies were evaluated based on their percentage of reads mapping to the parental reference (threshold *>* 90%).

### 2.4 Pharmacodynamic growth profiling assays

We performed growth-profiling assays through measurement of optical density over time for all parental strains and antibiotic-, phage-, and double phage-antibiotic resistant mutants strains generated to quantify (collateral) changes in antibiotic- or phage sensitivity (Figure 1). For the PAO1-1 resistant mutants, three mutants per parental strain were characterised for each antibiotic, with the exception of colistin and polymyxin B, for which eight mutants per parental strain were characterised. To this end, 96-well plates contained dilution series of the test compound, along with wells without antibiotic or phage serving as growth controls. Plates were prepared one day prior to the assay and stored at 4 °C until use. For inoculation, overnight pre-cultures (OD_600_ = 0.1) were diluted 1:100 in LB or MHB+ medium, resulting in a final cell density of 10^6^ CFU/mL. The plates were incubated at 37 °C and optical density at 600 nm was measured over a 24-hour period using a plate reader equipped with a 96-well plate stacker without shaking (PerkinElmer EnVision). All conditions were tested in three biological replicates. To verify the desired inoculum concentration, CFUs were determined by plating a 1:1000 dilution (100 *µL*) onto LB agar, incubating for 24 hours at 37 °C, and manually counting colonies to derive the initial bacterial concentration.

**Figure 1.**
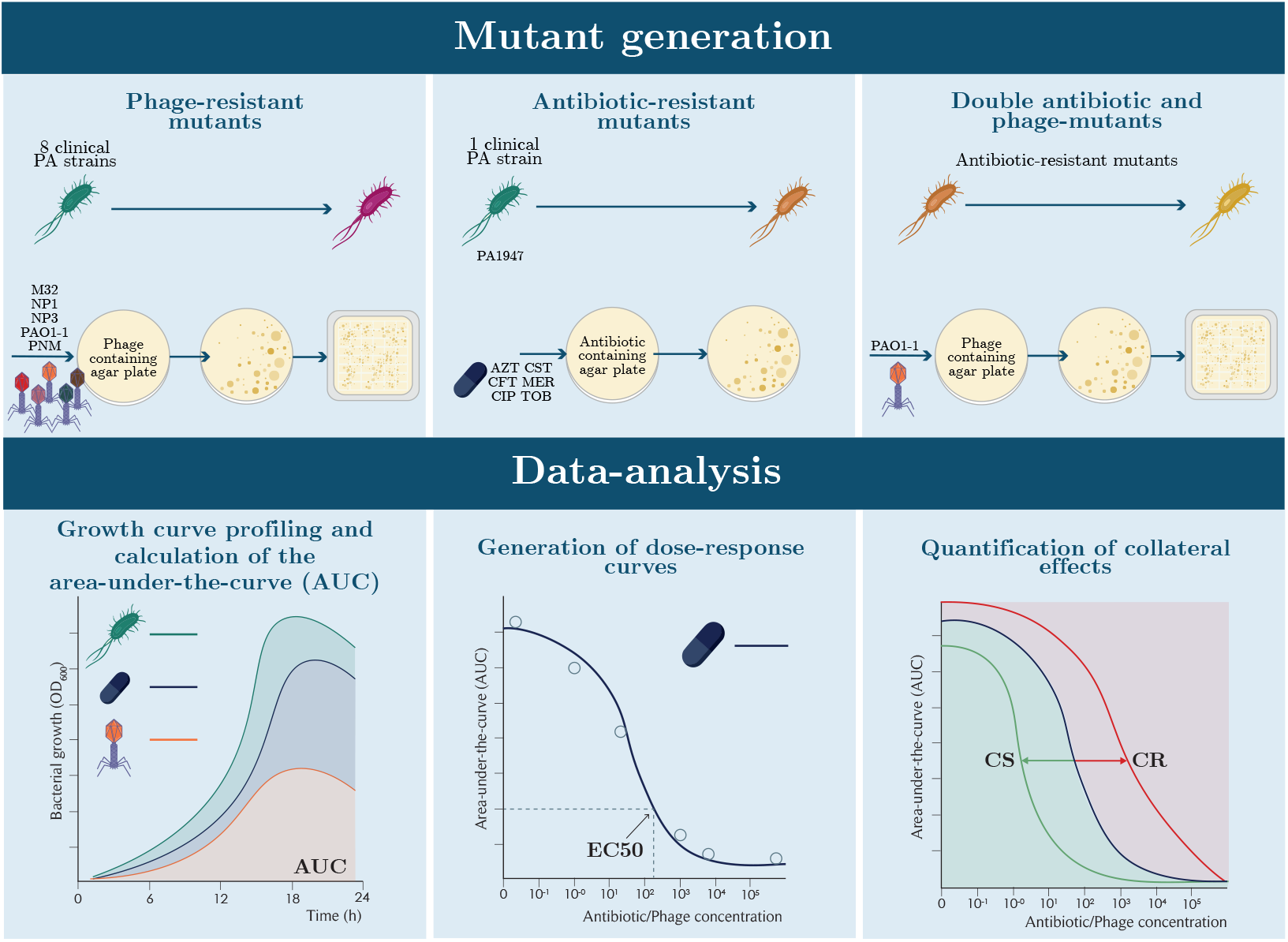
Workflow description of mutant generation, experimental approach and data analysis. Mutant generation. Phage-resistant mutants were obtained from eight independent strains by co-culturing with different phages (M32, NP1, NP3, PAO1-1, and PNM). After 24 hours, colonies appearing on the agar plates were isolated and further spread on square agar plates containing high-titre concentrations of the respective phage. Antibiotic-resistant mutants were generated from the parental strain PA1947 using a fluctuation assay with six antibiotics: aztreonam, ceftazidime, ciprofloxacin, colistin, meropenem, and tobramycin. Bacteria were plated on antibiotic-containing agar plates, and after 24 hours, the emerging colonies (mutants) were collected. Double mutants resistant to both antibiotics and phages were generated by exposing the antibiotic-resistant mutants to the PAO1-1 phage using the same co-culture procedure as for phage-resistant mutants. **Data-analysis.** OD_600_ growth-curve profiling is performed to quantify antibiotic sensitivity change in all mutants in relation to their parental strains, and to verify that phage resistance was retained in the mutants. The area-under-the-curve (AUC) is used to derive dose-response curves, and EC_50_ values were compared between mutants and parental strains by calculating the log_2_ fold change. A log_2_ transformed fold change below -0.5 indicates collateral sensitivity (CS), values between -0.5 and 0.5 indicate no collateral effect, and values above 0.5 indicate collateral resistance.

#### 2.4.1 Quantifying changes in antibiotic- and phage sensitivity

Hourly OD_600_ measurements from 6 to 24 hours, i.e., to capture active growth, were used to generate bacterial growth curves, from which the area under the curve (AUC) was calculated. AUC values were regressed against antibiotic (mg/L) or phage (PFU/mL) concentrations using a four-parameter log-logistic model, fitted separately for each parental strain, antibiotic, and phage with the drm function from the drc R package. For each fit, the half-maximum effective concentration (EC_50_) was extracted. The log2 fold-change in EC_50_ for each mutant relative its parental strain was then calculated (Equation 1) to estimate collateral effects.

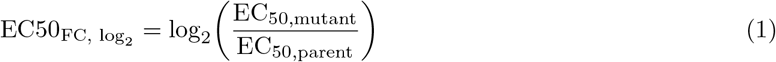

Collateral effects were classified as follows: collateral sensitivity (CS) if EC50_FC, log2_ *≤ −*0.5, and collateral resistance (CR) if EC50_FC, log2_ *≥* 0.5. The frequency of specific collateral effects was calculated for each antibiotic.

#### 2.4.2 Fitness cost of antibiotic- and phage resistance

The fitness of each individual mutant relative to its parental strain under phage-and antibiotic-free conditions was calculated using two different parameters; the maximum growth capacity and the growth rate. Maximum growth capacity was calculated as the percentage change in AUC_6-24_ of each mutant relative to its parental strain (without phage or antibiotic present). Maximum growth rate (*µ*_max_) was estimated as the maximal slope of the log-phase growth curve using the growthrates R package. The relative growth rate was calculated as the percentage change in *µ*_max_ of each mutant relative to its parental strain.

### 2.5 Statistical analysis

#### 2.5.1 Antibiotic sensitivity shifts in phage-resistant mutants

The log_2_-transformed fold change in EC_50_ relative to the parental strain (Equation 1), was used throughout as the metric of collateral effects. To estimate the mean EC50_FC, log2_ for each antibiotic and parental strain combination, while accounting for differences between parental strain backgrounds, we fitted a linear model with an antibiotic *×* parental strain interaction term using the base stats package in R:

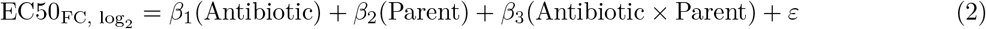

where *β*_1_ and *β*_2_ denote the categorical variables of antibiotic type and parental strain, respectively, *β*_3_ represents their interaction, and *ε* is the residual error term. The model was fitted without an intercept, which only removes the reference level for the first categorical variable (antibiotic), so each antibiotic coefficient directly represents its estimated mean EC_50,FC,log2_ , whereas the coefficients for parental strain and the antibiotic × strain interaction remain relative to the reference antibiotic and strain (Aztreonam and PA315, respectively). We performed post hoc pairwise comparisons using Tukey’s Honest Significant Difference (HSD) test, using the multcomp and emmeans packages, to identify which specific antibiotic–parental strain combinations differed significantly in EC50_FC, log2_ .

We also evaluated whether phage identity affected the estimated changes in antibiotic sensitivity in different phage-resistant mutants derived from phages NP1, NP3, M32, PAO1-1, and PNM (Section 2.4.1). Here, parental strain here was a confounding factor rather than a variable of interest, and unevenly represented across mutants due to initial phage sensitivities. We therefore fitted linear mixed models with a random intercept for parental strain (lme4 R package). For each antibiotic , we compared a null model containing solely the random intercept against a full model including phage identity as a fixed effect:

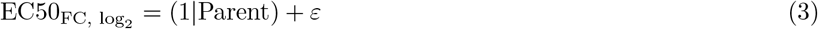

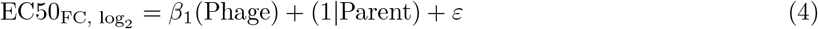

where *β*_1_ represents the effect of phage, (1*|*Parent) denotes the random intercept for parental strain, and *ε* is the residual error term. Whether adding phage improved model fit was evaluated with a likelihood ratio test comparing the nested models (Equations 3 and 4). For antibiotics with a significant phage effect, phage-level estimates were compared pairwise using Tukey’s HSD (multcomp, emmeans) packages.

To assess whether double antibiotic- and phage-resistant mutants differ from single antibiotic-resistant mutants in EC50_FC, log2_ , we compared the two groups for each antibiotic using a two-sample t-test (stats R package). Statistical significance was considered at *p <* 0.05, and the mean EC50_FC, log2_ values for each group were reported.

#### 2.5.2 Identification of genes associated with collateral effects

We evaluated associations between the EC50_FC, log2_ (Section 2.4.1) and occurrence of collateral sensitivity or resistance. Because the genetic basis of collateral effects may differ by antibiotic, we fitted a separate PLS regression model for each antibiotic, using the antibiotic’s EC50_FC, log2_ values as the response variable and genes with mutations presence/absence in comparison to its parent strain as predictors, using the pls package in R:

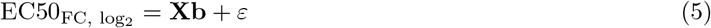

where **X** is a binary *n × p* gene presence/absence matrix across *n* mutant strains and *p* genes, **b** is the *p ×* 1 vector of PLS regression coefficients, and *ε* is the residual error term. Models were fitted using up to min(10*, p −* 1) components with leave-one-out cross-validation (CV) to determine the optimal number of components, and predictors were scaled to unit variance prior to fitting. Based on this, the first PLS component was retained for interpretation. Gene importance was assessed using the VIP score, and genes with a VIP score *>* 1 were considered to contribute meaningfully to the model. To assess the influence of individual observations on the regression fit, Cook’s distance was calculated from linear models fitted per antibiotic (Supplementary Figure S21). Genes were flagged as potentially influential when their mean Cook’s distance exceeded the conventional threshold of 4*/n*, where *n* is the number of unique genes per antibiotic.

## 3 Results

### 3.1 Antibiotic-specific collateral effects

We profiled collateral effects in antibiotic sensitivity towards seven antibiotics in PAO1-1 phage resistant mutants of eight different parental strains relative to their parental strain (Figure 2a, Supplementary Figure S8). Statistically signifiant CS effects for colistin and polymyxin B were found in a joint analysis across all tested mutant strains derived from different parental strains (p*<*0.001 and p=0.004, respectively; Supplementary Table S4). For other antibiotics, variability of collateral effects across different individual mutant strains was larger (Figure 2b). Indeed, collateral effects for a certain antibiotic differed with statistical significance between parental strains (ANOVA, p=0.049). Overall, we find that parental strain genetic background influenced the observed collateral effects for specific antibiotics (Supplementary table S5). The frequency of CS was found highest for colistin, polymyxin B, and tobramycin, with an incidence of 56.9%, 37.0%, and 21.9%, respectively, while CR was found highest among the *β*-lactams meropenem, aztreonam, and ceftazidime, with a frequency of 28.1%, 21.9%, and 12.5%, respectively. Finally, to evaluate if correlations exist between observed CS effects across antibiotics for specific PAO1-1 phage-resistant mutants, we calculated the Pearson correlation coefficients for all pairwise combinations (Supplementary Figure S13). The highest correlation was observed between the polymyxins, colistin and polymyxin B (r=0.92). Another high correlation was observed between the *β*-lactams aztreonam and ceftazidime (r=0.72).

**Figure 2.**
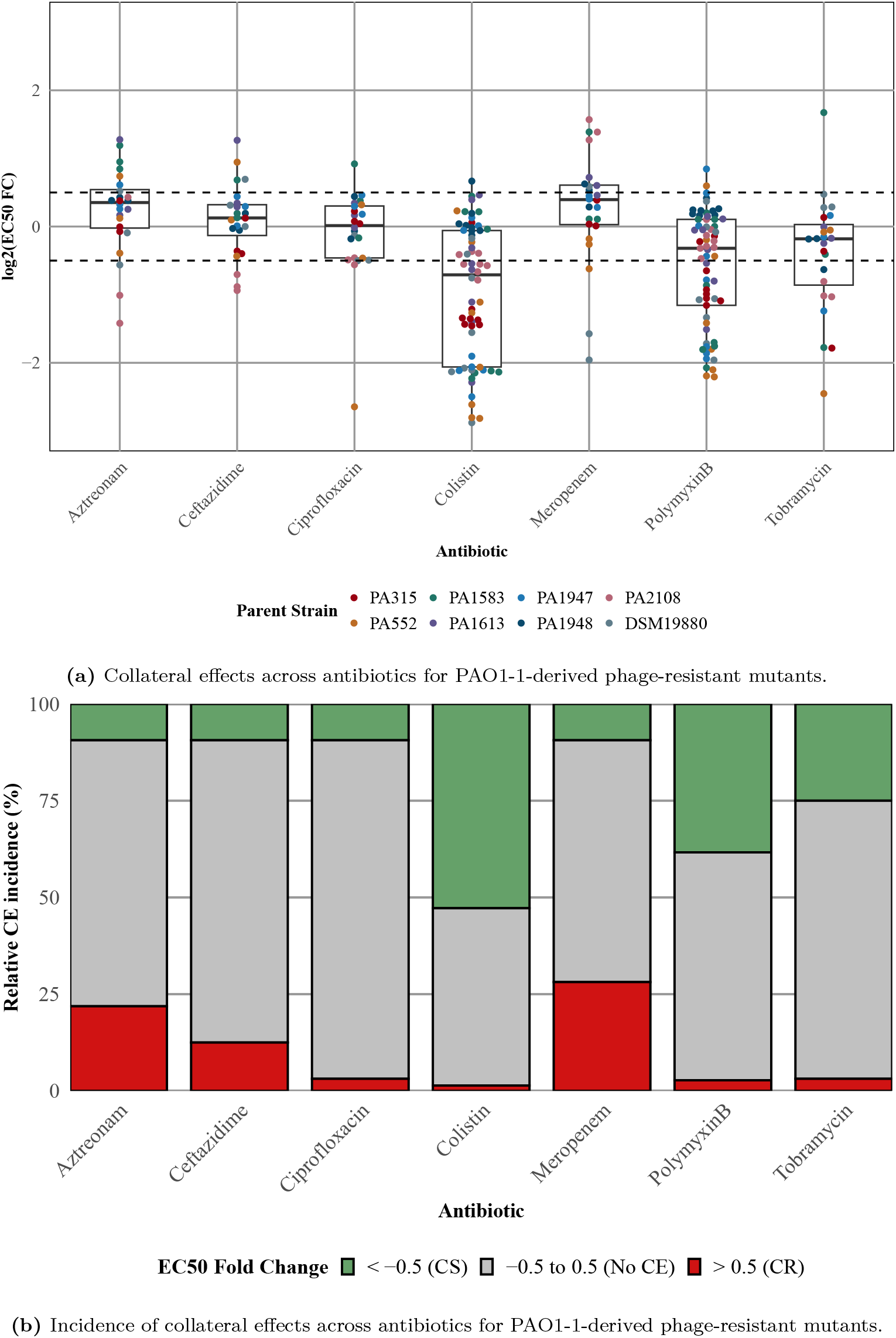
Collateral effects and their incidence in PAO1-1 single phage-resistant mutants. **A.** The boxplot shows the median, range, and standard error of EC50_FC, log2_ across different antibiotics for PAO1-1-derived phage-resistant mutants, categorized as collateral sensitivity (CS; *≤* -0.5), no collateral effect (CE; between -0.5 and 0.5), or collateral resistance (CR; *≥* 0.5). Three independent mutants per parental strain (indicated by colour; n=8) were measured for each antibiotic, except colistin and polymyxin B, for which additional mutants were included due to their larger effects. Each data point represents the mean EC50_FC, log2_ of a single mutant, averaged over three biological replicates. **B.** The relative incidence (%) of collateral effects across antibiotics, based on categorizing each mutant’s EC50_FC, log2_ as CS (green), no CE (grey), or CR (red).

### 3.2 Phage-specific collateral effects

To evaluate potential differences in collateral effects towards antibiotics across different phages, we profiled mutants derived from up to eight parental PA strains for each of five phages against six antibiotics relative to their parental strain (Supplementary Figures S1, S2, S3, S5, S4, S8, and S10). In terms of overall frequency, M32- and NP3-resistant mutants displayed more similar CE patterns overall, with occasional collateral sensitivity to ciprofloxacin, ceftazidime, and colistin, whereas NP1- and PNM-resistant mutants differed more markedly, showing phage- and strain-specific patterns of collateral sensitivity and -resistance (Figure 3). When evaluating these effects specifically within antibiotics, statistically significant differences in EC50_FC, log2_ between phage-resistant groups for ceftazidime, colistin, and tobramycin (p*<*0.001, p=0.040, p*<*0.001, respectively, likelihood ratio test; Figure 4 and Supplementary Table S6). For ceftazidime, mutants resistant to the phages NP1, NP3, and PNM showed a shift in EC50 relative to M32 consistent with a trend towards CS (M32 vs. NP3: p=0.002, M32 vs. PNM: p*<*0.001, and M32 vs. NP1: p=0.725). Similarly, PAO1-1-resistant mutants showed no meaningful shift relative to M32 (p=0.999), and differed significantly from NP3- and PNM-resistant mutants (p*<*0.001 for both; Supplementary Table S7; Figure 4). For tobramycin, solely PAO1-1-resistant mutants showed CS, exhibiting significantly greater sensitivity than mutants resistant to any of the other four phages (M32, NP1, NP3, and PNM: p*<*0.001, p=0.049, p*<*0.001, and p*<*0.001, respectively; Supplementary Table S7; Figure 4). For colistin, despite a significant overall phage effect, post hoc pairwise comparisons did not identify significant differences between specific phage groups (Supplementary Table S7).

**Figure 3.**
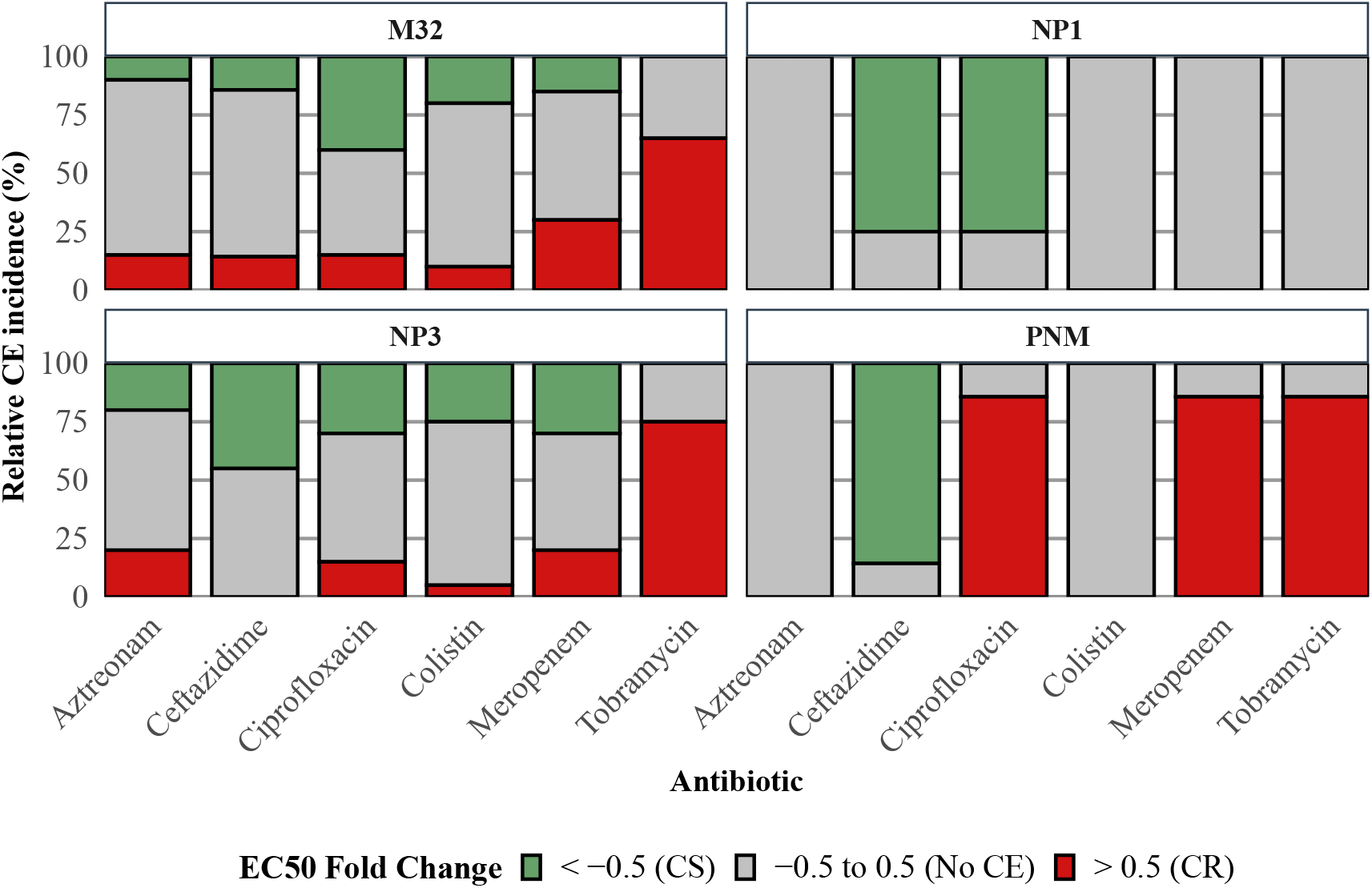
Incidence of collateral effects in single phage-resistant mutants against six different antibiotics. The relative incidence (%) of specific collateral effects across different antibiotics in different phage-resistant mutants. After log_2_ transformation, values *≤ −*0.5 indicate collateral sensitivity (CS; green), and values *≥* 0.5 indicate collateral resistance (CR; red), while no collateral effects (CE; grey) lie between -0.5 and 0.5. The parental strain from which each mutant was derived varied by phage due to differences in sensitivity (NP1 and PNM, n = 1; PAO1-1, n = 8; NP3 and M32, n = 5).

**Figure 4.**
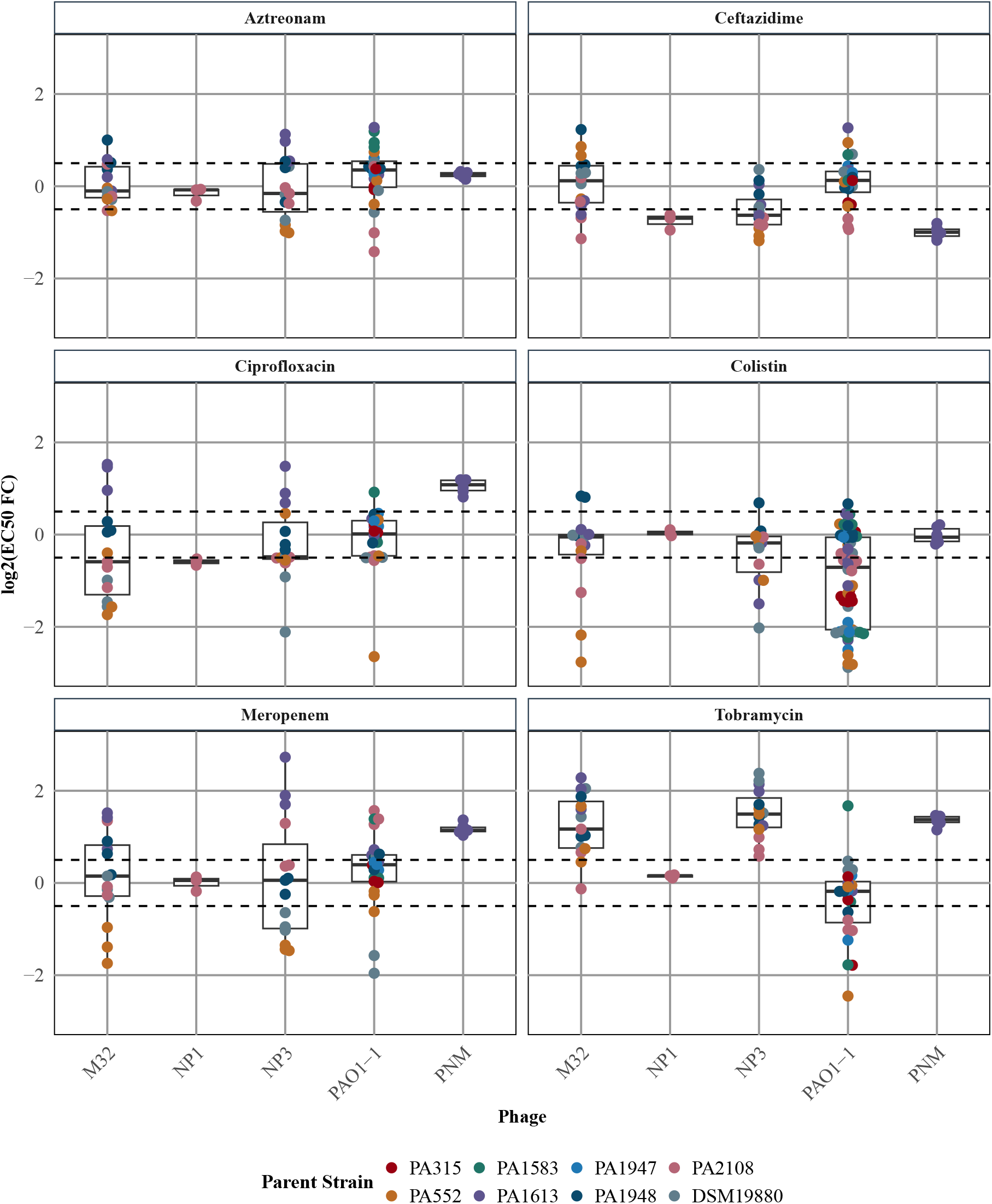
Collateral effects in single phage-resistant mutants derived from five different phages. The boxplot shows the median, range, and standard error of EC50_FC, log2_ across different antibiotics for each phage (M32, NP1, NP3, PAO1-1, and PNM) across multiple single phage-resistant mutants. After log_2_ transformation, values *≤ −*0.5 indicate collateral sensitivity (CS), and values *≥* 0.5 indicate collateral resistance (CR), while no collateral effects (CE) lie between -0.5 and 0.5. Each data point represents the mean of three biological replicates. The colour indicates the parental strain from which each mutant was derived, varying by phage due to differences in sensitivity (NP1 and PNM, n = 1; PAO1-1, n = 8; NP3 and M32, n = 5).

### 3.3 Collateral effects of combined antibiotic and phage resistance

To evaluate whether pre-existing antibiotic resistance influences collateral effects of phage resistance, we determined antibiotic sensitivity in antibiotic-resistant mutants before and after evolving additional PAO1-1 phage resistance. The antibiotic mutants generated had a decreased, but varying change in their antibiotic sensitivity compared to the parental strains: aztreonam (16-fold), ceftazidime (16 fold), ciprofloxacin (4-32 fold), colistin (1-4 fold), meropenem (64-256 fold), and tobramycin (4-16 fold) (Supplementary Table S2, Supplementary Figures S6, S7, S9, and S11). These antibiotic-resistant strains were evolved to develop phage resistance, after which we again evaluated antibiotic sensitivity in these strains. Our results demonstrate that in these double antibiotic- and phage-resistant mutants a significant decrease in EC50_FC, log2_ for colistin and tobramycin compared to single antibiotic-resistant mutants (p=0.011 and p=0.008, respectively, two-sample *t*-test; Supplementary Table S8; Figure 5). For aztreonam, ceftazidime, ciprofloxacin, and meropenem, the EC50_FC, log2_ remained elevated and comparable to that of the single antibiotic-resistant mutants.

**Figure 5.**
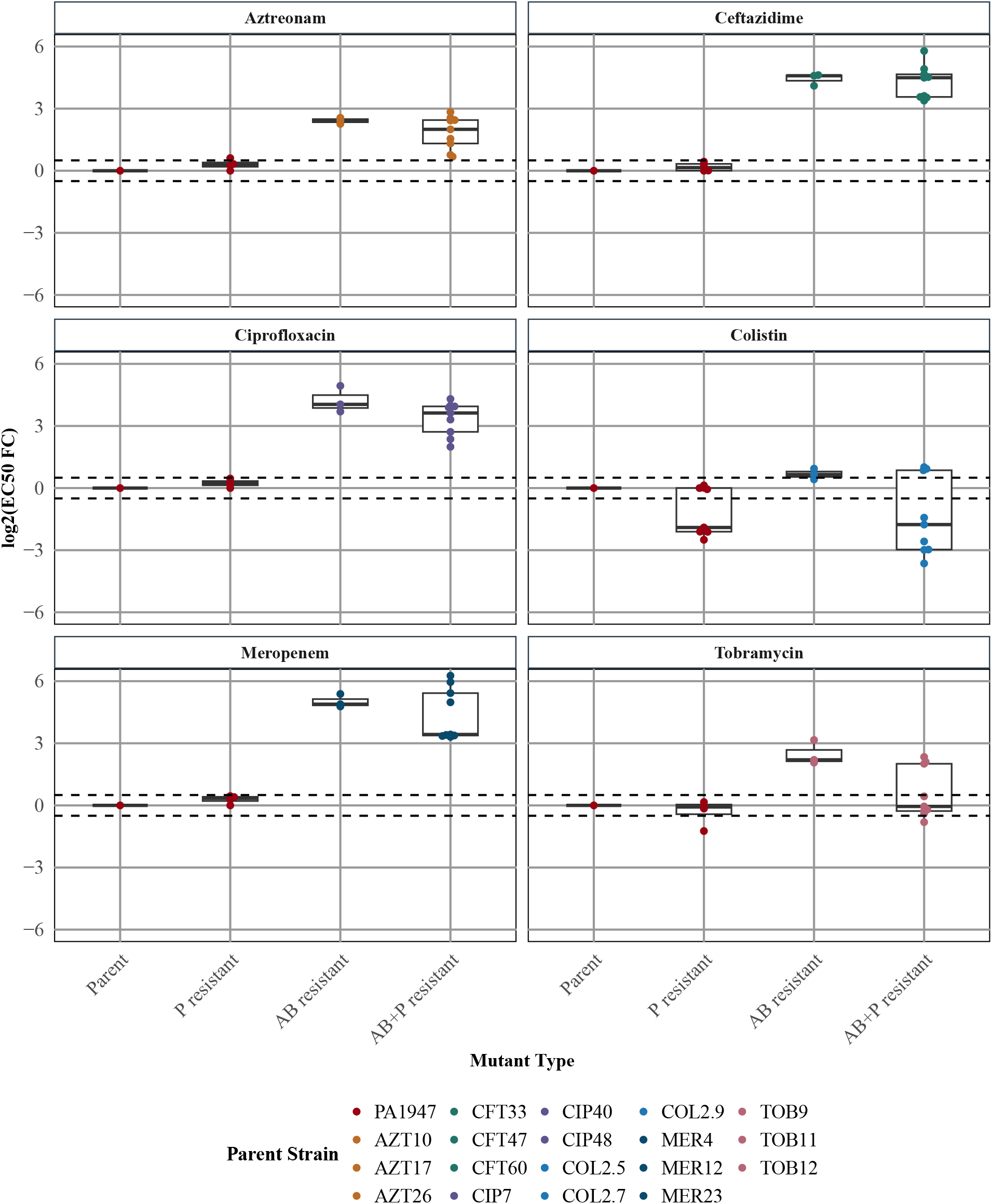
Change in antibiotic sensitivity in different resistant groups. The boxplot shows the median, range, and standard error of EC50_FC, log2_ across different antibiotics in strain PA1947 (Parent), the single PAO1-1 phage mutants (P resistant), the single antibiotic-resistant mutants (AB resistant), and the double antibiotic- and PAO1-1 phage resistant mutants (AB+P resistant). After log_2_ transformation, values *≤ −*0.5 indicate collateral sensitivity (CS), and values *≥* 0.5 indicate collateral resistance (CR), while no collateral effects (CE) lie between -0.5 and 0.5. Each data point represents the mean of three biological replicates.

### 3.4 Antibiotic-resistant mutants are associated with increased phage sensitivity

To investigate whether antibiotic resistance affects phage sensitivity, we quantified the EC50_FC, log2_ of phage PAO1-1 for the parental strain PA1947 and its three independent antibiotic-resistant mutants per antibiotic (Supplementary Figure S12). Across all single antibiotic mutants, the antibiotic-resistant mutants exhibited lower EC_50_ values to PAO1-1 phage compared to the parental strain, indicating collateral sensitivity to the phage (Figure 6). This effect was most pronounced for colistin-resistant mutants.

**Figure 6.**
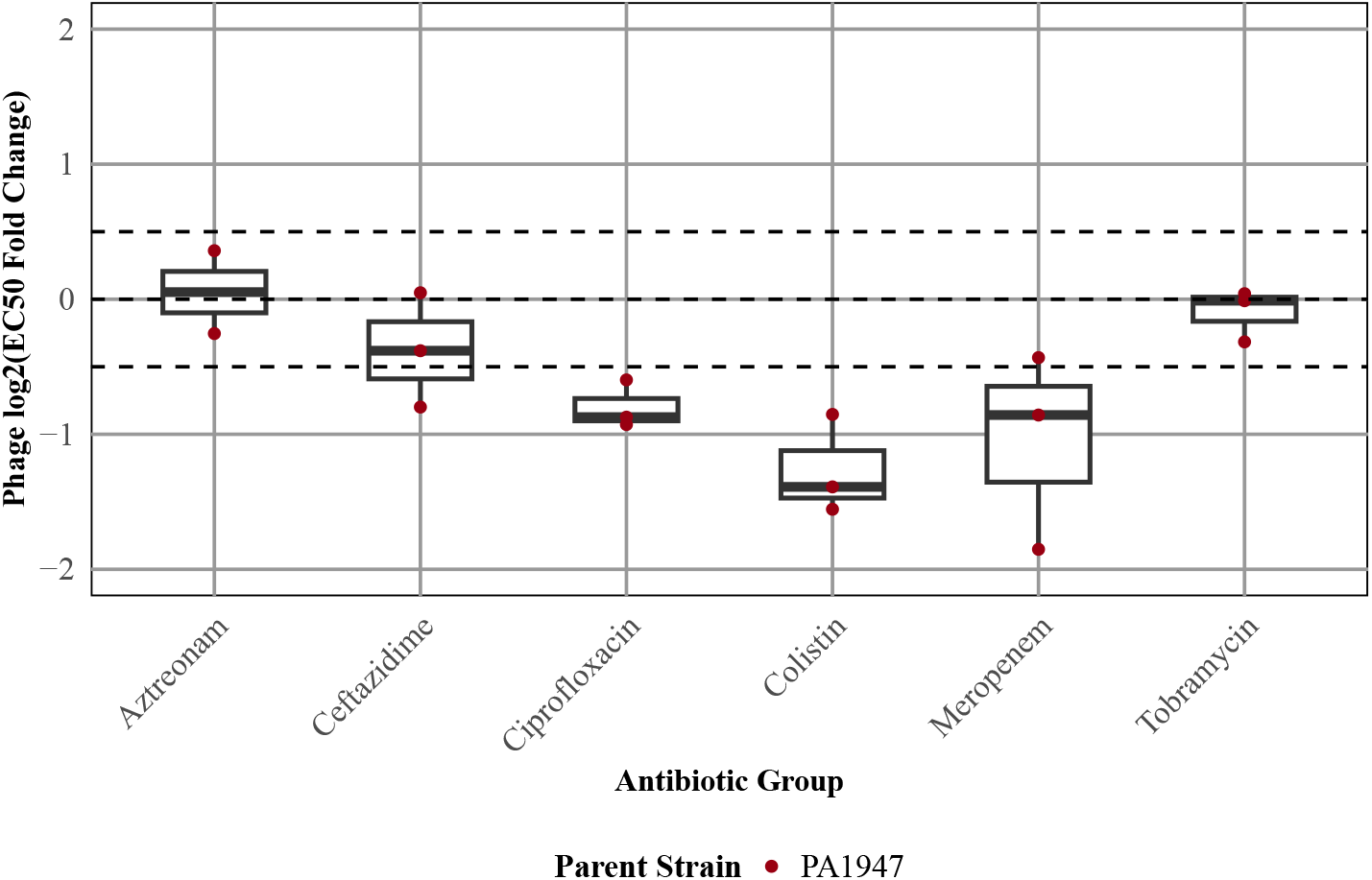
Collateral effects in single antibiotic-resistant mutants towards the PAO1-1 phage. The boxplot shows the median, range, and standard error of EC50_FC, log2_ across different single-antibiotic mutants per antibiotic group. The EC50_FC, log2_ of phage PAO1-1 for the parental strain PA1947 and its antibiotic-resistant mutants, showing the relative change in phage sensitivity compared to the parental strain. After log_2_ transformation of EC_50_, values *≤ −*0.5 indicate collateral sensitivity (CS), while values *≥* 0.5 indicate collateral resistance (CR), while no collateral effects (CE) lie between -0.5 and 0.5. Dashed horizontal lines indicate the thresholds for defining collateral sensitivity and -resistance. Each data point represents the mean of three biological replicates.

### 3.5 Fitness effects of phage resistance

Among the mutants tested, acquiring antibiotic resistance was associated with a lower fitness cost than acquiring phage resistance. Most single antibiotic-resistant mutants retained close to 100% of their maximum growth capacity, and growth rate was increased in several antibiotic-resistant mutants, particularly those resistant to tobramycin, meropenem, and ciprofloxacin (Supplementary Figures S14A and S14C). In contrast, nearly all single phage-resistant mutants exhibited a significant reduction in growth capacity, with some retaining only 30% of their maximum growth potential. Growth rate, however, was also increased in phage-resistant mutants: M32-, NP3-, and PAO1-1-resistant mutants remained mostly between 50-100%, with a subset reaching 200-400%, while PNM-resistant mutants showed a larger overall increase, ranging between 150-400% (Supplementary Figures S14B and S14D).

### 3.6 Gene-level associations with collateral effects

In the PAO1-1 phage-resistant mutants generated, whole genome sequencing demonstrated mutation counts which varied considerably across parental strains, ranging from 7 (PA1947) to 122 (PA2108) mutations per strain, with most strains falling between 18 and 26 mutations (Supplementary Table S3). Despite sharing the same parental background, the proportion of mutations shared across all three PAO1-1-resistant mutants within a parental strain was low, ranging from zero to five shared genes for most strains and approximately 20% for PA2108.

Between different parental strains, mutations in several recurring specific genes across up to three different parental strains were found (Figure 7). The genes found mutated across different parental strains were *hemH*, *metH*, *suhB*, *tse1*, *ccoN*, *dnaX*, *pqqA*, *rrs*, *valS*, and *wzy*. These genes are associated with DNA replication and transcription (*suhB*, *dnaX*, *rrs*, *valS*), virulence and regulatory systems (*metH*, *tse1*), metabolism and biosynthesis (*hemH*, *pqqA*), the cell envelope (*wzy*), and other processes (*ccoN* ; Gene Ontology Biological Process annotation (UniProt and QuickGO for *Pseudomonas aeruginosa* PAO1, taxon ID 208964; Supplementary Table S9).

**Figure 7.**
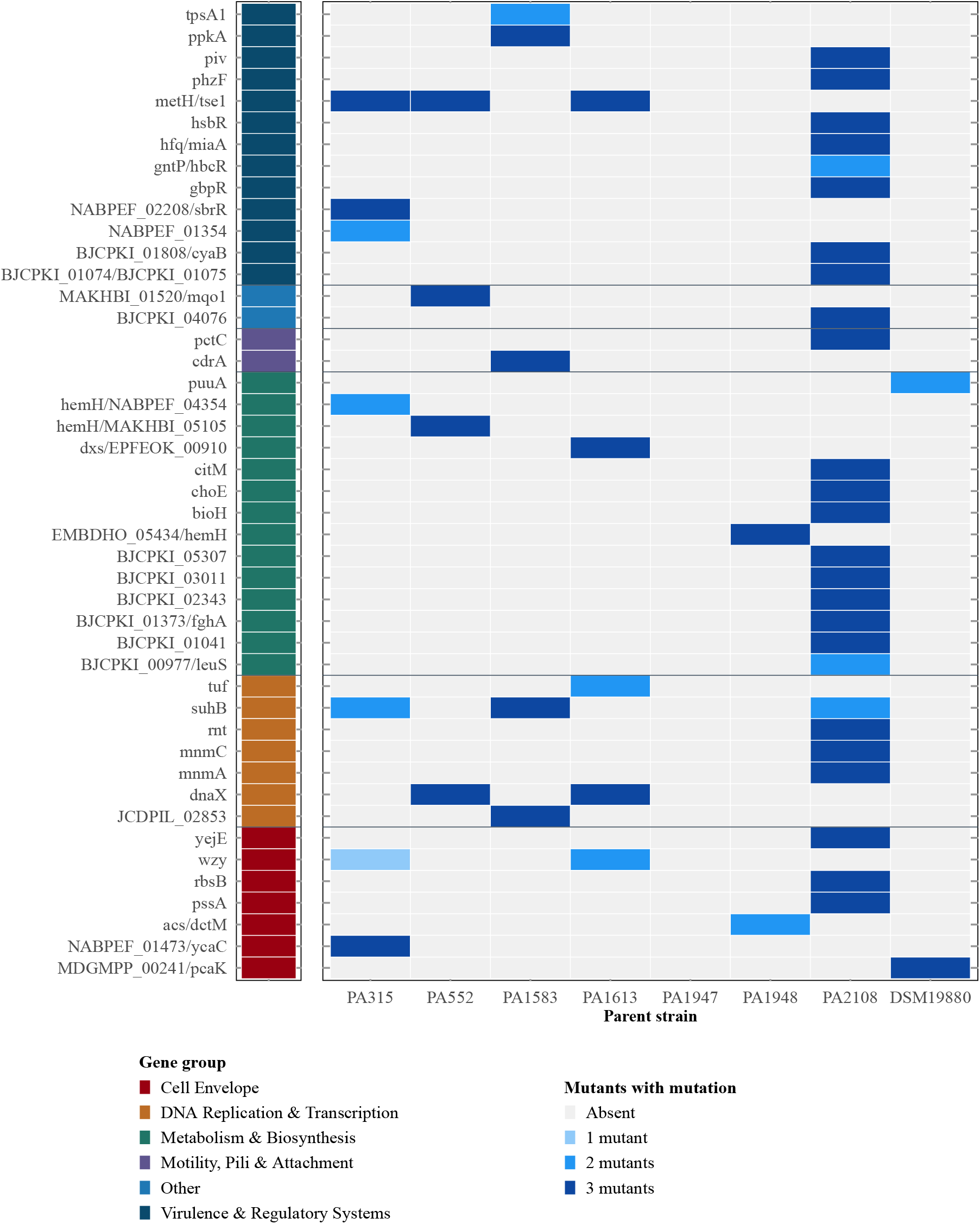
Distribution of genetic mutations across PAO1-1 phage-resistant mutants. Genes are grouped by functional category based on Gene Ontology annotation (Supplementary Table S9). For each parental strain, three independent phage-resistant mutants were sequenced. Blue shading indicates how many of the three mutants carry a mutation in a given gene, ranging from absent (light grey) to present in all three mutants (dark blue).

To identify genes associated with potential collateral effects alterations in antibiotic sensitivity, we regressed the incidence of mutated genes to the antibiotic-specific EC50_FC, log2_ as a continuous measure of collateral resistance or -sensitivity using PLS regression. The percentage for which gene identity could explain variation in the EC50_FC, log2_ differed among antibiotics (38-79%; Supplementary Figure S20). Across all antibiotics, a single PLS component was sufficient to capture the gene-associated variation in the EC50_FC, log2_ . The PLS regression coefficients and VIP (*>*1) scores identified genes contributing to either increased (CR) or decreased (CS) sensitivity per antibiotic (Figure 8).

**Figure 8.**
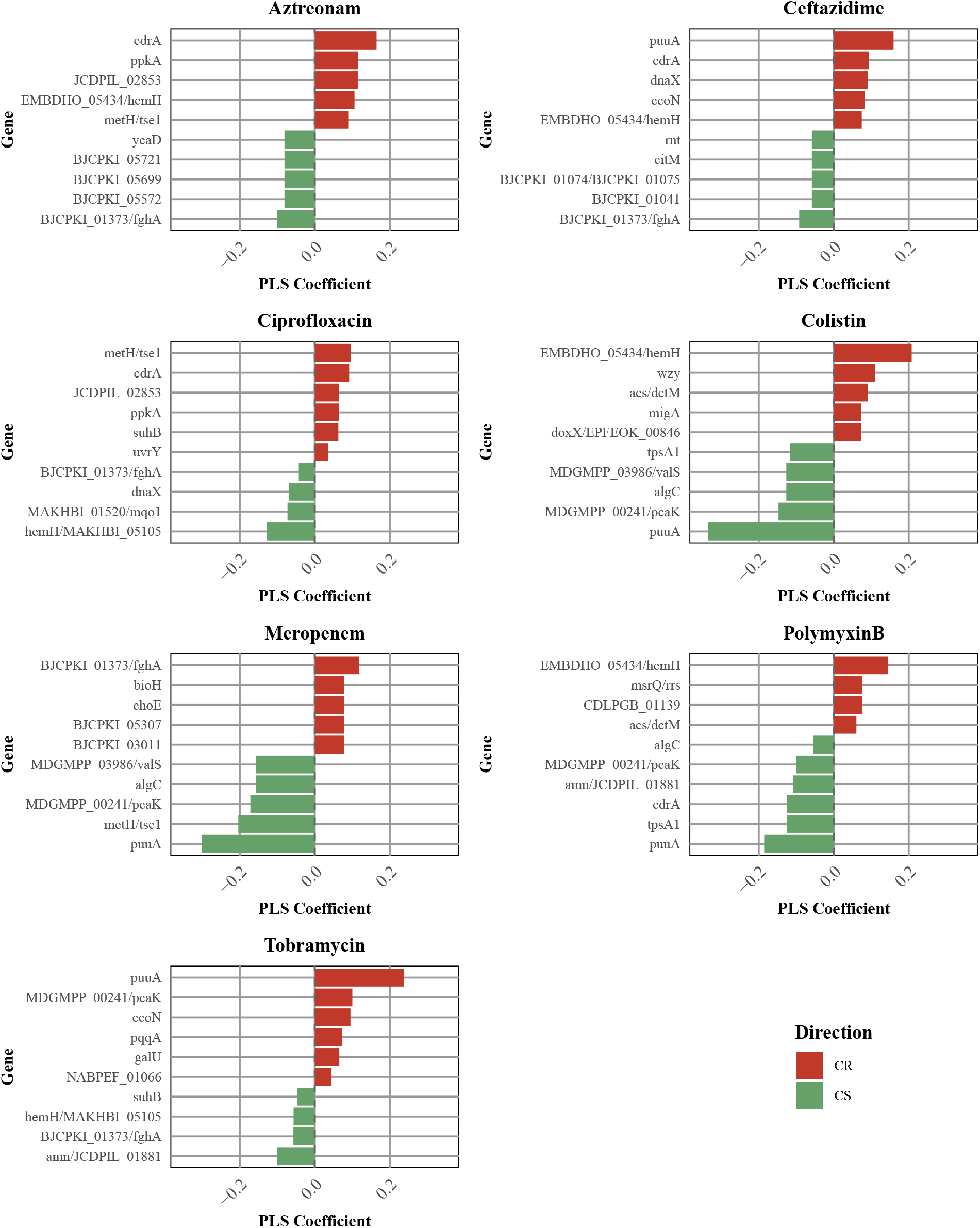
Partial Least Squares (PLS) regression coefficients per antibiotic. Each panel shows the top 5 genes contributing to collateral resistance (CR, red) and sensitivity (CS, green) based on VIP*>*1. The x-axis represents the PLS coefficient magnitude and direction. The y-axis represents the identified genes.

Mutations in *cdrA* were the highest-ranked association with CR for both aztreonam and ceftazidime, suggesting a shared resistance mechanism across these two *β*-lactams. Mutations in *puuA* showed the most pronounced CS association for colistin, meropenem, and polymyxin B, yet conversely ranked as CR-related for tobramycin and ceftazidime, indicating an antibiotic class-dependent effect on sensitivity. Mutations in *hemH* were the most-pronounced CR-related association for both colistin and polymyxin B, and also highly ranked in aztreonam and ceftazidime, while displaying a CS association for ciprofloxacin and tobramycin. Mutations in *algC* were associated with CS for colistin, polymyxin B, and meropenem. A nonsense mutation at Y438 introduces a premature stop codon, likely resulting in a truncated, non-functional protein.

## 4 Discussion

The current study provides a systematic insight into the occurrence of CEs across PA strains, phages, and antibiotics, and potential genetic associations related to such collateral effects.

Phage-resistant mutants against phage PAO1-1 exhibited relatively consistent CS effects for colistin and polymyxin B. Despite the use of different phages, previous studies have reported similar increased in colistin- and polymyxin B sensitivities.^12,14,22^ Our findings broadly agree with Wannasrichan *et al.* (2022), who induced phage resistance in a PAO1 strain with the use of another *Pbunavirus* phage JJ01.^14^ Both theses studies also reported an increased colistin sensitivity (1- to 3-fold decreases in MIC and EC50_FC, log2_ , respectively), minor decreased sensitivity towards the *β*-lactams aztreonam, ceftazidime, and meropenem (both studies not significant) and no shifts for ciprofloxacin (both studies not significant). The exception was tobramycin, where Wannasrichan *et al.* (2022) reported a 1-fold MIC increase in most mutants, while we observed largely stable sensitivity with occasional 1 to 2.5-fold EC50_FC, log2_ decreases.

We compared phage-resistant mutants derived from five different phages and found that sensitivity to ceftazidime and tobramycin depended on which specific phage resistance was evolved, whereas collateral effects for the remaining antibiotics was similar regardless of phage identity. Variability in collateral effects among different phages likely reflects stronger selective pressure on specific phage resistance mechanisms at phage binding sites.^23^ Comparisons with other studies are challenging, as most focus on a single phage when investigating phage resistance and antibiotic sensitivity, or they use phage cocktails rather than individual phages. In *Acinetobacter baumannii*, two different phages exhibited varying levels of sensitivity to antibiotics such as ciprofloxacin and meropenem, highlighting phage-specific collateral effects.^24^

Since phage therapy is often considered for antibiotic resistant infections, we assessed whether antibiotic resistance affects the collateral effects of subsequent phage resistance by evolving PAO1-1 phage resistance in antibiotic-resistant mutants. These double antibiotic- and phage-resistant mutants displayed strong CS to colistin and tobramycin, with EC50_FC, log2_ values dropping below those of the parental strain for colistin, and returning to parental-strain levels for tobramycin. Since the MICs of the parental strain fall below the EUCAST clinical breakpoint for both antibiotics, this suggests a return to clinical susceptibility, while resistance to the *β*-lactams (aztreonam, ceftazidime, and meropenem) was largely maintained. The restoration of clinical susceptibility through phage resistance has been reported in other pathogens. For example, in colistin-resistant *Klebsiella pneumoniae*, phage resistance reduced MIC values back to therapeutically effective levels^25^, consistent with the colistin and tobramycin patterns observed here.

Among the antibiotics mutants, the largest increase in PAO1-1 phage sensitivity was observed in colistin-resistant mutants, suggesting a shared mechanistic basis between colistin resistance and PAO1-1 phage sensitivity. This is consistent with previous findings for *Klebsiella pneumoniae*, where colistin-resistant strains showed greater phage sensitivity than their non-resistant counterparts^25^, pointing to a broader pattern linking colistin resistance to phage sensitivity. Together with the finding above that phage resistance increases colistin sensitivity, this points to a reciprocal collateral relationship between colistin- and phage resistance that is of particular clinical interest.

Phage-resistant mutants often exhibit fitness costs, such as reduced maximal cell densities^26^, slower growth and decreased biofilm formation^14^, and impaired twitching motility.^27^ In line with previous results, we observed a decrease in maximum growth capacity in phage-resistant mutants consistent over five different phages, while the maximum growth capacity for antibiotic mutants remains relatively similar. The growth rate for phage-resistant mutants not only decreased, but also showed increased growth rates. This has been observed previously in phage-resistant bacteria, where resistance mutations shifted population growth dynamics resulting in higher densities^28^, and may further reflect compensatory mutations that partially restore fitness following the initial resistance cost.^29^ The magnitude of this cost appears to depend on the type of resistance mutation acquired, with LPS- and capsule biosynthesis mutants showing significantly greater growth reductions than membrane protein mutants.^30^

We performed exploratory whole genome sequencing to evaluate potential associations between observed mutated genes and CS- or CR phenotypes. Notably, these analyses should be interpreted with appropriate caution as the number of evaluated strains is insufficient to robustly link individual mutations to CS or CR phenotypes. Larger, more targeted mutant panels will be needed to validate these associations. Having established this, we focused on mutations in three genes that emerged most consistently as top contributors in the partial least squares (PLS) analysis linking mutated genes to antibiotic CS effects in phage-resistant mutants.

Mutations in *algC* were associated with CS for colistin, meropenem and polymyxin B. Phosphomannomutase algC is involved in the production of UDP-glucose, which is needed for LPS core biosynthesis.^13,31^ Notably, CS was observed across mechanistically distinct antibiotic classes, monobactams and polymyxins, both of which rely on outer membrane integrity or LPS as their primary target, underscoring the broad impact of *algC* loss-of-function. Mutations in *algC* are frequently reported in phage-resistant mutants, consistent with its role as a common phage receptor component.^31–34^

Mutations in the *cdrA* gene were associated with CR to aztreonam, ceftazidime, and ciprofloxacin, and associated with CS to polymyxin B. *CdrA* is a fibrillar adhesin that cross-links the Psl polysaccharide matrix and promotes cell-cell aggregation and biofilm maturation.^35,36^ Loss-of-function mutations in *cdrA* likely disrupt this matrix, and the resulting effect on antibiotic sensitivity appears to depend on how each antibiotic reaches its target. For the *β*-lactams aztreonam and ceftazidime, as well as for ciprofloxacin, a disorganized or non-functional matrix may restructure the outer membrane in a way that limits antibiotic penetration, producing CR. Polymyxins act directly on the outer membrane and LPS, which may disrupt the membrane integrity in a way that increases antibiotic access, consistent with the CS observed for polymyxin B.

Mutations in *puuA* were CS-associated for colistin, meropenem, and polymyxin B, and CR-associated for ceftazidime and tobramycin. *γ*-glutamylputrescine synthetase PuuA catalyzes the first step of putrescine utilization, and excess putrescine is toxic and inhibits growth and protein synthesis.^37,38^ Mutations disrupting *puuA* function likely impair putrescine catabolism, leading to elevated intracellular putrescine levels.^39,40^ This is seemingly paradoxical for cationic antimicrobials such as colistin and polymyxin B, since exogenous polyamines are known to increase outer membrane positive charge and reduce their efficacy.^41,42^ However, the metabolic burden of impaired putrescine turnover likely outweighs any membrane charge effect, sensitizing the cell overall. Mutations in *puuA* might also explain the sharp decrease in maximal carrying capacity, given its role in metabolism.

We quantified antibiotic- and phage sensitivity through estimation of the EC_50_, derived from full dose-response growth curves for total bacterial growth as quantified using the AUC. This approach allowed sensitivity to be determined in a consistent manner for both antibiotics and phages, alongside other growth parameters such as maximal carrying capacity and growth rate, and enabled high-throughput assessment across many mutant-antibiotic-phage combinations. The methodology quantifies drug effects on overall bacterial burden, and is thereby a more robust measure of drug effects compared to e.g., MIC-type single-type point measurements. This methodological decision does however mean our results are not directly translatable to MICs equivalents, although it is likely proportional relationships between e.g. overall bacterial suppression and MIC are likely to apply.

The current study observed variable magnitudes of CS effects due to phage resistance. Whether these observed CS effects are sufficient to lead to a therapeutic benefit are related to the therapeutic index of the antibiotic for which CS is observed. When considering antibiotics with a narrow therapeutic index (i.e., antibiotics with a higher toxicity risk), the occurrence of CS would effectively increase this therapeutic windows by lowering the concentration required to eradicate a pathogen, and thereby increasing the likelihood of therapeutic success. In addition, when considering a strain at the boundary of sensitive and clinically resistant, the occurrence of CS could re-sensitize such a pathogen to treatment. In this context, our finding of e.g., the observed for colistin is of clinical relevance, as colistin is a toxic antibiotic with a very narrow therapeutic index and where a modest concentration reduction could shift the balance between efficacy and toxicity.^43^ Because collateral effects were largely antibiotic-specific, phage exposure could be used to steer bacterial evolution towards a CS-associated genotype for a specific antibiotic which can be incorporated in the treatment protocol.^11^ Similarly, in case of CR that was observed for several antibiotic-phage combinations, explicit consideration in the context of phage-antibiotic combination therapy is warranted, as CR could potentially compromise treatment efficacy. Clinical application of CS-based antibiotic-phage combination therapies is conditional on the robustness of observed CS effects across strains, but which is also phage-specific. Our study demonstrates that whilst CS effects have been observed for different antibiotics, variability is high, suggesting there are limitations regarding the broader potential for CS across a wider range of antibiotics.

In conclusion, this study systematically characterized collateral effects across eight PA strains, seven antibiotics, and five phages, testing single antibiotic- and phage-resistant mutants as well as double mutants for changes in antibiotic sensitivity. CS was observed most frequently for colistin and polymyxin B, and colistin resistance was likewise associated with increased phage sensitivity. Colistin therefore stands out as a promising lead for the rational design of future phage-antibiotic combination therapies against multidrug-resistant *Pseudomonas aeruginosa*.

## 5 Author contributions

LRB, TG and JGCVH conceived the project. LRB, KN and JGCVH designed the experiments. LRB and KN performed experimental work. LRB and KN performed data analysis. LRB and LBZ performed statistical analysis. LRB wrote the initial draft. LRB, KN, LBZ, TG and JGCVH revised the manuscript.

## 6 Acknowledgements

We are grateful to Shireen E. Baars for creating the graphical illustrations of the experimental setup and the data analysis. We also thank Apostolos Liakopoulos for his support with the interpretation of the sequencing results.

## 8 Transparency declarations

Nothing to declare.

## 9 Data availability statement

The raw dataset of this study is available upon request.

## 11 Supplementary

### 11.1 Antibiotic information

**Supplementary Table S1.** Details on antibiotic dilutions, including solvent compatibility, storage conditions, and manufacturer information.

| Antibiotic | Solvent | Storage Condition | Manufacturer |
| --- | --- | --- | --- |
| Aztreonam | DMSO | −20 °C | Acros Organics |
| Ceftazidime | DMSO | −80 °C | Acros Organics |
| Ciprofloxacin | 0.1 M HCl | −20 °C | Chem-Impex |
| Colistin | Water | −80 °C | Cayman Chemical |
| Meropenem | DMSO | −20 °C | ChemCruz |
| Polymyxin B | Water | −80 °C | Thermo Scientific |
| Tobramycin | Water | −20 °C | Chem-Impex |

### 11.2 Overview of single antibiotic-resistant mutants

**Supplementary Table S2.** Minimal inhibitory concentration (MIC; mg/L) and fold change (FC) of antibiotic-resistant mutants compared to their individual parental strain(PA1947).

| Antibiotic | Strain | MIC (mg/L) | FC | Antibiotic | Strain | MIC (mg/L) | FC |
| --- | --- | --- | --- | --- | --- | --- | --- |
| Aztreonam<br>EUCAST breakpoint: 16 | PA1947 | 2 | – | Colistin<br>EUCAST breakpoint: 4 | PA1947 | 2 | – |
|  | AZT10 | 32 | x16 |  | COL2.5 | 4–8 | x2–4 |
|  | AZT17 | 32 | x16 |  | COL2.7 | 4–8 | x2–4 |
|  | AZT26 | 32 | x16 |  | COL2.9 | 2–8 | x1–4 |
| Ceftazidime<br>EUCAST breakpoint: 8 | PA1947 | 1 | – | Meropenem<br>EUCAST breakpoint: 8 | PA1947 | 0.125 | – |
|  | CFT33 | 16 | x16 |  | MER4 | 8–16 | x64–128 |
|  | CFT47 | 16 | x16 |  | MER12 | 8–16 | x64–128 |
|  | CFT60 | 16 | x16 |  | MER23 | 8–32 | x64–256 |
| Ciprofloxacin<br>EUCAST breakpoint: 0.5 | PA1947 | 0.25 | – | Tobramycin<br>EUCAST breakpoint: 2 | PA1947 | 0.25 | – |
|  | CIP7 | 1–8 | x4–32 |  | TOB9 | 2–4 | x8–16 |
|  | CIP40 | 1–4 | x4–16 |  | TOB11 | 1–4 | x4–16 |
|  | CIP48 | 1–8 | x4–32 |  | TOB12 | 1–2 | x4–8 |

### 11.3 Antibiotic dose–response curves in single phage-resistant mutants

**Supplementary Figure S1.**
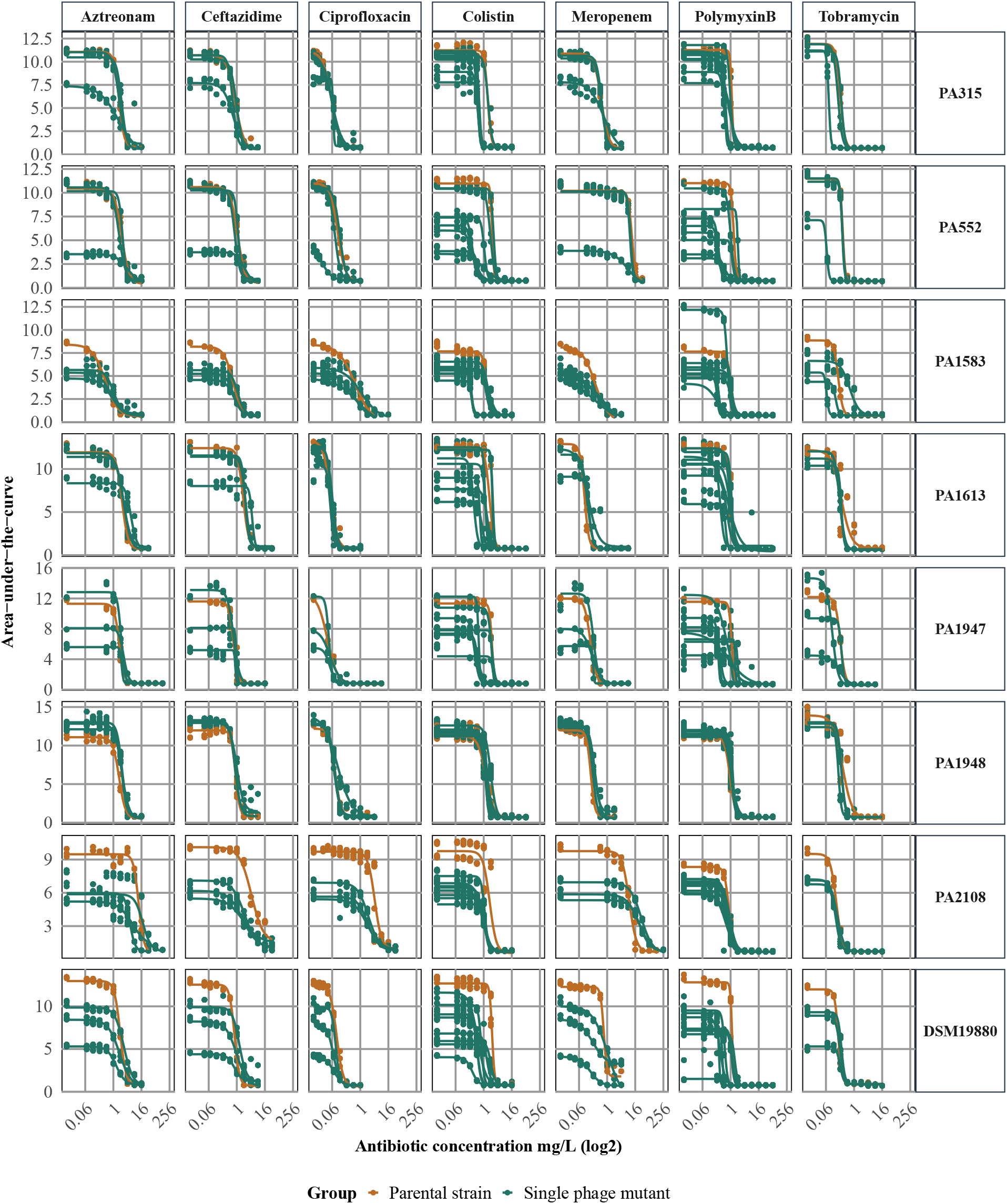
Dose–response curves (AUC vs. antibiotic concentration, mg/L; log_2_ scale) for PAO1-1 single phage-resistant mutants (green). For each antibiotic, three independent mutants derived from the same parental strain are shown (orange), except for colistin and polymyxin B, for which eight mutants were measured. The original parental strain is displayed on the right y-axis. Curves were fitted using *drm* in R from three biological replicates per concentration.

**Supplementary Figure S2.**
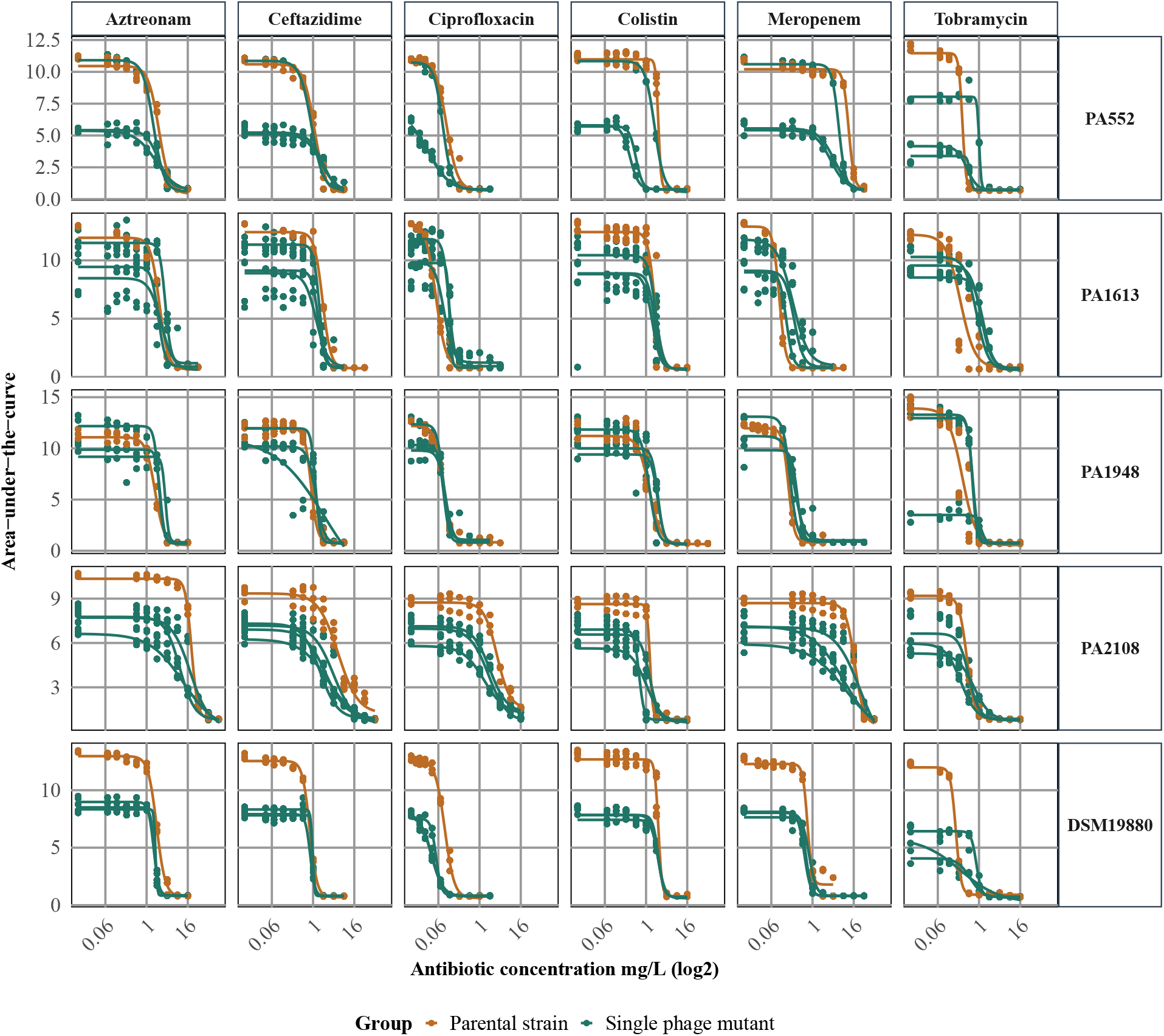
Dose–response curves (AUC vs. antibiotic concentration, mg/L; log_2_ scale) for M32 single phage-resistant mutants (green). For each antibiotic, three independent mutants derived from the same parental strain are shown (orange). The original parental strain is displayed on the right y-axis. Curves were fitted using *drm* in R from three biological replicates per concentration.

**Supplementary Figure S3.**
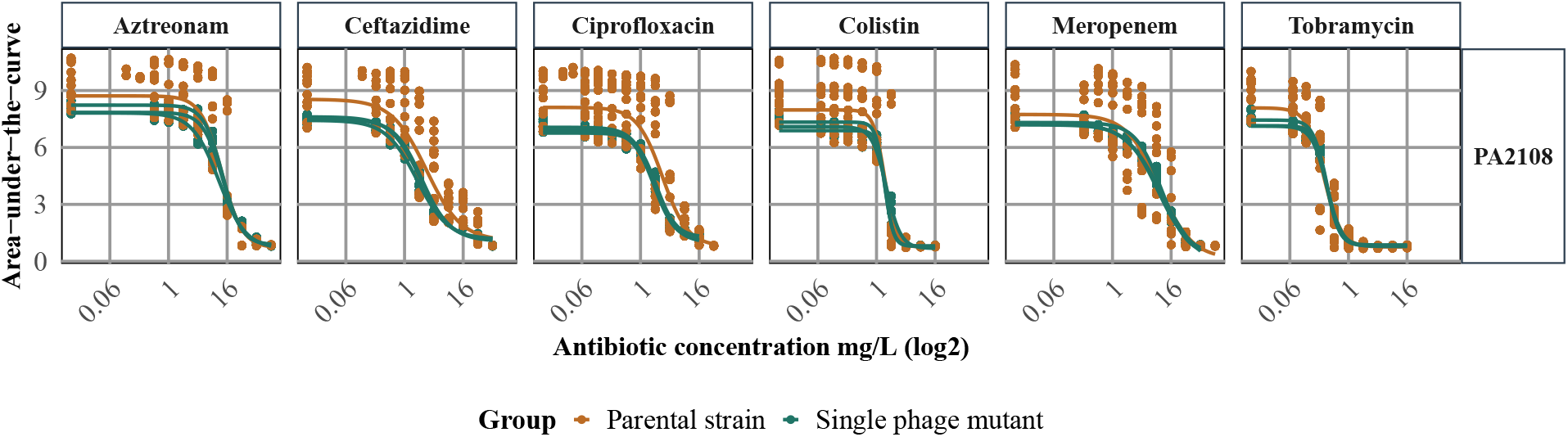
Dose–response curves (AUC vs. antibiotic concentration, mg/L; log_2_ scale) for NP1 single phage-resistant mutants (green). For each antibiotic, three independent mutants derived from the same parental strain are shown (orange). The original parental strain is displayed on the right y-axis. Curves were fitted using *drm* in R from three biological replicates per concentration.

**Supplementary Figure S4.**
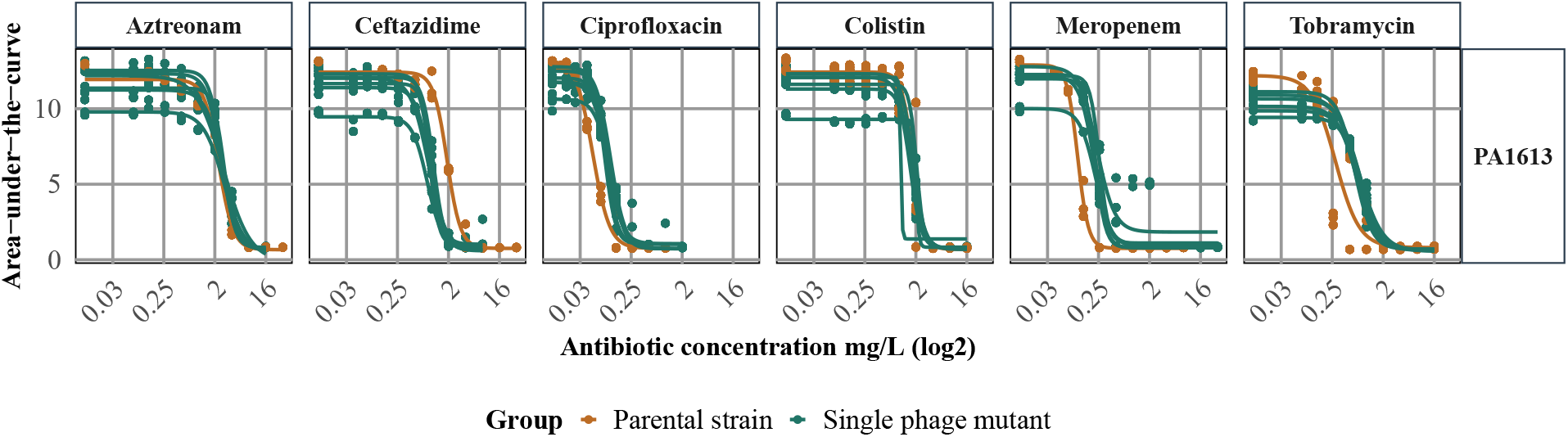
Dose–response curves (AUC vs. antibiotic concentration, mg/L; log_2_ scale) for PNM single phage-resistant mutants (green). For each antibiotic, six independent mutants derived from the same parental strain are shown (orange). The original parental strain is displayed on the right y-axis. Curves were fitted using *drm* in R from three biological replicates per concentration.

**Supplementary Figure S5.**
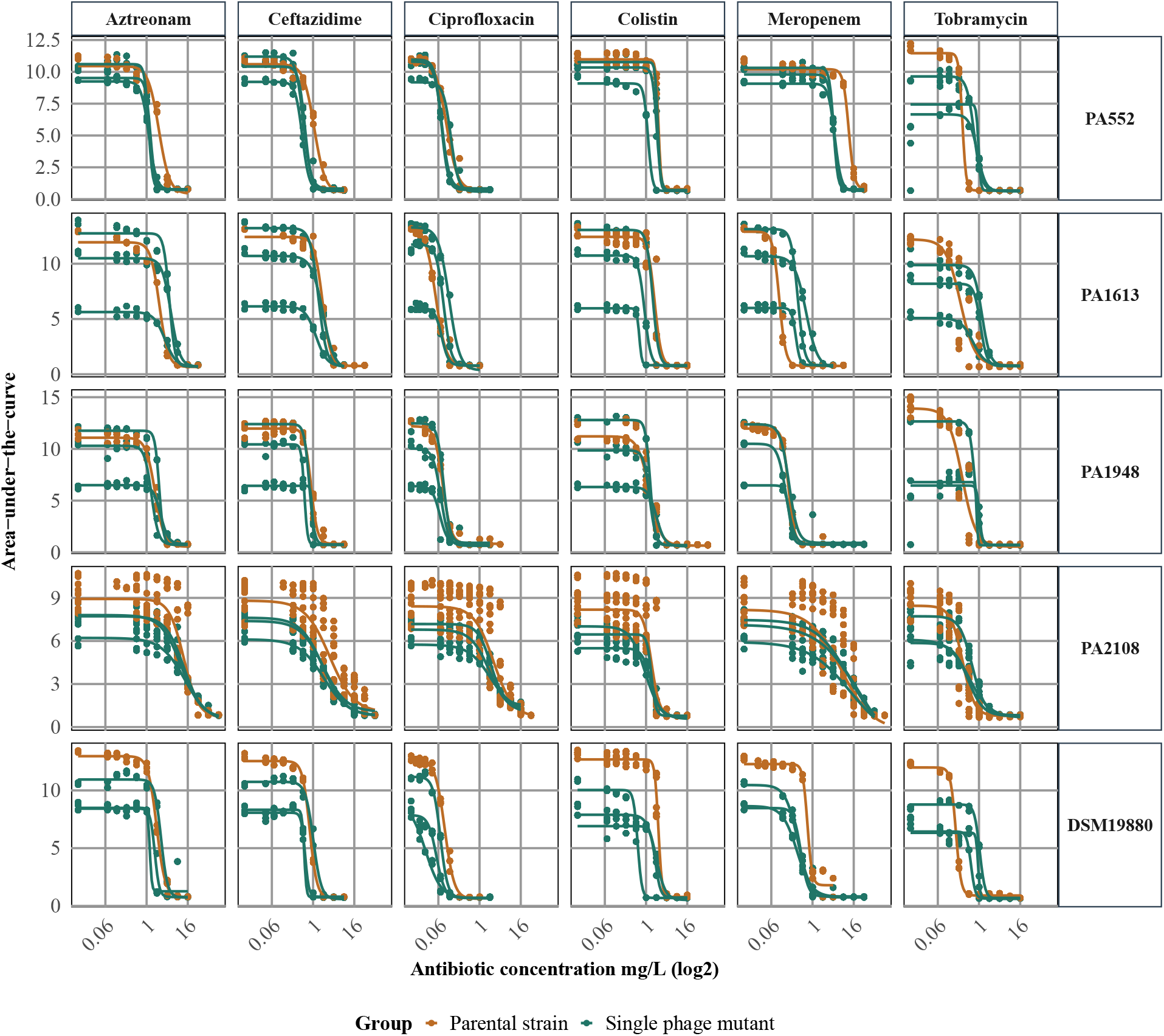
Dose–response curves (AUC vs. antibiotic concentration, mg/L; log_2_ scale) for NP3 single phage-resistant mutants (green). For each antibiotic, three independent mutants derived from the same parental strain are shown (orange). The original parental strain is displayed on the right y-axis. Curves were fitted using *drm* in R from three biological replicates per concentration.

### 11.4 Antibiotic dose–response curves in single antibiotic-resistant mutants

**Supplementary Figure S6.**
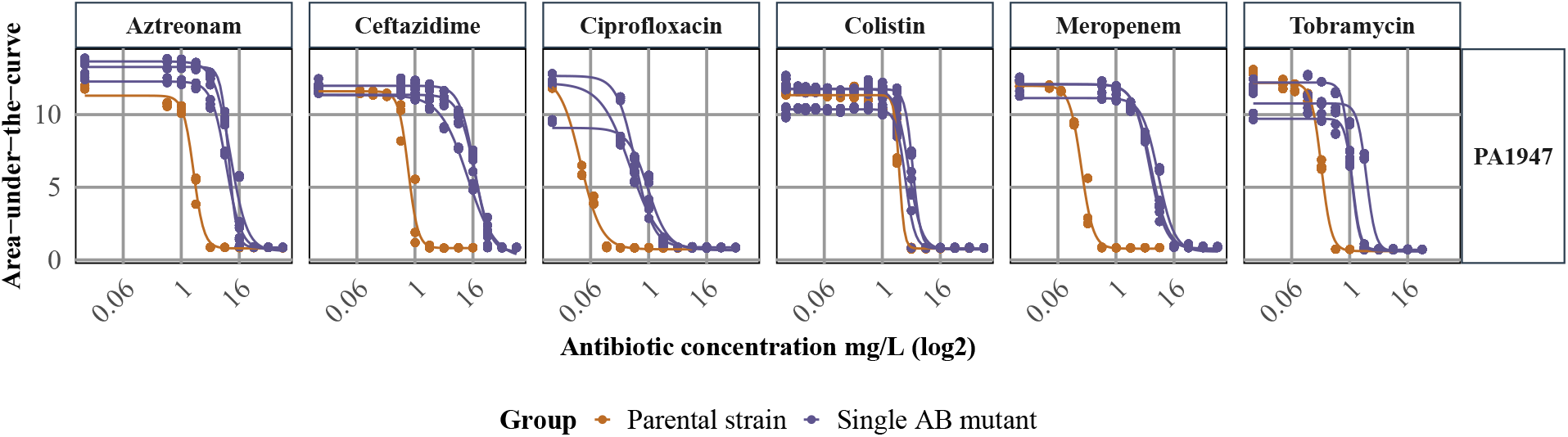
Dose–response curves (AUC vs. antibiotic concentration, mg/L; log_2_ scale) for single antibiotic-resistant mutants (purple) and three independent mutants derived from the same parental strain (orange). The original parental strain is displayed on the right y-axis. Curves were fitted using *drm* in R from three biological replicates per concentration.

### 11.5 Antibiotic dose–response curves in double antibiotic- and phage-resistant mutants

**Supplementary Figure S7.**
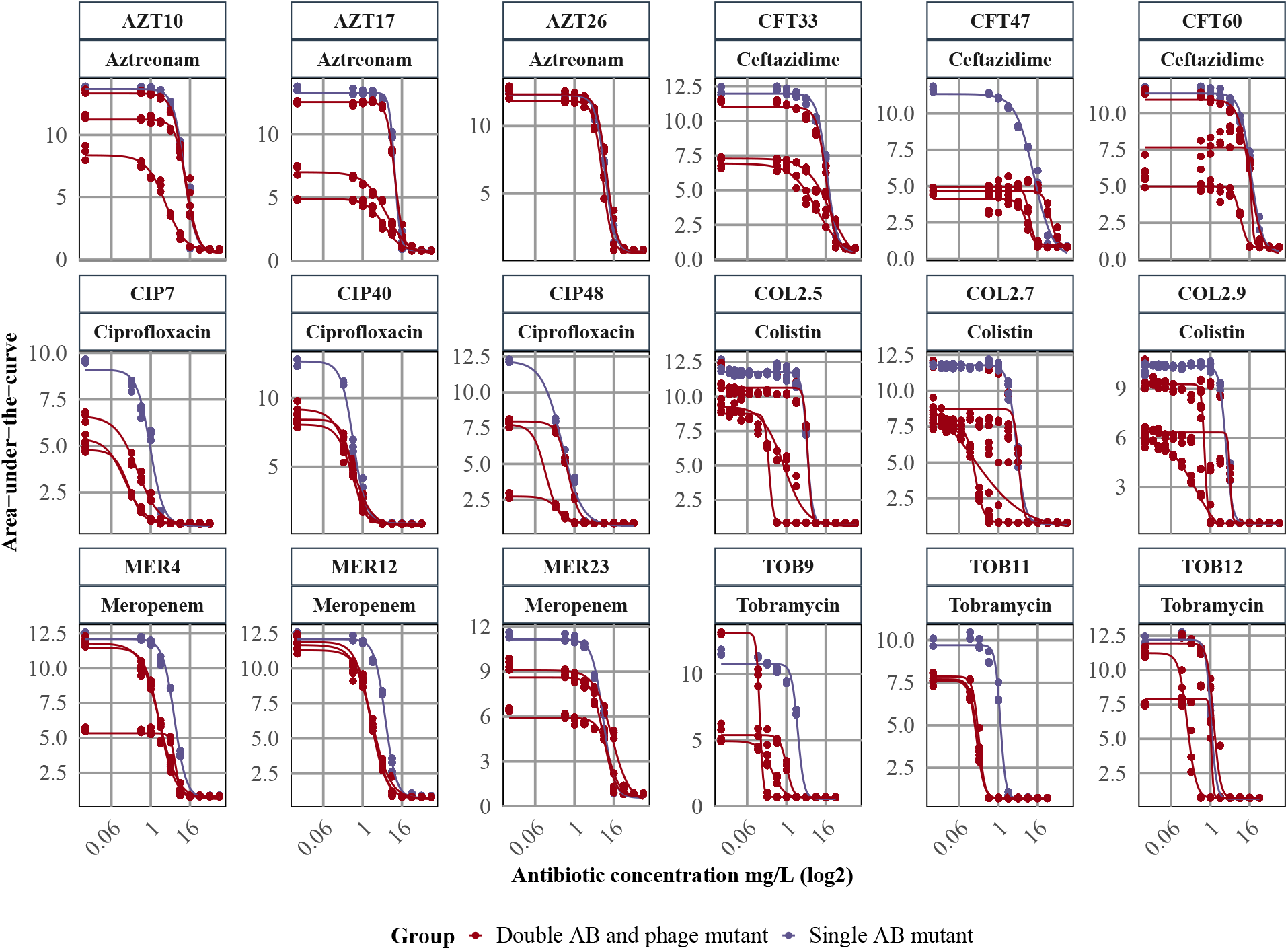
Dose–response curves (AUC vs. antibiotic concentration, mg/L; log_2_ scale) for double antibiotic- and phage-resistant mutants (red). Single antibiotic-resistant mutants are shown in purple as the parental reference, with individual antibiotic sensitivity levels measured. Curves were fitted using *drm* in R from three biological replicates per concentration.

### 11.6 Phage dose–response curves in single- and double phage-resistant mutants

**Supplementary Figure S8.**
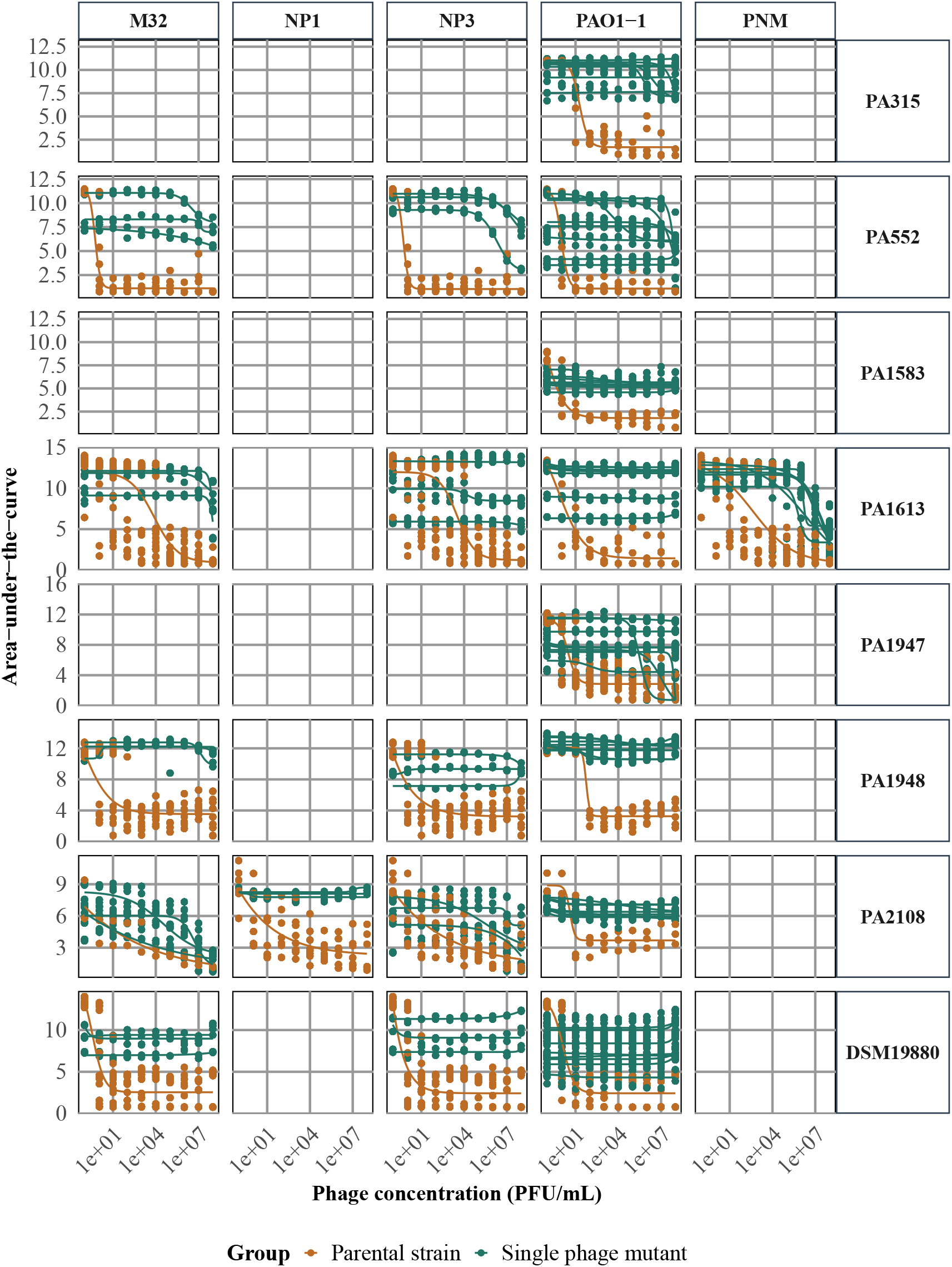
Dose–response curves (AUC vs. phage concentration, PFU/mL; log_10_ scale) for single phage-resistant mutants (green) for phages M32, NP1, NP3, PAO1-1, and PNM. For each case, three/eight independent mutants derived from the same parental strain are shown (orange) and tested against the phage to which they evolved resistance. The original parental strain is shown at the top. Curves were fitted using *drm* in R from three biological replicates per concentration.

**Supplementary Figure S9.**
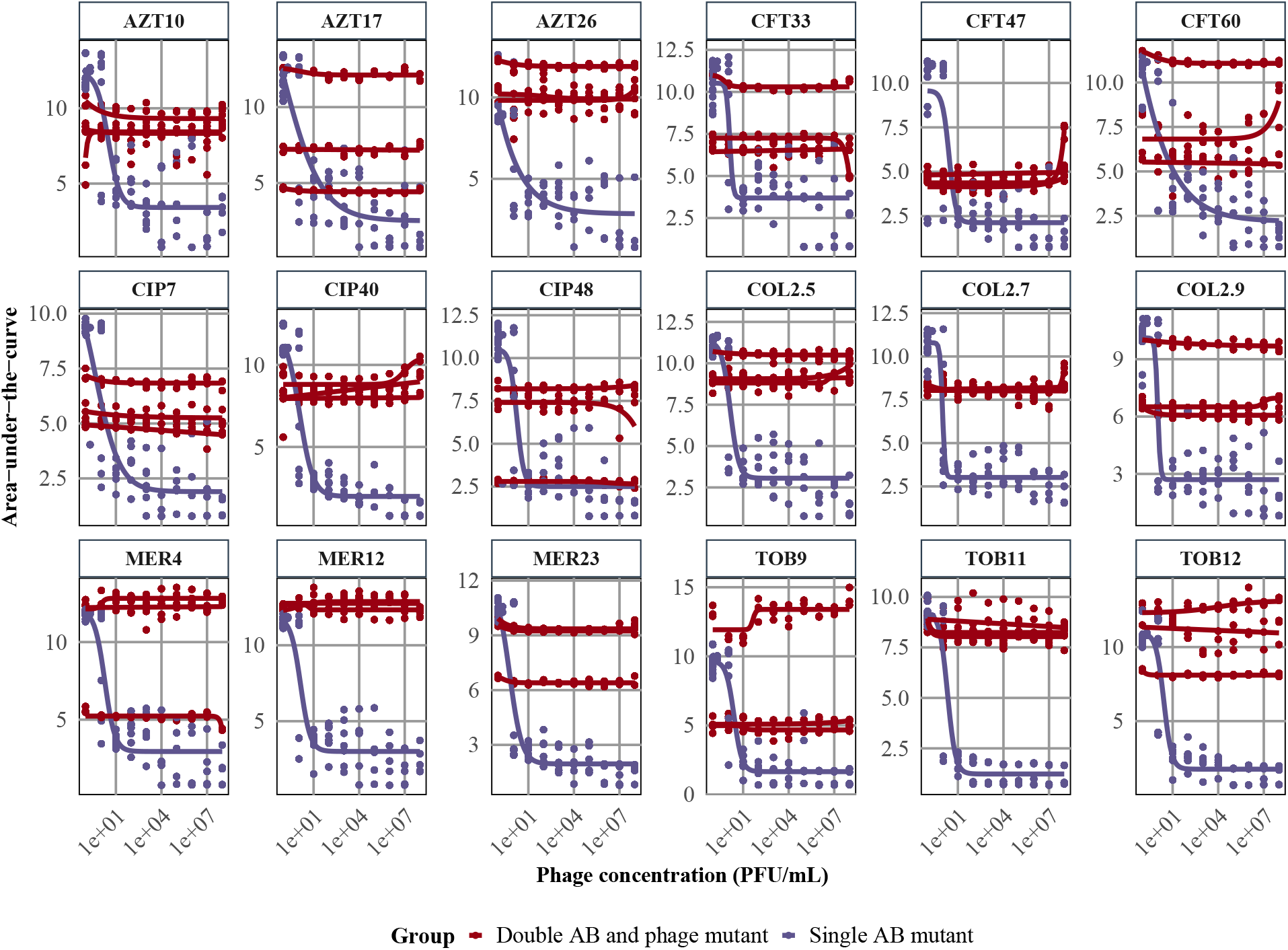
Dose–response curves (AUC vs. phage concentration, PFU/mL; log_10_ scale) for single antibiotic-resistant strains (purple) and their antibiotic- and PAO1-1 phage-resistant mutants (red). For each condition, three independent mutants derived from the same parental strain were measured for PAO1-1 sensitivity to assess phage-resistance levels. The parental strain is shown at the top. Curves were fitted using *drm* in R from three biological replicates per concentration.

### 11.7 Overview of collateral effects in phage-resistant mutants

**Supplementary Figure S10.**
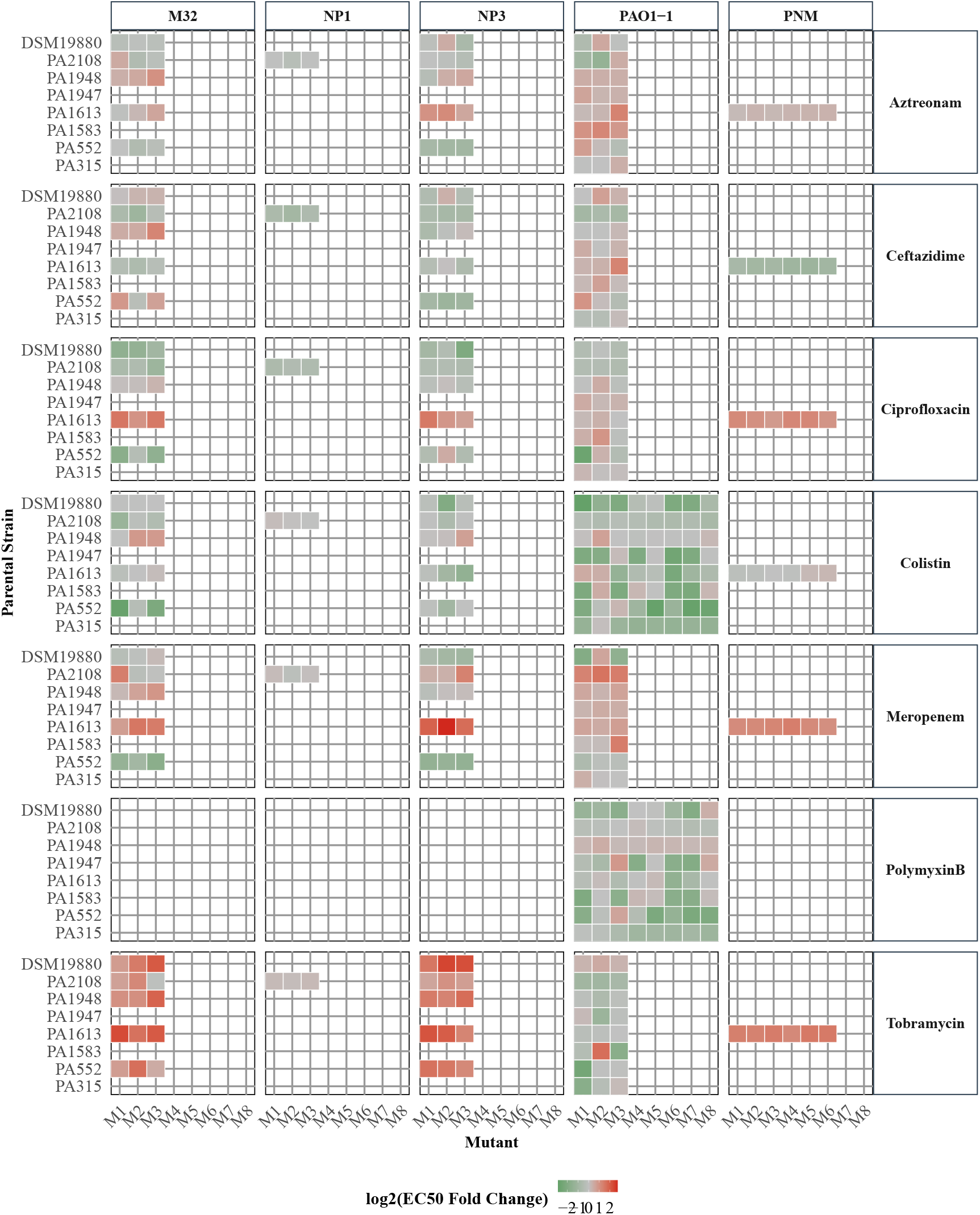
Overview of collateral effects in single phage-resistant mutants. The x-axis shows the mutant number for each parental strain (y-axis). For each individual mutant, the EC50_FC, log2_ , relative to its parent strain, is displayed for the antibiotics listed above. The right y-axis displays the phage to which each mutant is resistant. Values represent the mean EC50_FC, log2_ across three biological replicates. After log_2_ transformation, values *≤ −*0.5 indicate collateral sensitivity (CS; green), while values *≥* 0.5 indicate collateral resistance (CR; red).

### 11.8 Overview of collateral effects in double antibiotic- and phage-resistant mutants

**Supplementary Figure S11.**
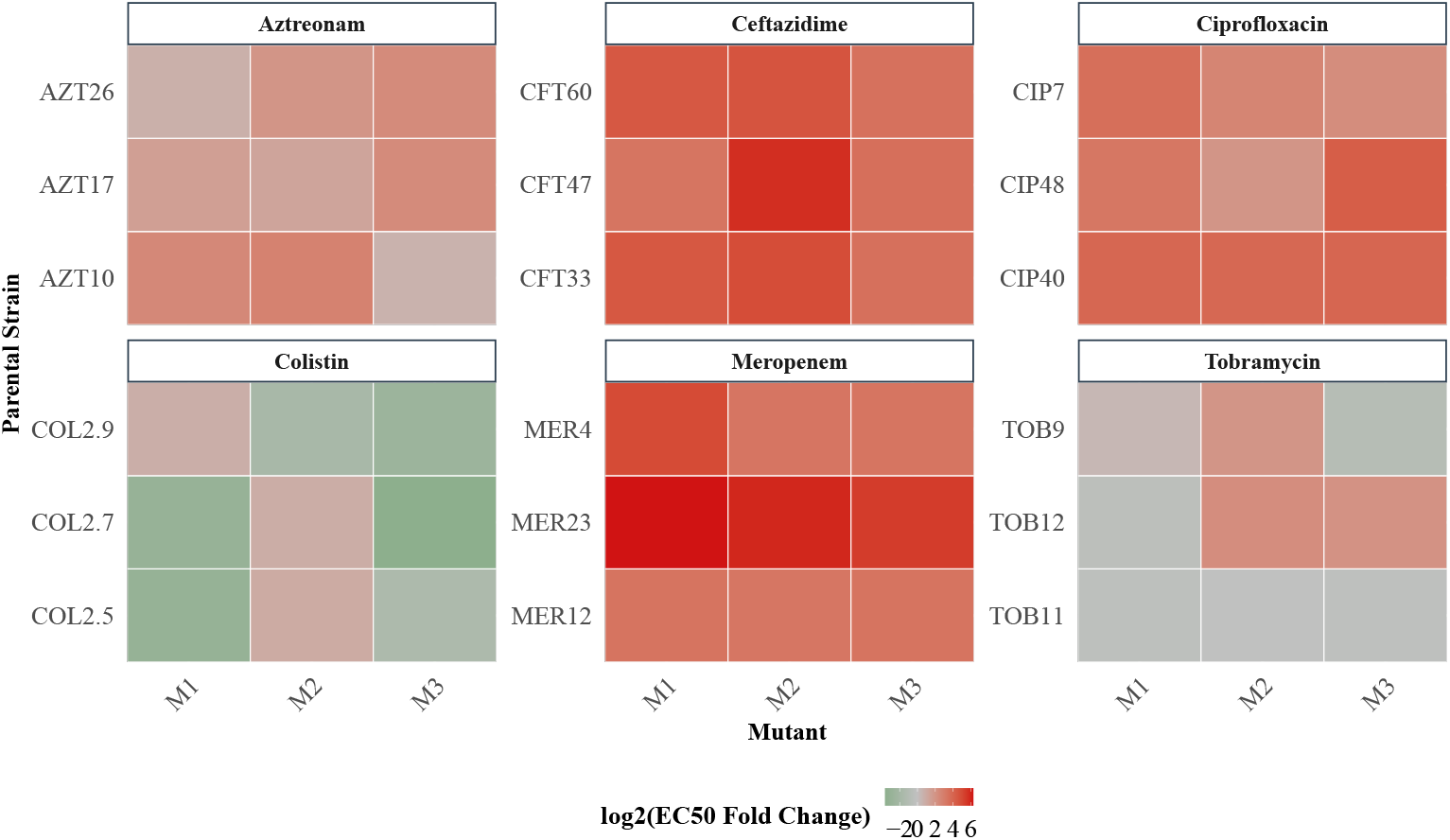
Overview of collateral effects in double antibiotic- and PAO1-1 phage-resistant mutants. The x-axis shows the individual mutant number for each parental strain (y-axis). For each individual mutant, the EC50_FC, log2_ , relative to its parent strain, is displayed for the antibiotics displayed on top. Values represent the mean EC50_FC, log2_ across three biological replicates. Green indicates collateral sensitivity (CS), whereas red indicates collateral resistance (CR). Values represent the mean EC50_FC, log2_ across three biological replicates.

### 11.9 Dose-response curves for phage PAO1-1 across antibiotic mutants

**Supplementary Figure S12.**
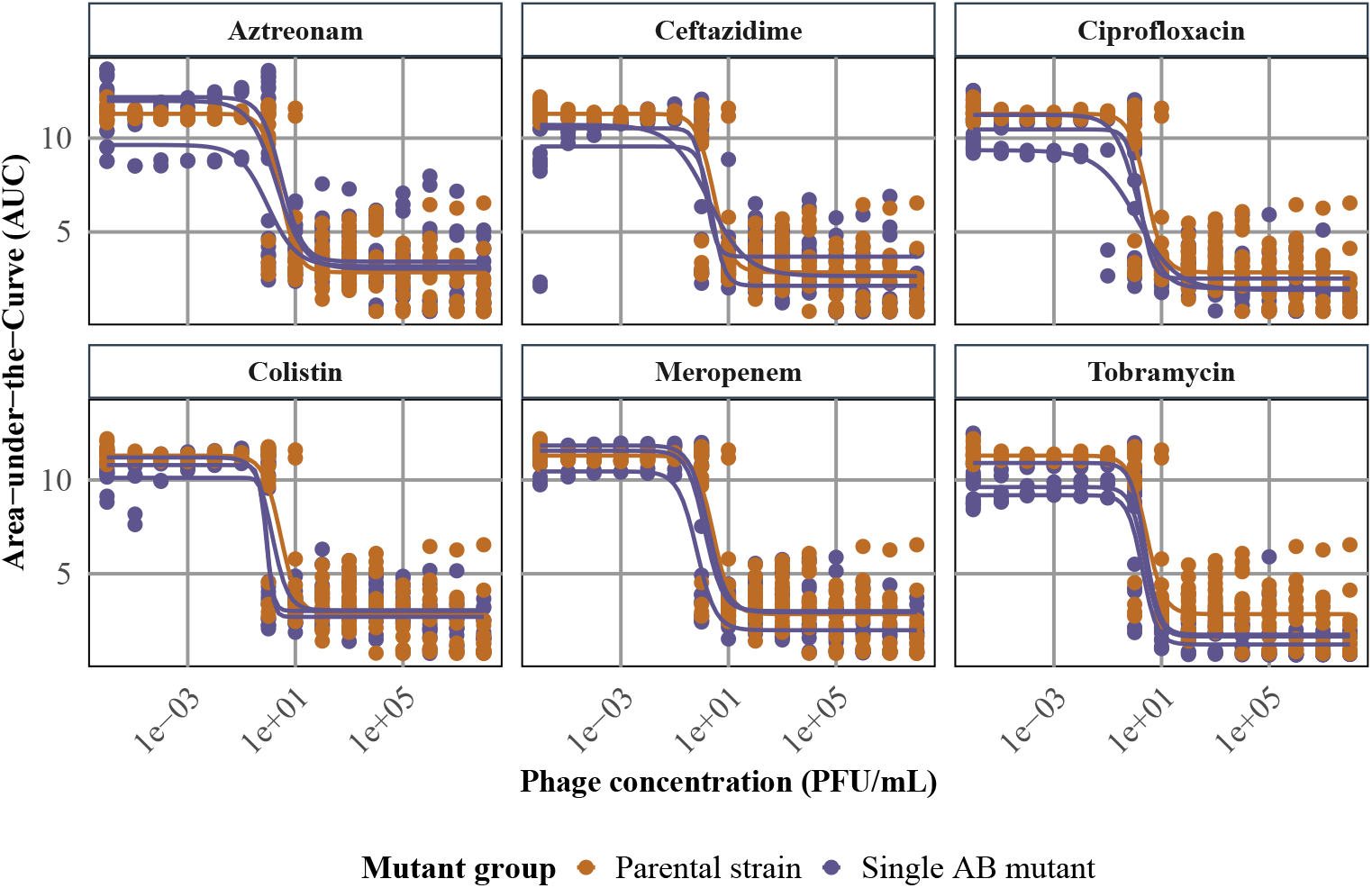
Dose-response curves for the PAO1-1 phage across single-antibiotic mutants. Dose–response curves (AUC vs. phage concentration, PFU/mL; log_10_ scale) for single antibiotic-resistant strains (purple) and the parental strain PA1947 (orange) against the PAO1-1 phage. The antibiotic to which each mutant was selected is indicated at the top. Curves were fitted using *drm* in R from three biological replicates per concentration.

### 11.10 Correlation of antibiotic EC50_FC, log2_ in PAO1-1 mutants

**Supplementary Figure S13.**
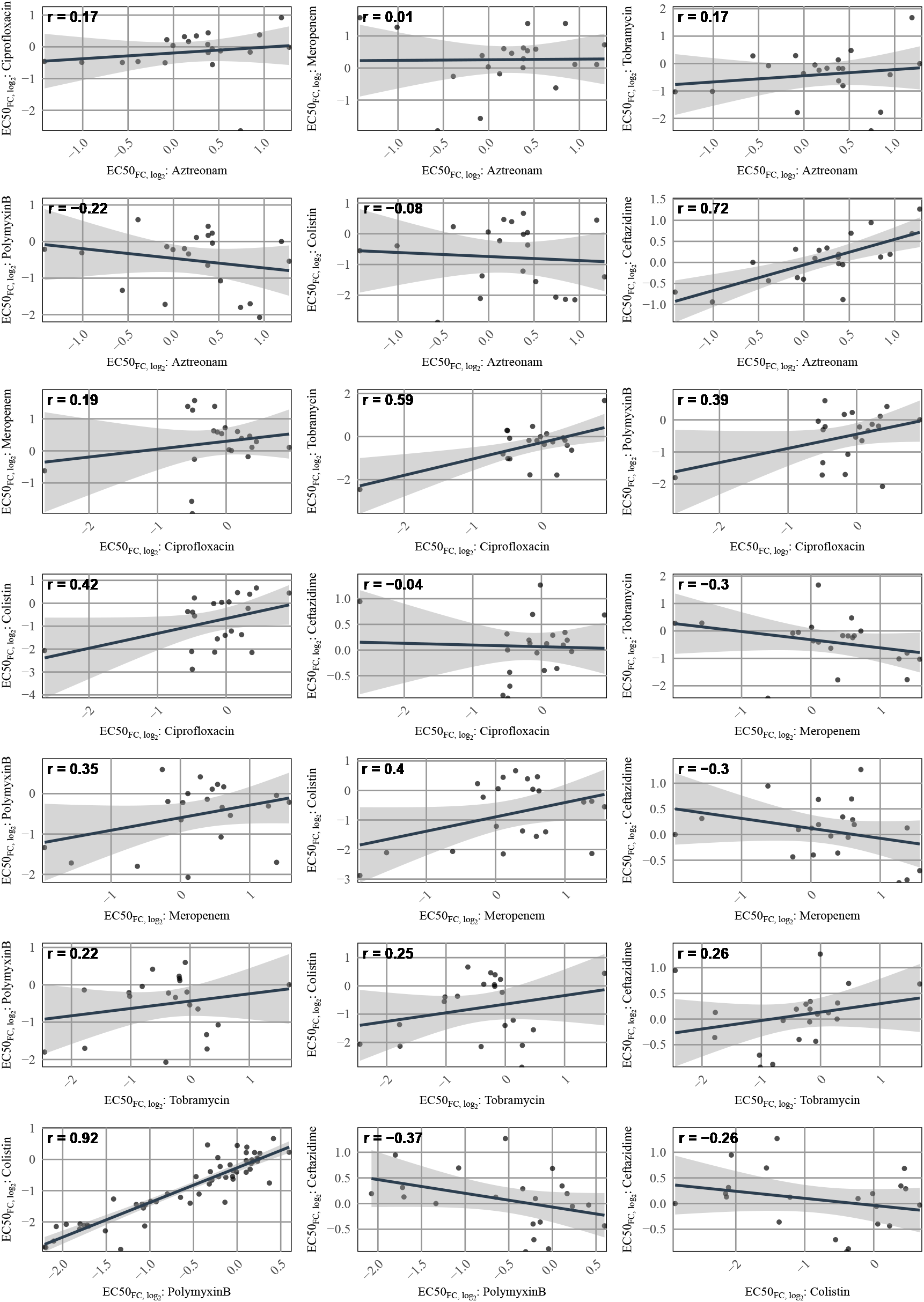
**Pearson correlation of antibiotic EC50_FC, log_**_2_ **in PAO1-1 phage-resistant mutants.** Each point represents the EC50_FC, log2_ of an individual mutant measured for the two antibiotics displayed. The regression line indicates the Pearson correlation between the two antibiotics, and the shaded area shows the standard error of the fit.

### 11.11 Fitness costs associated to phage- and antibiotic resistance

**Supplementary Figure S14.**
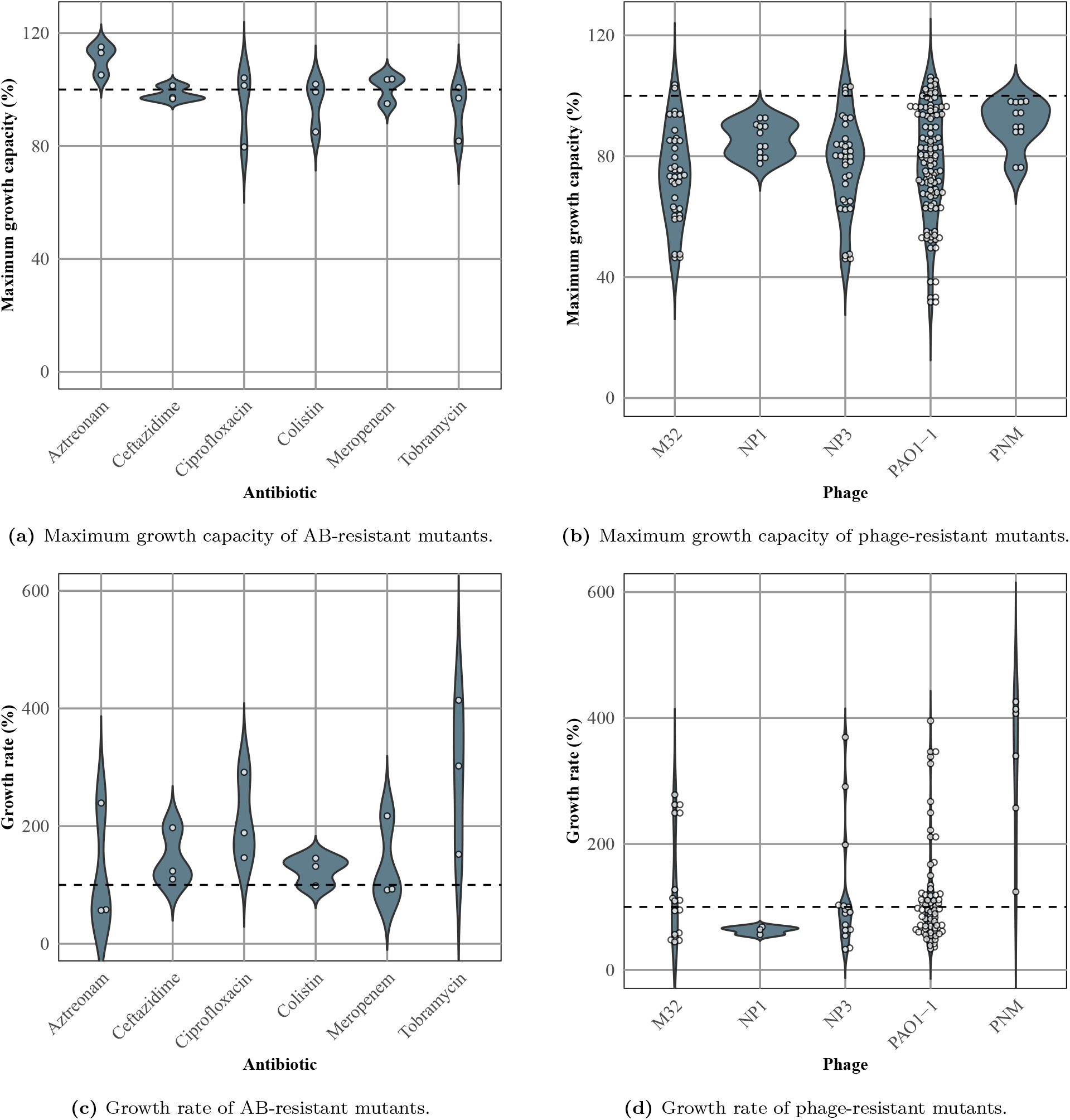
Comparison of fitness costs associated with resistance. Fitness is measured by (A,B) maximum growth capacity and (C,D) growth rate, each relative to the respective parental strain. Panels (A) and (C) show antibiotic-resistant mutants, and panels (B) and (D) show phage-resistant mutants. Dashed horizontal lines indicate the baseline for no fitness cost. Every data-point is the mean of three biological replicates.

### 11.12 Whole-genome sequencing

#### 11.12.1 Quality control: Baseread

**Supplementary Figure S15.**
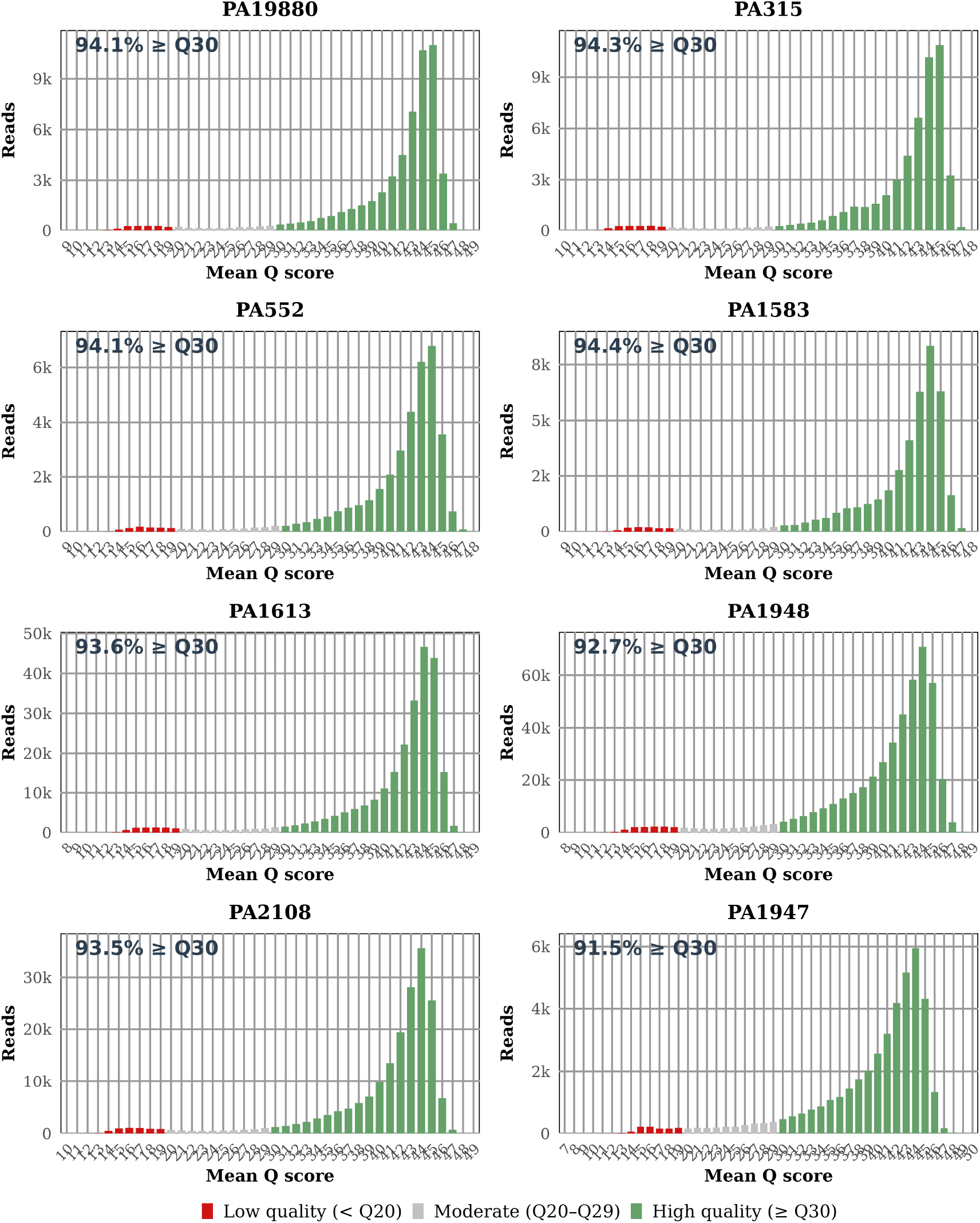
Mean quality score (Q value) per read for each parental strain sequenced by long-read Nanopore sequencing. The percentage of reads with *Q ≥* 30 is indicated in the upper left corner.

**Supplementary Figure S16.**
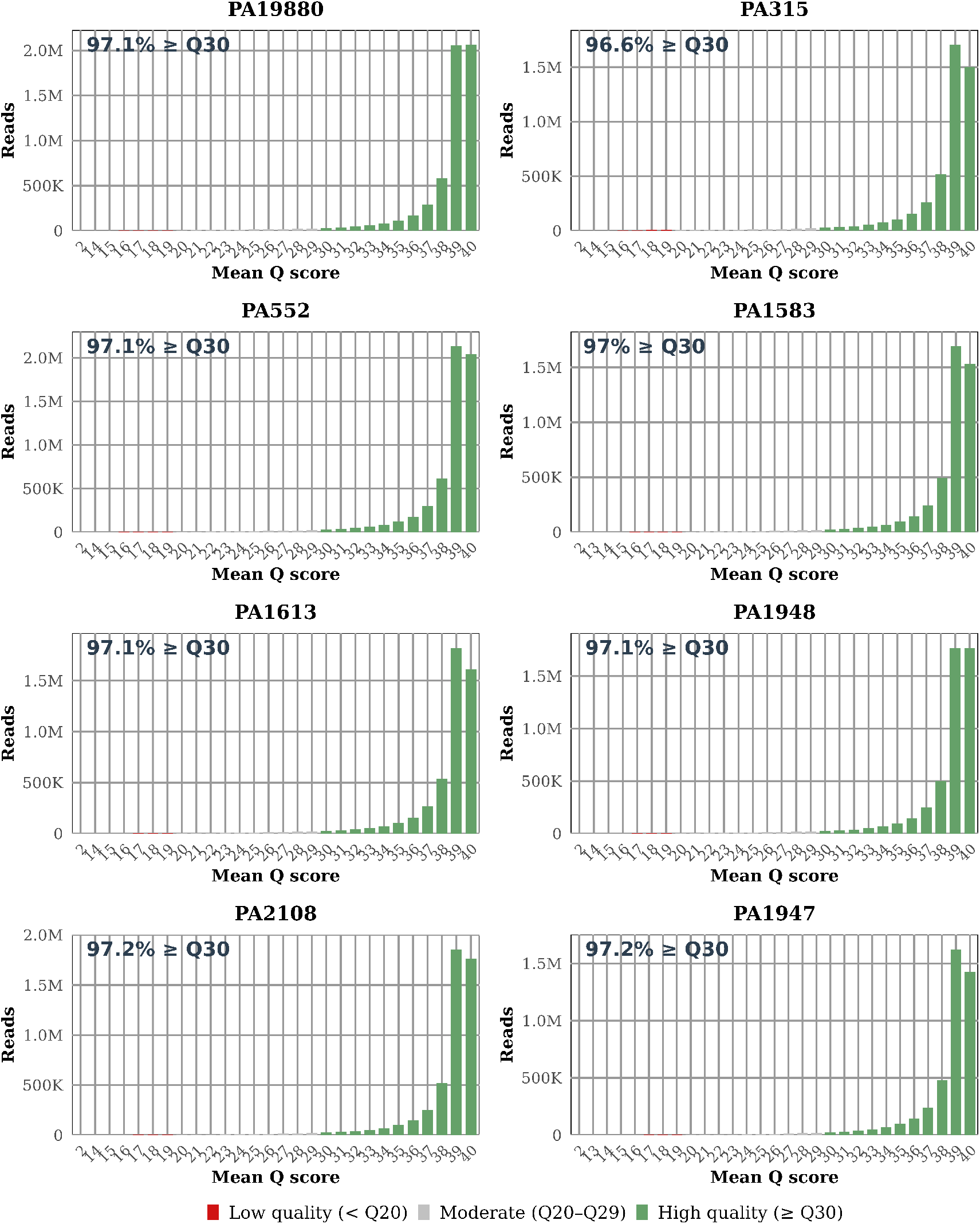
Mean quality score (Q value) per read for each parental strain sequenced by short-read Illumina sequencing. The percentage of reads with *Q ≥* 30 is indicated in the upper left corner.

**Supplementary Figure S17.**
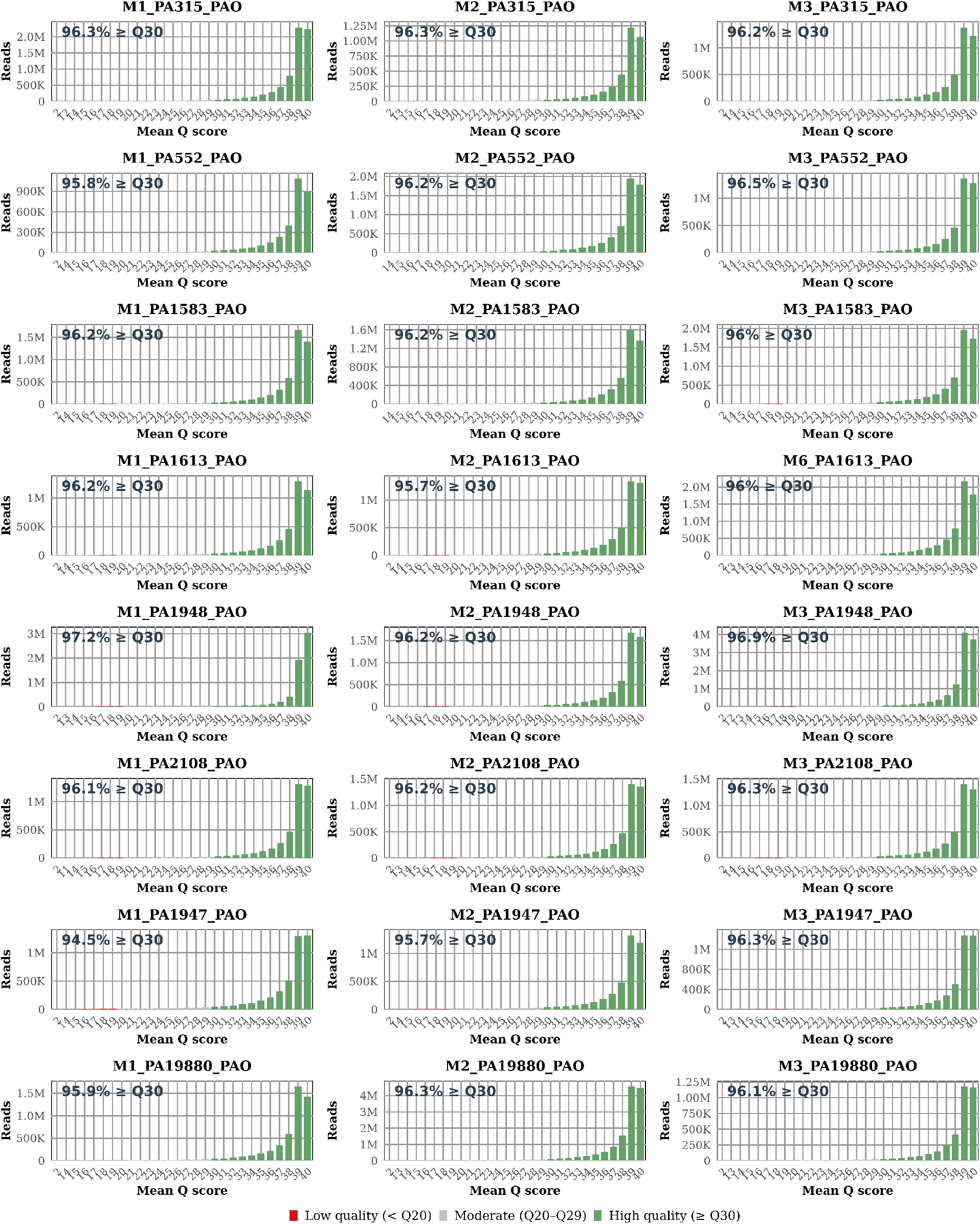
Mean quality score (Q value) per read for each single PAO1-1-resistant mutant sequenced by short-read Illumina sequencing. The percentage of reads with *Q ≥* 30 is indicated in the upper left corner.

#### 11.12.2 Quality control: Assembly

**Supplementary Figure S18.**
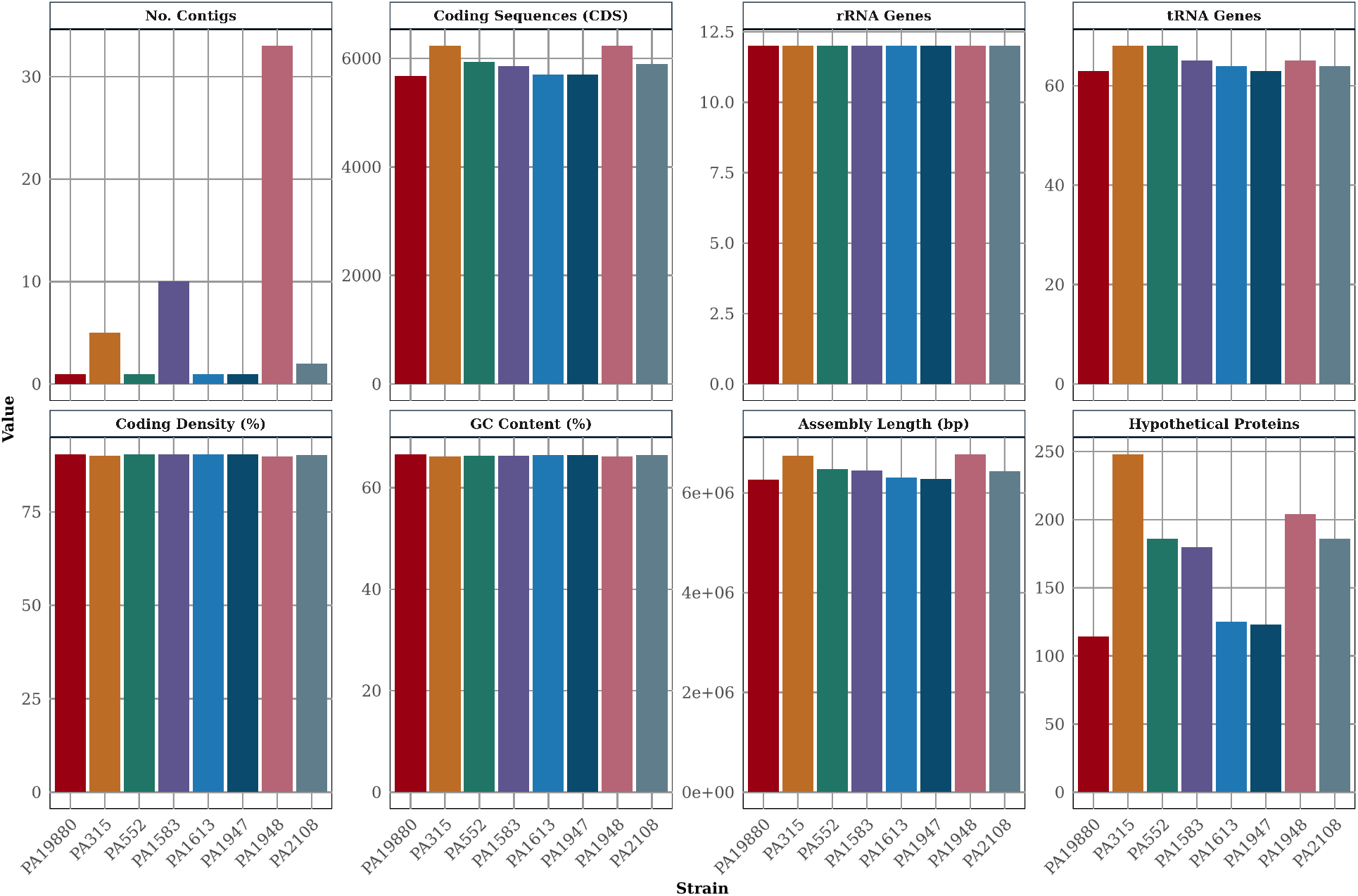
Assembly quality control metrics for parental strains sequenced by long-read Nanopore sequencing, showing the number of contigs, coding sequences (CDS), rRNAs, tRNAs, coding density, GC content, assembly length, and hypothetical proteins per strain.

**Supplementary Figure S19.**
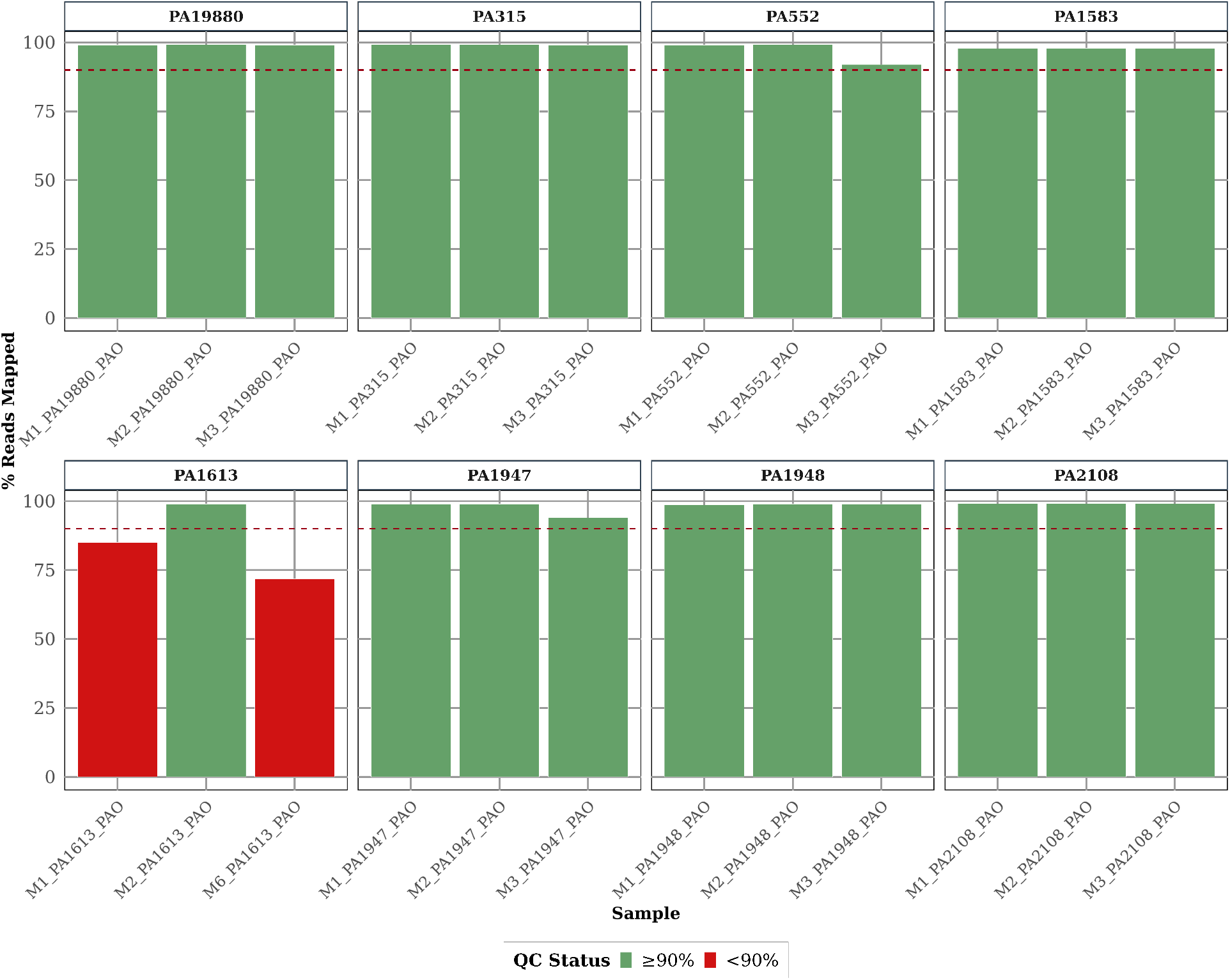
Short-read Illumina mapping coverage of three independent mutants per parental strain. Coverage is indicated as follows: green represents *≥*90% and red represents *<*90%. Strains M1 PA1613 PAO and M6 PA1613 PAO fall below the coverage threshold due to phage DNA contamination; however, as this does not affect variant calling, these samples are retained for downstream analysis.

#### 11.12.3 Number of mutations found across parental strains

**Supplementary Table S3.** Mutation profile of sequenced parental strains and their accompanying phage-resistant mutants.

| Parent | $n_{\text{mutations}}$ | $n_{\text{genes}}$ | $n_{\text{intergenic}}$ | $n_{\text{nonsynonymous}}$ | $n_{\text{synonymous}}$ | $n_{\text{nonsense}}$ |
| --- | --- | --- | --- | --- | --- | --- |
| DSM19880 | 24 | 1 | 1 | 5 | 7 | 1 |
| PA315 | 51 | 3 | 36 | 7 | 1 | 0 |
| PA552 | 24 | 4 | 11 | 3 | 7 | 0 |
| PA1583 | 24 | 5 | 2 | 7 | 11 | 1 |
| PA1613 | 18 | 3 | 7 | 7 | 2 | 0 |
| PA1947 | 7 | 0 | 1 | 2 | 4 | 0 |
| PA1948 | 21 | 1 | 17 | 1 | 2 | 0 |
| PA2108 | 122 | 25 | 25 | 63 | 26 | 0 |

$n_{\text{mutations}}$ , total number of mutations within all three phage-resistant mutants per parental strain; $n_{\text{genes}}$ , number of shared genes mutated for all three phage-resistant mutants within the same parental strain; $n_{\text{intergenic}}$ , $n_{\text{nonsynonymous}}$ , $n_{\text{synonymous}}$ , $n_{\text{nonsense}}$ refer to the predicted functional effect of each SNP to be located in intergenic regions or to be nonsynonymous, synonymous, or nonsense. Information for the remaining mutants regarding substitutions, insertions, and deletions is found in the supplementary.

### 11.13 Statistical tests

#### 11.13.1 Linear regression model for collateral effects among antibiotics and parental strains

**Supplementary Table S4.**
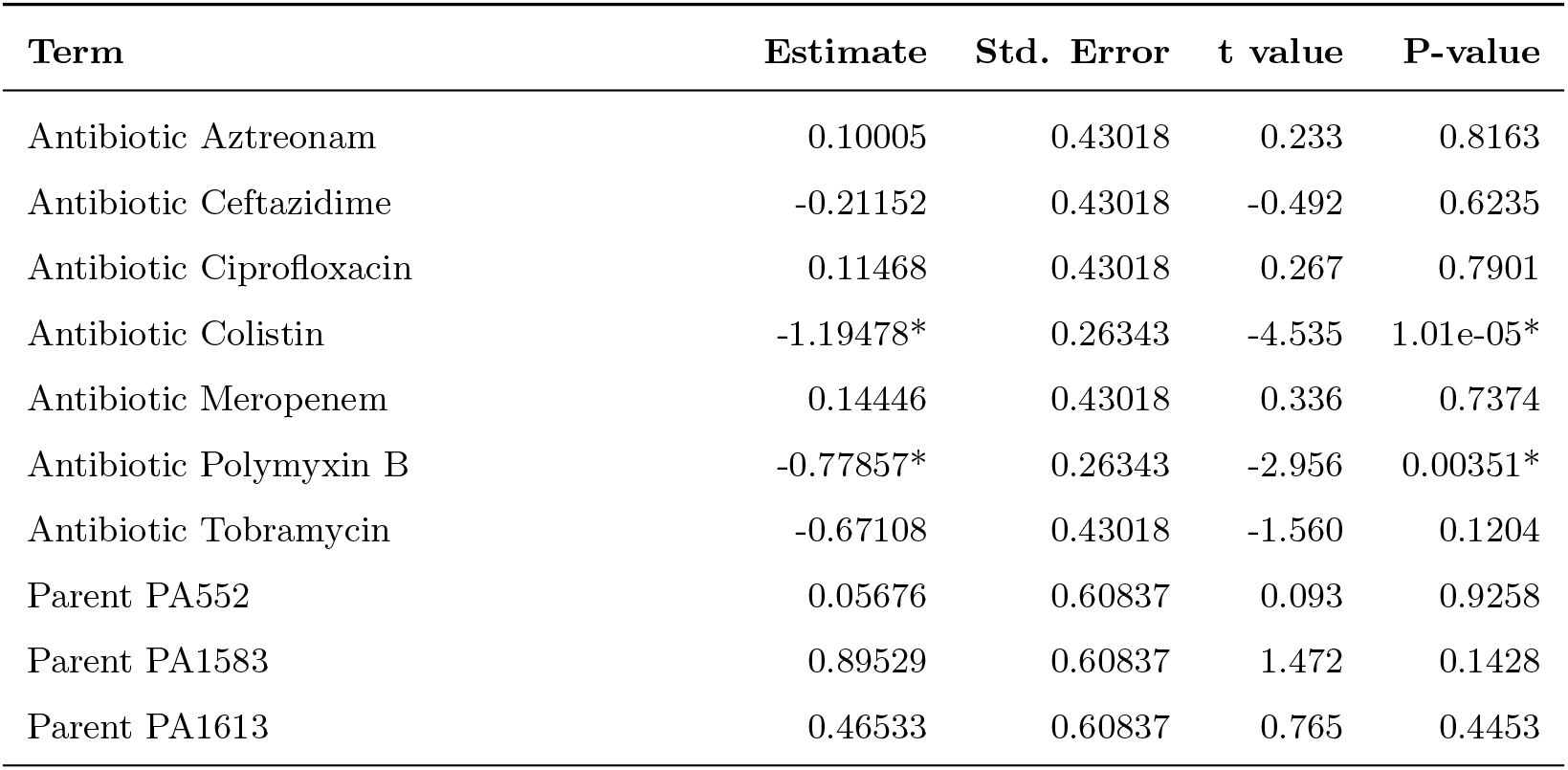
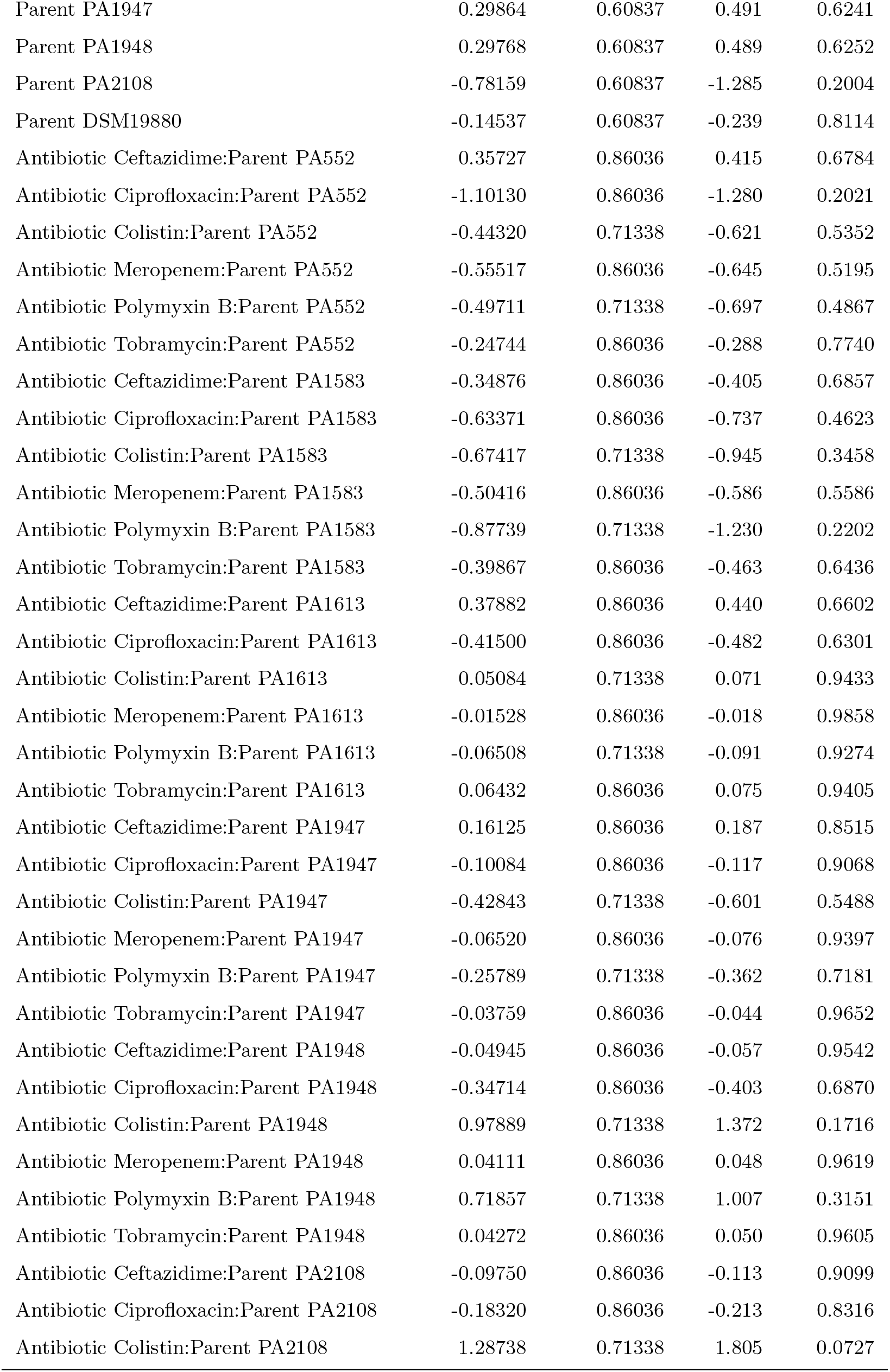
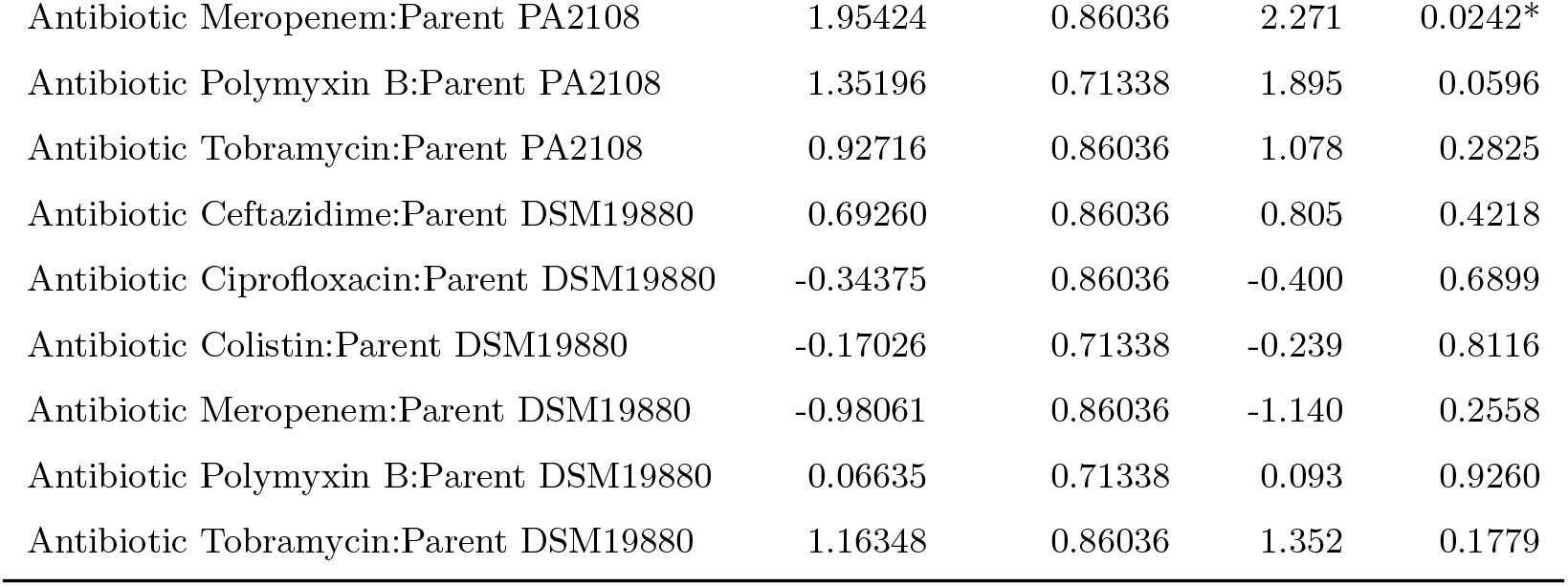
Regression estimates for log_2_(EC50 FC) differences in single PAO1-1 mutants. To investigate the effect of antibiotics on the log_2_ EC50 fold-change in single PAO1-1 mutants, a linear regression model was fitted for each antibiotic, parent, and their interaction. Significant terms (p *<* 0.05) are indicated with *.

| Term | Estimate | Std. Error | t value | P-value |
| --- | --- | --- | --- | --- |
| Antibiotic Aztreonam | 0.10005 | 0.43018 | 0.233 | 0.8163 |
| Antibiotic Ceftazidime | -0.21152 | 0.43018 | -0.492 | 0.6235 |
| Antibiotic Ciprofloxacin | 0.11468 | 0.43018 | 0.267 | 0.7901 |
| Antibiotic Colistin | -1.19478* | 0.26343 | -4.535 | 1.01e-05* |
| Antibiotic Meropenem | 0.14446 | 0.43018 | 0.336 | 0.7374 |
| Antibiotic Polymyxin B | -0.77857* | 0.26343 | -2.956 | 0.00351* |
| Antibiotic Tobramycin | -0.67108 | 0.43018 | -1.560 | 0.1204 |
| Parent PA552 | 0.05676 | 0.60837 | 0.093 | 0.9258 |
| Parent PA1583 | 0.89529 | 0.60837 | 1.472 | 0.1428 |
| Parent PA1613 | 0.46533 | 0.60837 | 0.765 | 0.4453 |
| Parent PA1947 | 0.29864 | 0.60837 | 0.491 | 0.6241 |
| Parent PA1948 | 0.29768 | 0.60837 | 0.489 | 0.6252 |
| Parent PA2108 | -0.78159 | 0.60837 | -1.285 | 0.2004 |
| Parent DSM19880 | -0.14537 | 0.60837 | -0.239 | 0.8114 |
| Antibiotic Ceftazidime:Parent PA552 | 0.35727 | 0.86036 | 0.415 | 0.6784 |
| Antibiotic Ciprofloxacin:Parent PA552 | -1.10130 | 0.86036 | -1.280 | 0.2021 |
| Antibiotic Colistin:Parent PA552 | -0.44320 | 0.71338 | -0.621 | 0.5352 |
| Antibiotic Meropenem:Parent PA552 | -0.55517 | 0.86036 | -0.645 | 0.5195 |
| Antibiotic Polymyxin B:Parent PA552 | -0.49711 | 0.71338 | -0.697 | 0.4867 |
| Antibiotic Tobramycin:Parent PA552 | -0.24744 | 0.86036 | -0.288 | 0.7740 |
| Antibiotic Ceftazidime:Parent PA1583 | -0.34876 | 0.86036 | -0.405 | 0.6857 |
| Antibiotic Ciprofloxacin:Parent PA1583 | -0.63371 | 0.86036 | -0.737 | 0.4623 |
| Antibiotic Colistin:Parent PA1583 | -0.67417 | 0.71338 | -0.945 | 0.3458 |
| Antibiotic Meropenem:Parent PA1583 | -0.50416 | 0.86036 | -0.586 | 0.5586 |
| Antibiotic Polymyxin B:Parent PA1583 | -0.87739 | 0.71338 | -1.230 | 0.2202 |
| Antibiotic Tobramycin:Parent PA1583 | -0.39867 | 0.86036 | -0.463 | 0.6436 |
| Antibiotic Ceftazidime:Parent PA1613 | 0.37882 | 0.86036 | 0.440 | 0.6602 |
| Antibiotic Ciprofloxacin:Parent PA1613 | -0.41500 | 0.86036 | -0.482 | 0.6301 |
| Antibiotic Colistin:Parent PA1613 | 0.05084 | 0.71338 | 0.071 | 0.9433 |
| Antibiotic Meropenem:Parent PA1613 | -0.01528 | 0.86036 | -0.018 | 0.9858 |
| Antibiotic Polymyxin B:Parent PA1613 | -0.06508 | 0.71338 | -0.091 | 0.9274 |
| Antibiotic Tobramycin:Parent PA1613 | 0.06432 | 0.86036 | 0.075 | 0.9405 |
| Antibiotic Ceftazidime:Parent PA1947 | 0.16125 | 0.86036 | 0.187 | 0.8515 |
| Antibiotic Ciprofloxacin:Parent PA1947 | -0.10084 | 0.86036 | -0.117 | 0.9068 |
| Antibiotic Colistin:Parent PA1947 | -0.42843 | 0.71338 | -0.601 | 0.5488 |
| Antibiotic Meropenem:Parent PA1947 | -0.06520 | 0.86036 | -0.076 | 0.9397 |
| Antibiotic Polymyxin B:Parent PA1947 | -0.25789 | 0.71338 | -0.362 | 0.7181 |
| Antibiotic Tobramycin:Parent PA1947 | -0.03759 | 0.86036 | -0.044 | 0.9652 |
| Antibiotic Ceftazidime:Parent PA1948 | -0.04945 | 0.86036 | -0.057 | 0.9542 |
| Antibiotic Ciprofloxacin:Parent PA1948 | -0.34714 | 0.86036 | -0.403 | 0.6870 |
| Antibiotic Colistin:Parent PA1948 | 0.97889 | 0.71338 | 1.372 | 0.1716 |
| Antibiotic Meropenem:Parent PA1948 | 0.04111 | 0.86036 | 0.048 | 0.9619 |
| Antibiotic Polymyxin B:Parent PA1948 | 0.71857 | 0.71338 | 1.007 | 0.3151 |
| Antibiotic Tobramycin:Parent PA1948 | 0.04272 | 0.86036 | 0.050 | 0.9605 |
| Antibiotic Ceftazidime:Parent PA2108 | -0.09750 | 0.86036 | -0.113 | 0.9099 |
| Antibiotic Ciprofloxacin:Parent PA2108 | -0.18320 | 0.86036 | -0.213 | 0.8316 |
| Antibiotic Colistin:Parent PA2108 | 1.28738 | 0.71338 | 1.805 | 0.0727 |

| **Term** | **Estimate** | **Std. Error** | **t value** | **P-value** |
| --- | --- | --- | --- | --- |
| Antibiotic Meropenem:Parent PA2108 | 1.95424 | 0.86036 | 2.271 | 0.0242* |
| Antibiotic Polymyxin B:Parent PA2108 | 1.35196 | 0.71338 | 1.895 | 0.0596 |
| Antibiotic Tobramycin:Parent PA2108 | 0.92716 | 0.86036 | 1.078 | 0.2825 |
| Antibiotic Ceftazidime:Parent DSM19880 | 0.69260 | 0.86036 | 0.805 | 0.4218 |
| Antibiotic Ciprofloxacin:Parent DSM19880 | -0.34375 | 0.86036 | -0.400 | 0.6899 |
| Antibiotic Colistin:Parent DSM19880 | -0.17026 | 0.71338 | -0.239 | 0.8116 |
| Antibiotic Meropenem:Parent DSM19880 | -0.98061 | 0.86036 | -1.140 | 0.2558 |
| Antibiotic Polymyxin B:Parent DSM19880 | 0.06635 | 0.71338 | 0.093 | 0.9260 |
| Antibiotic Tobramycin:Parent DSM19880 | 1.16348 | 0.86036 | 1.352 | 0.1779 |

**Supplementary Table S5.** Post hoc Tukey test for pairwise comparisons of Parent × Antibiotic combinations. Reported are mean differences in log_2_(EC50 fold change), lower and upper 95% confidence intervals (lwr, upr), and adjusted p-values (p adj). Significant comparisons (p*<*0.05) are indicated with *. Only the comparisons with significant p-values are shown, and from here, the comparisons are relevant within the same antibiotic indicated in **bold**.

| Comparison | diff | lwr | upr | p adj |
| --- | --- | --- | --- | --- |
| Aztreonam:PA1583–Colistin:PA315 | 2.1901 | 0.1138 | 4.2664 | 0.0228* |
| Meropenem:PA2108–Colistin:PA315 | 2.5119 | 0.4356 | 4.5882 | 0.0018* |
| Meropenem:PA2108–PolymyxinB:PA315 | 2.0957 | 0.0194 | 4.1720 | 0.0439* |
| Aztreonam:PA1583–Colistin:PA552 | 2.5766 | 0.5003 | 4.6529 | 0.0010* |
| Meropenem:PA1583–Colistin:PA552 | 2.1168 | 0.0405 | 4.1931 | 0.0381* |
| Aztreonam:PA1613–Colistin:PA552 | 2.1466 | 0.0703 | 4.2229 | 0.0310* |
| Ceftazidime:PA1613–Colistin:PA552 | 2.2138 | 0.1376 | 4.2901 | 0.0192* |
| Meropenem:PA1613–Colistin:PA552 | 2.1757 | 0.0994 | 4.2520 | 0.0253* |
| **Colistin:PA1948–Colistin:PA552** | 1.6630 | 0.1296 | 3.1965 | 0.0145* |
| PolymyxinB:PA1948–Colistin:PA552 | 1.8189 | 0.2854 | 3.3523 | 0.0027* |
| Meropenem:PA2108–Colistin:PA552 | 2.8983 | 0.8220 | 4.9746 | <0.001* |
| Aztreonam:PA1583–PolymyxinB:PA552 | 2.2143 | 0.1380 | 4.2906 | 0.0191* |
| Meropenem:PA2108–PolymyxinB:PA552 | 2.5360 | 0.4597 | 4.6123 | 0.0014* |
| Colistin:PA1947–Aztreonam:PA1583 | -2.3199 | -4.3962 | -0.2436 | 0.0086* |
| Colistin:DSM19880–Aztreonam:PA1583 | -2.5058 | -4.5821 | -0.4295 | 0.0019* |
| Meropenem:PA2108–Colistin:PA1583 | 2.2908 | 0.2145 | 4.3671 | 0.0108* |
| Meropenem:PA2108–PolymyxinB:PA1583 | 2.0778 | 0.0015 | 4.1541 | 0.0495* |
| Colistin:DSM19880–Ceftazidime:PA1613 | -2.1430 | -4.2193 | -0.0667 | 0.0318* |
| Colistin:DSM19880–Meropenem:PA1613 | -2.1049 | -4.1812 | -0.0286 | 0.0413* |
| PolymyxinB:PA1948–Colistin:PA1947 | 1.5622 | 0.0288 | 3.0957 | 0.0385* |
| Meropenem:PA2108–Colistin:PA1947 | 2.6417 | 0.5654 | 4.7180 | 0.0006* |
| **Colistin:DSM19880–Colistin:PA1948** | -1.5922 | -3.1256 | -0.0588 | 0.0291* |
| Colistin:DSM19880–PolymyxinB:PA1948 | -1.7481 | -3.2815 | -0.2146 | 0.0059* |
| Colistin:DSM19880–Meropenem:PA2108 | -2.8275 | -4.9038 | -0.7512 | <0.001* |
| PolymyxinB:DSM19880–Meropenem:PA2108 | -2.1747 | -4.2510 | -0.0984 | 0.0254* |

#### 11.13.2 Linear mixed model for phage differences between antibiotics

**Supplementary Table S6.**
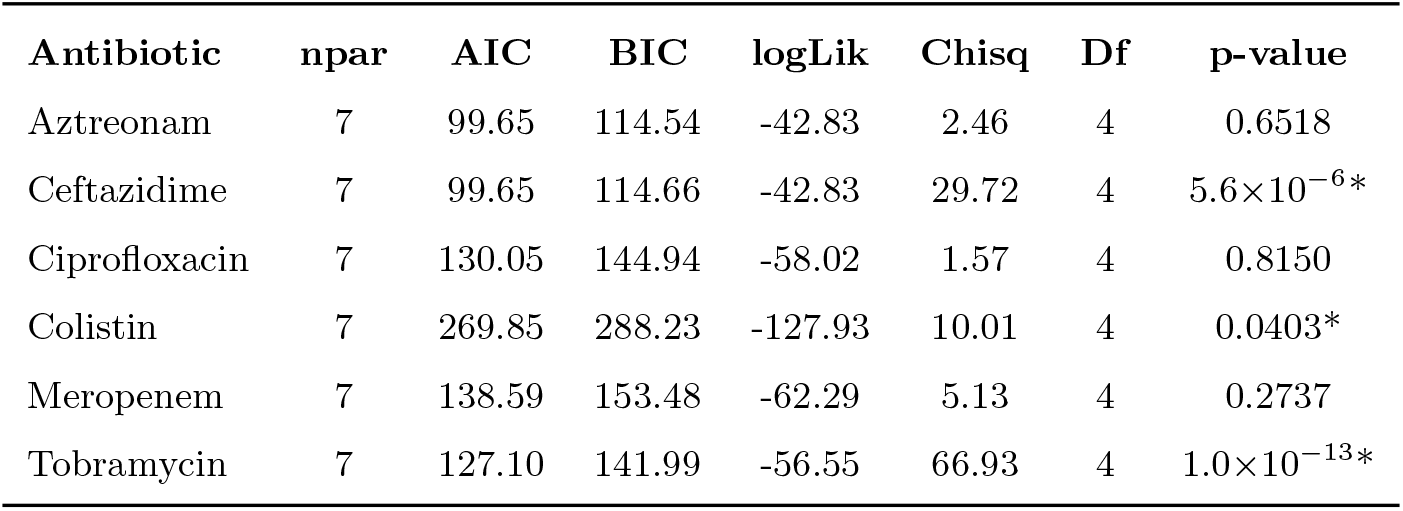
Model comparison between random-intercept and phage-effect models for each antibiotic. Two mixed-effects models were compared using ANOVA: (1) a null model including only random intercepts for parental strains, and (2) a full model including phage effects. Shown are AIC (Akaike Information Criterion), BIC (Bayesian Information Criterion), log-likelihood, Chi-square statistics, and corresponding p-values. Significant differences (p*<*0.05) are indicated with *.

| Antibiotic | npar | AIC | BIC | logLik | Chisq | Df | p-value |
| --- | --- | --- | --- | --- | --- | --- | --- |
| Aztreonam | 7 | 99.65 | 114.54 | -42.83 | 2.46 | 4 | 0.6518 |
| Ceftazidime | 7 | 99.65 | 114.66 | -42.83 | 29.72 | 4 | $5.6 \times 10^{-6}$* |
| Ciprofloxacin | 7 | 130.05 | 144.94 | -58.02 | 1.57 | 4 | 0.8150 |
| Colistin | 7 | 269.85 | 288.23 | -127.93 | 10.01 | 4 | 0.0403* |
| Meropenem | 7 | 138.59 | 153.48 | -62.29 | 5.13 | 4 | 0.2737 |
| Tobramycin | 7 | 127.10 | 141.99 | -56.55 | 66.93 | 4 | $1.0 \times 10^{-13}$* |

**Supplementary Table S7.**
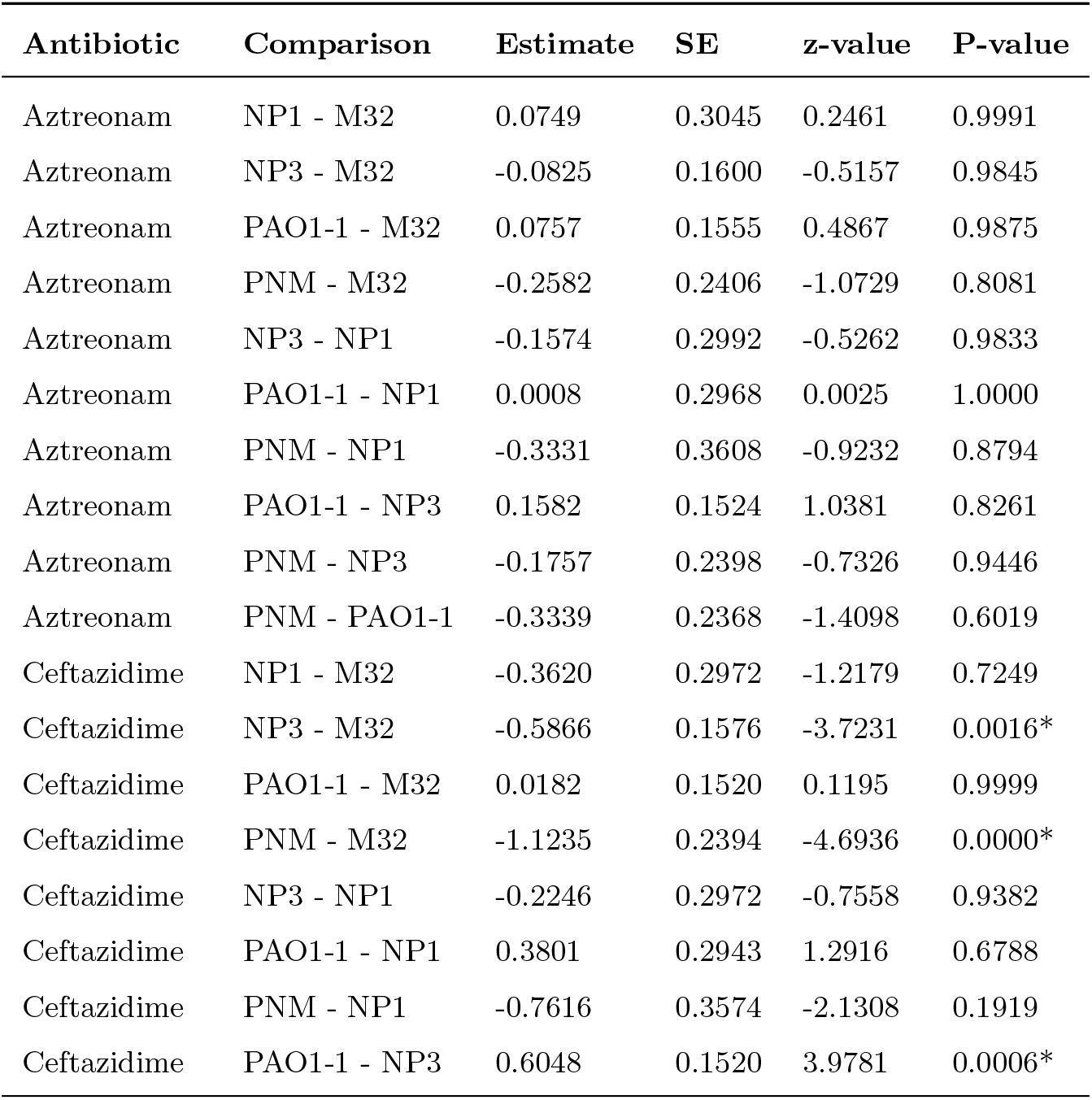
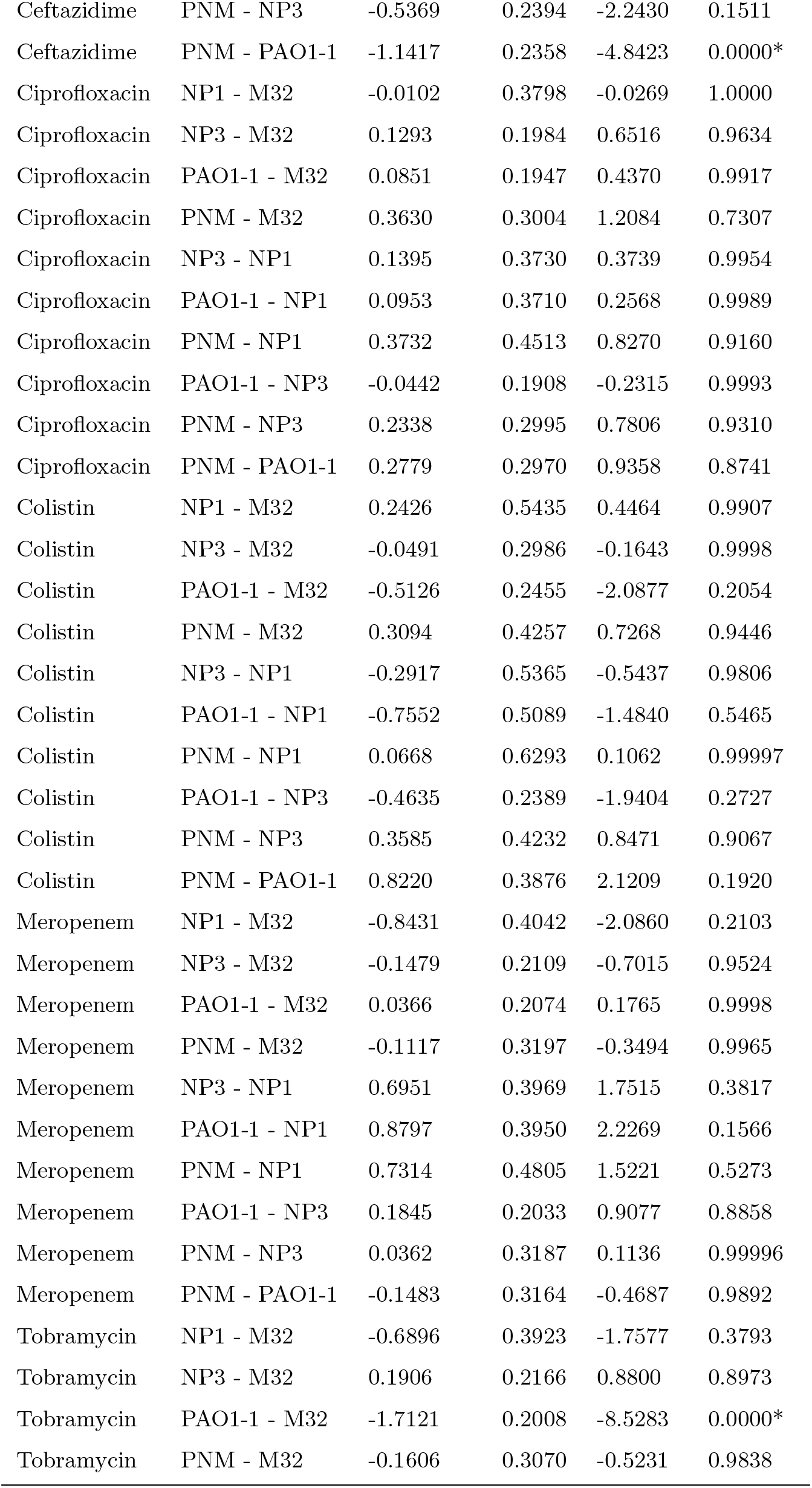
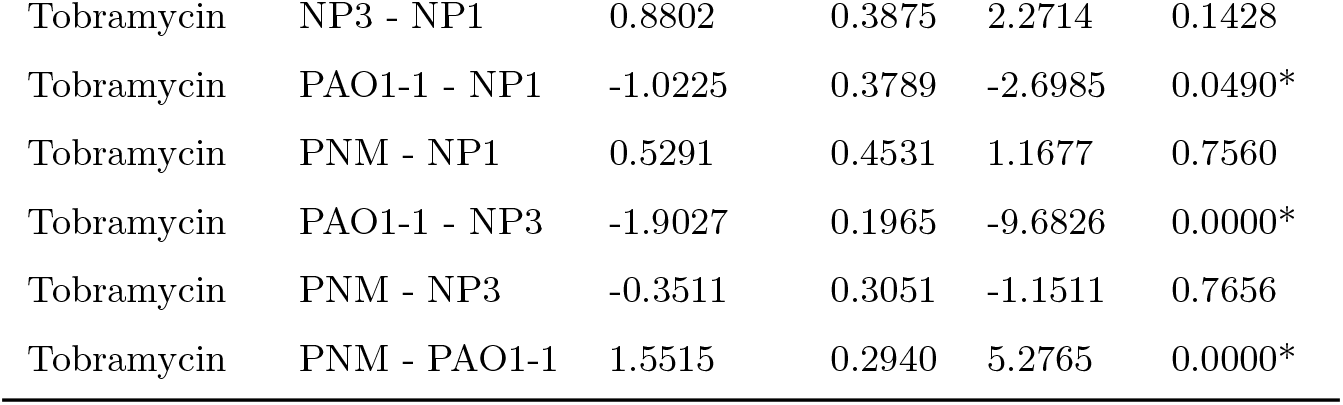
Pairwise comparisons of different single-phage mutants per antibiotic. Estimated contrasts, standard errors (SE), z-values, and p-values are reported. Significant contrasts (p*<*0.05) are marked with *.

| Antibiotic | Comparison | Estimate | SE | z-value | P-value |
| --- | --- | --- | --- | --- | --- |
| Aztreonam | NP1 - M32 | 0.0749 | 0.3045 | 0.2461 | 0.9991 |
| Aztreonam | NP3 - M32 | -0.0825 | 0.1600 | -0.5157 | 0.9845 |
| Aztreonam | PAO1-1 - M32 | 0.0757 | 0.1555 | 0.4867 | 0.9875 |
| Aztreonam | PNM - M32 | -0.2582 | 0.2406 | -1.0729 | 0.8081 |
| Aztreonam | NP3 - NP1 | -0.1574 | 0.2992 | -0.5262 | 0.9833 |
| Aztreonam | PAO1-1 - NP1 | 0.0008 | 0.2968 | 0.0025 | 1.0000 |
| Aztreonam | PNM - NP1 | -0.3331 | 0.3608 | -0.9232 | 0.8794 |
| Aztreonam | PAO1-1 - NP3 | 0.1582 | 0.1524 | 1.0381 | 0.8261 |
| Aztreonam | PNM - NP3 | -0.1757 | 0.2398 | -0.7326 | 0.9446 |
| Aztreonam | PNM - PAO1-1 | -0.3339 | 0.2368 | -1.4098 | 0.6019 |
| Ceftazidime | NP1 - M32 | -0.3620 | 0.2972 | -1.2179 | 0.7249 |
| Ceftazidime | NP3 - M32 | -0.5866 | 0.1576 | -3.7231 | 0.0016* |
| Ceftazidime | PAO1-1 - M32 | 0.0182 | 0.1520 | 0.1195 | 0.9999 |
| Ceftazidime | PNM - M32 | -1.1235 | 0.2394 | -4.6936 | 0.0000* |
| Ceftazidime | NP3 - NP1 | -0.2246 | 0.2972 | -0.7558 | 0.9382 |
| Ceftazidime | PAO1-1 - NP1 | 0.3801 | 0.2943 | 1.2916 | 0.6788 |
| Ceftazidime | PNM - NP1 | -0.7616 | 0.3574 | -2.1308 | 0.1919 |
| Ceftazidime | PAO1-1 - NP3 | 0.6048 | 0.1520 | 3.9781 | 0.0006* |
| Ceftazidime | PNM - NP3 | -0.5369 | 0.2394 | -2.2430 | 0.1511 |
| Ceftazidime | PNM - PAO1-1 | -1.1417 | 0.2358 | -4.8423 | 0.0000* |
| Ciprofloxacin | NP1 - M32 | -0.0102 | 0.3798 | -0.0269 | 1.0000 |
| Ciprofloxacin | NP3 - M32 | 0.1293 | 0.1984 | 0.6516 | 0.9634 |
| Ciprofloxacin | PAO1-1 - M32 | 0.0851 | 0.1947 | 0.4370 | 0.9917 |
| Ciprofloxacin | PNM - M32 | 0.3630 | 0.3004 | 1.2084 | 0.7307 |
| Ciprofloxacin | NP3 - NP1 | 0.1395 | 0.3730 | 0.3739 | 0.9954 |
| Ciprofloxacin | PAO1-1 - NP1 | 0.0953 | 0.3710 | 0.2568 | 0.9989 |
| Ciprofloxacin | PNM - NP1 | 0.3732 | 0.4513 | 0.8270 | 0.9160 |
| Ciprofloxacin | PAO1-1 - NP3 | -0.0442 | 0.1908 | -0.2315 | 0.9993 |
| Ciprofloxacin | PNM - NP3 | 0.2338 | 0.2995 | 0.7806 | 0.9310 |
| Ciprofloxacin | PNM - PAO1-1 | 0.2779 | 0.2970 | 0.9358 | 0.8741 |
| Colistin | NP1 - M32 | 0.2426 | 0.5435 | 0.4464 | 0.9907 |
| Colistin | NP3 - M32 | -0.0491 | 0.2986 | -0.1643 | 0.9998 |
| Colistin | PAO1-1 - M32 | -0.5126 | 0.2455 | -2.0877 | 0.2054 |
| Colistin | PNM - M32 | 0.3094 | 0.4257 | 0.7268 | 0.9446 |
| Colistin | NP3 - NP1 | -0.2917 | 0.5365 | -0.5437 | 0.9806 |
| Colistin | PAO1-1 - NP1 | -0.7552 | 0.5089 | -1.4840 | 0.5465 |
| Colistin | PNM - NP1 | 0.0668 | 0.6293 | 0.1062 | 0.99997 |
| Colistin | PAO1-1 - NP3 | -0.4635 | 0.2389 | -1.9404 | 0.2727 |
| Colistin | PNM - NP3 | 0.3585 | 0.4232 | 0.8471 | 0.9067 |
| Colistin | PNM - PAO1-1 | 0.8220 | 0.3876 | 2.1209 | 0.1920 |
| Meropenem | NP1 - M32 | -0.8431 | 0.4042 | -2.0860 | 0.2103 |
| Meropenem | NP3 - M32 | -0.1479 | 0.2109 | -0.7015 | 0.9524 |
| Meropenem | PAO1-1 - M32 | 0.0366 | 0.2074 | 0.1765 | 0.9998 |
| Meropenem | PNM - M32 | -0.1117 | 0.3197 | -0.3494 | 0.9965 |
| Meropenem | NP3 - NP1 | 0.6951 | 0.3969 | 1.7515 | 0.3817 |
| Meropenem | PAO1-1 - NP1 | 0.8797 | 0.3950 | 2.2269 | 0.1566 |
| Meropenem | PNM - NP1 | 0.7314 | 0.4805 | 1.5221 | 0.5273 |
| Meropenem | PAO1-1 - NP3 | 0.1845 | 0.2033 | 0.9077 | 0.8858 |
| Meropenem | PNM - NP3 | 0.0362 | 0.3187 | 0.1136 | 0.99996 |
| Meropenem | PNM - PAO1-1 | -0.1483 | 0.3164 | -0.4687 | 0.9892 |
| Tobramycin | NP1 - M32 | -0.6896 | 0.3923 | -1.7577 | 0.3793 |
| Tobramycin | NP3 - M32 | 0.1906 | 0.2166 | 0.8800 | 0.8973 |
| Tobramycin | PAO1-1 - M32 | -1.7121 | 0.2008 | -8.5283 | 0.0000* |
| Tobramycin | PNM - M32 | -0.1606 | 0.3070 | -0.5231 | 0.9838 |
| Tobramycin | NP3 - NP1 | 0.8802 | 0.3875 | 2.2714 | 0.1428 |
| Tobramycin | PAO1-1 - NP1 | -1.0225 | 0.3789 | -2.6985 | 0.0490* |
| Tobramycin | PNM - NP1 | 0.5291 | 0.4531 | 1.1677 | 0.7560 |
| Tobramycin | PAO1-1 - NP3 | -1.9027 | 0.1965 | -9.6826 | 0.0000* |
| Tobramycin | PNM - NP3 | -0.3511 | 0.3051 | -1.1511 | 0.7656 |
| Tobramycin | PNM - PAO1-1 | 1.5515 | 0.2940 | 5.2765 | 0.0000* |

#### 11.13.3 T-test double antibiotic- and phage-resistant mutants

**Supplementary Table S8.** T-test for log_2_(EC50 FC) comparisons between single antibiotic-resistant and double antibiotic- and phage-resistant mutants. The pairwise T-test was used to asses differences in log_2_(EC50 FC) between the single antibiotic-resistant mutants and the double antibiotic- and phage resistant mutants. Test statistics, degrees of freedom, p-values, group means, and significance are reported. Significant comparisons are indicated by *.

| Antibiotic | t-statistic | df | p-value | Mean AB-resistant | Mean AB+P-resistant |
| --- | --- | --- | --- | --- | --- |
| Aztreonam | 2.054 | 9.350 | 0.069 | 2.416 | 1.841 |
| Ceftazidime | 0.513 | 9.644 | 0.619 | 4.438 | 4.275 |
| Ciprofloxacin | 1.915 | 4.376 | 0.122 | 4.225 | 3.347 |
| Colistin | 3.233 | 8.843 | 0.011* | 0.672 | -1.397 |
| Meropenem | 1.376 | 9.922 | 0.199 | 5.017 | 4.388 |
| Tobramycin | 3.549 | 7.765 | 0.008* | 2.475 | 0.574 |

#### 11.13.4 Partial Least Squares regression analysis

**Supplementary Figure S20.**
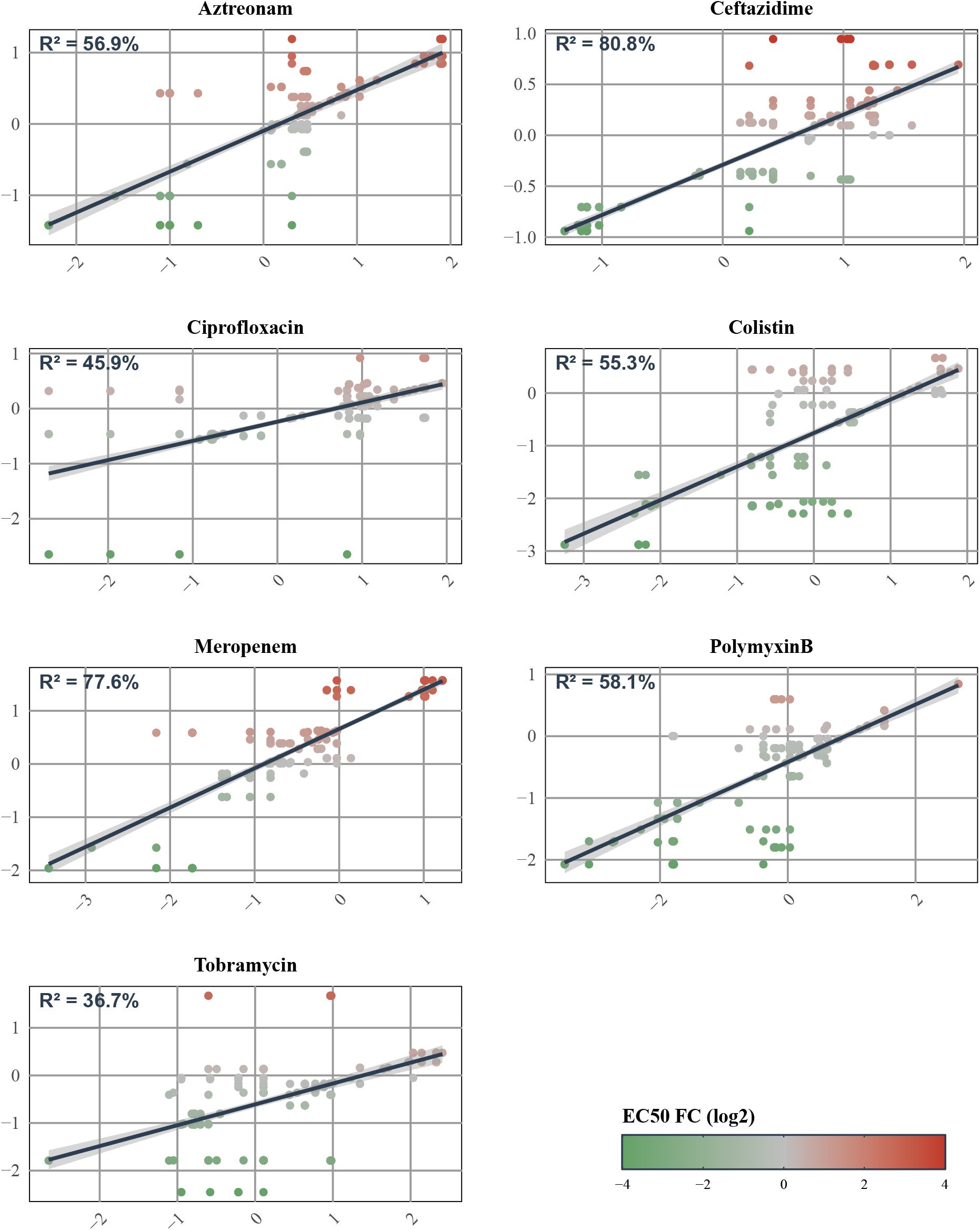
Partial Least Squares (PLS) regression analysis. The variance explained (R^2^) in log_2_(EC50_fc_) (y-axis) by gene identity per antibiotic, derived from PLS regression with a single latent component (x-axis).

**Supplementary Figure S21.**
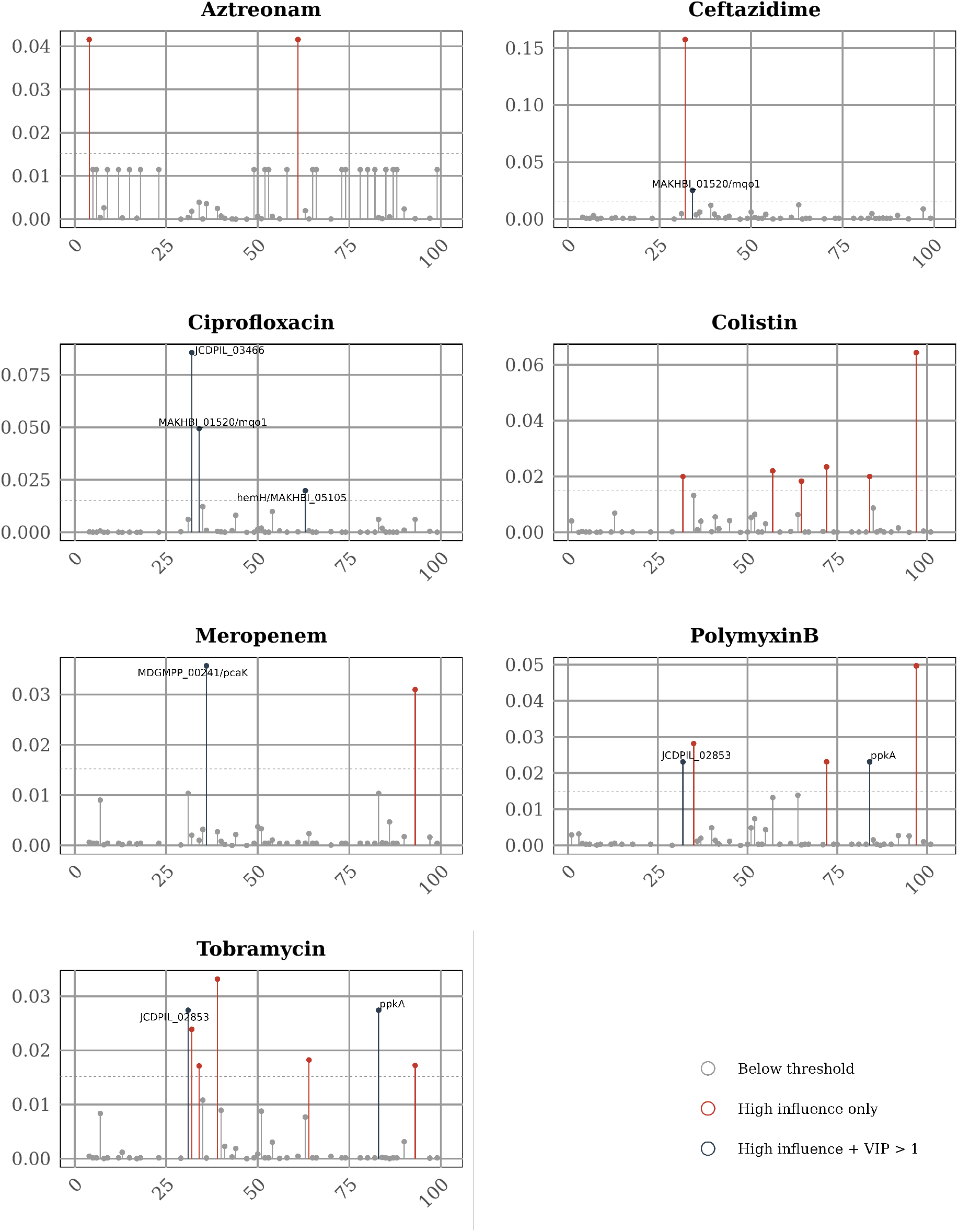
Mean Cook’s distance per gene per antibiotic. Mean Cook’s distance is plotted per gene, averaged across all mutants carrying mutations in that gene. The dashed line denotes the conventional influence threshold of 4/n, where n is the number of unique genes per antibiotic. Genes are colored by category: below threshold (grey), above threshold with VIP *≤* 1 (red; outlier-driven), and above threshold with VIP *>* 1 (dark; priority genes with confirmed PLS signal). Gene labels are shown for priority genes only.

### 11.14 Genetics mutations

**Supplementary Table S9.**
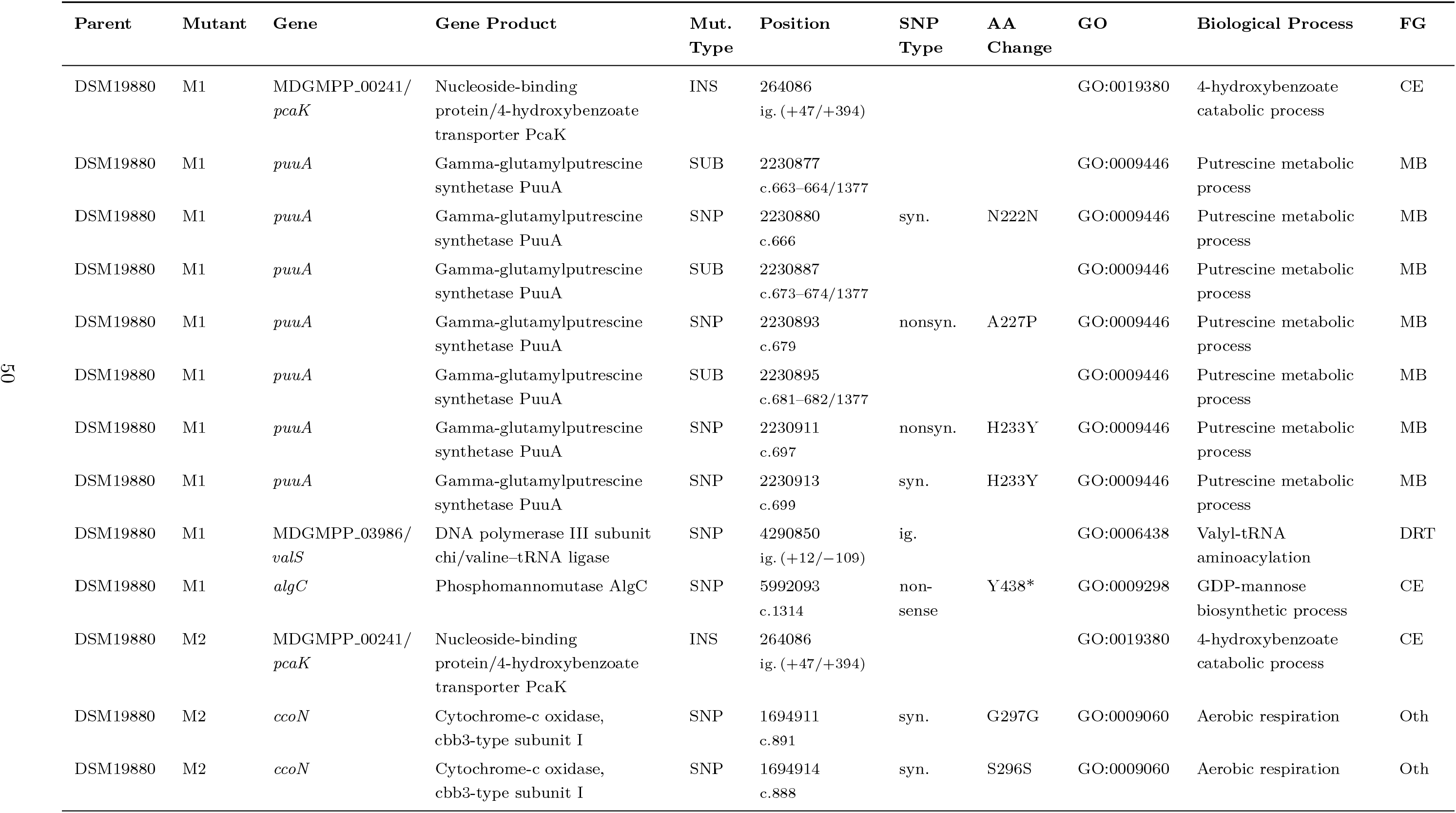
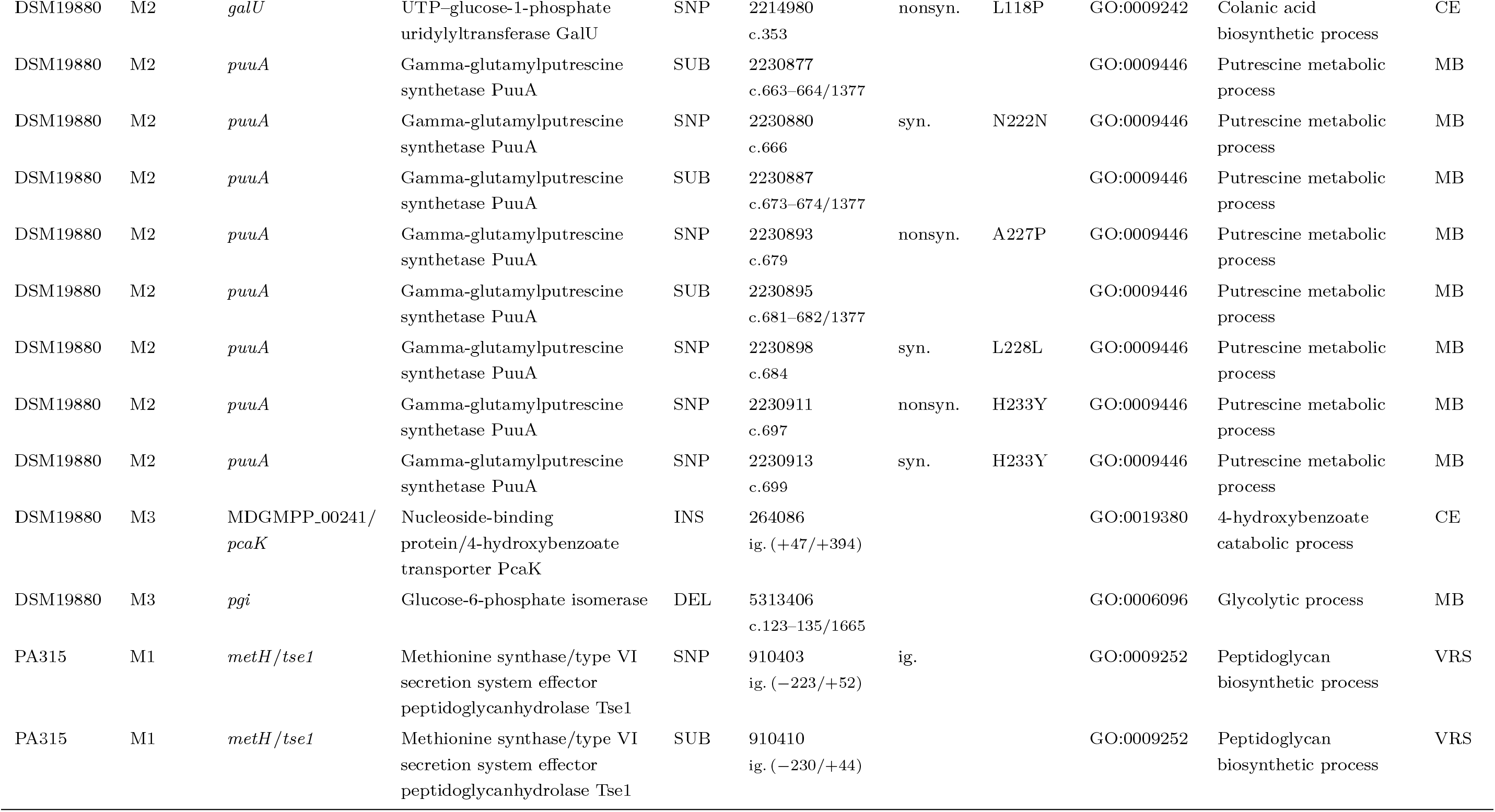
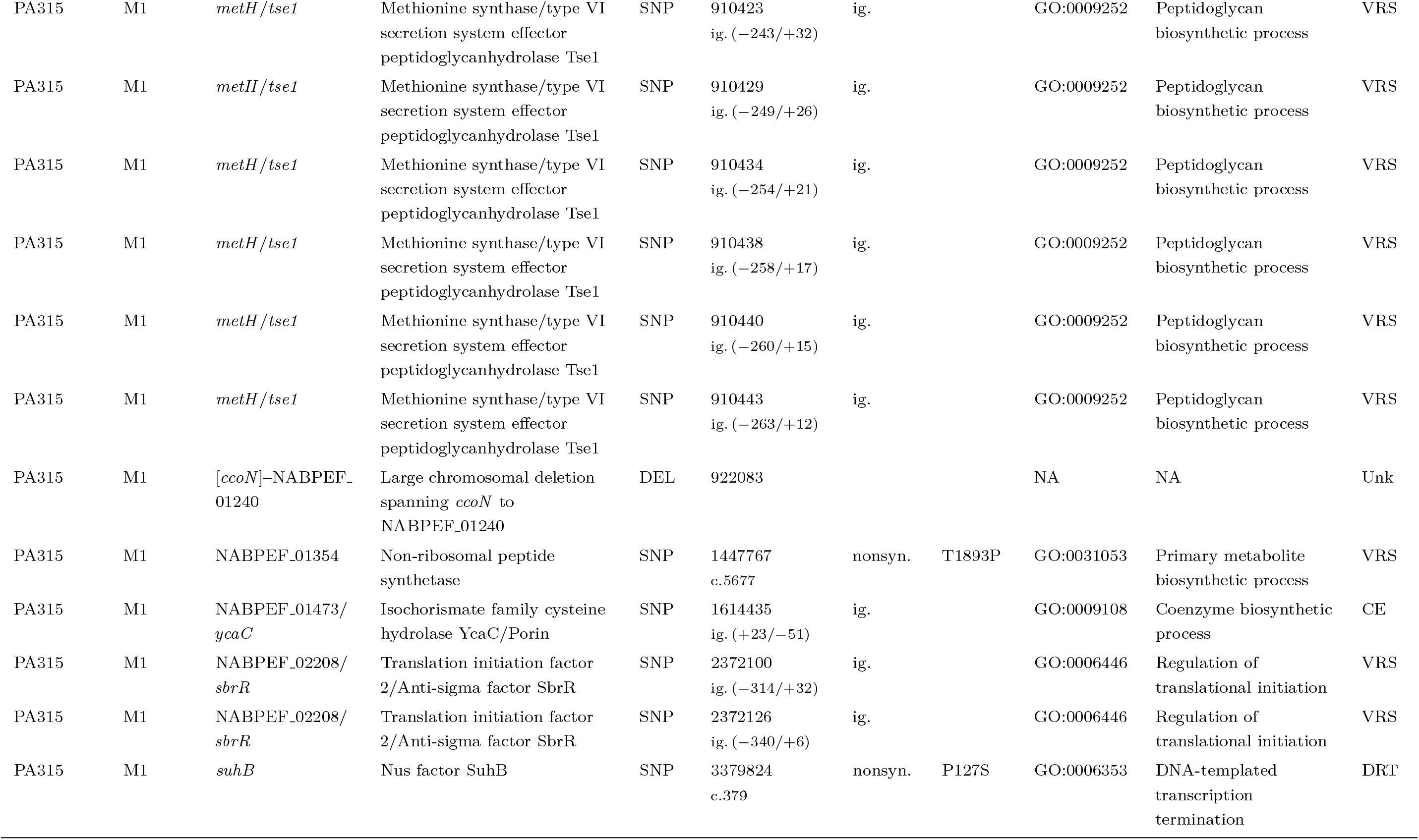
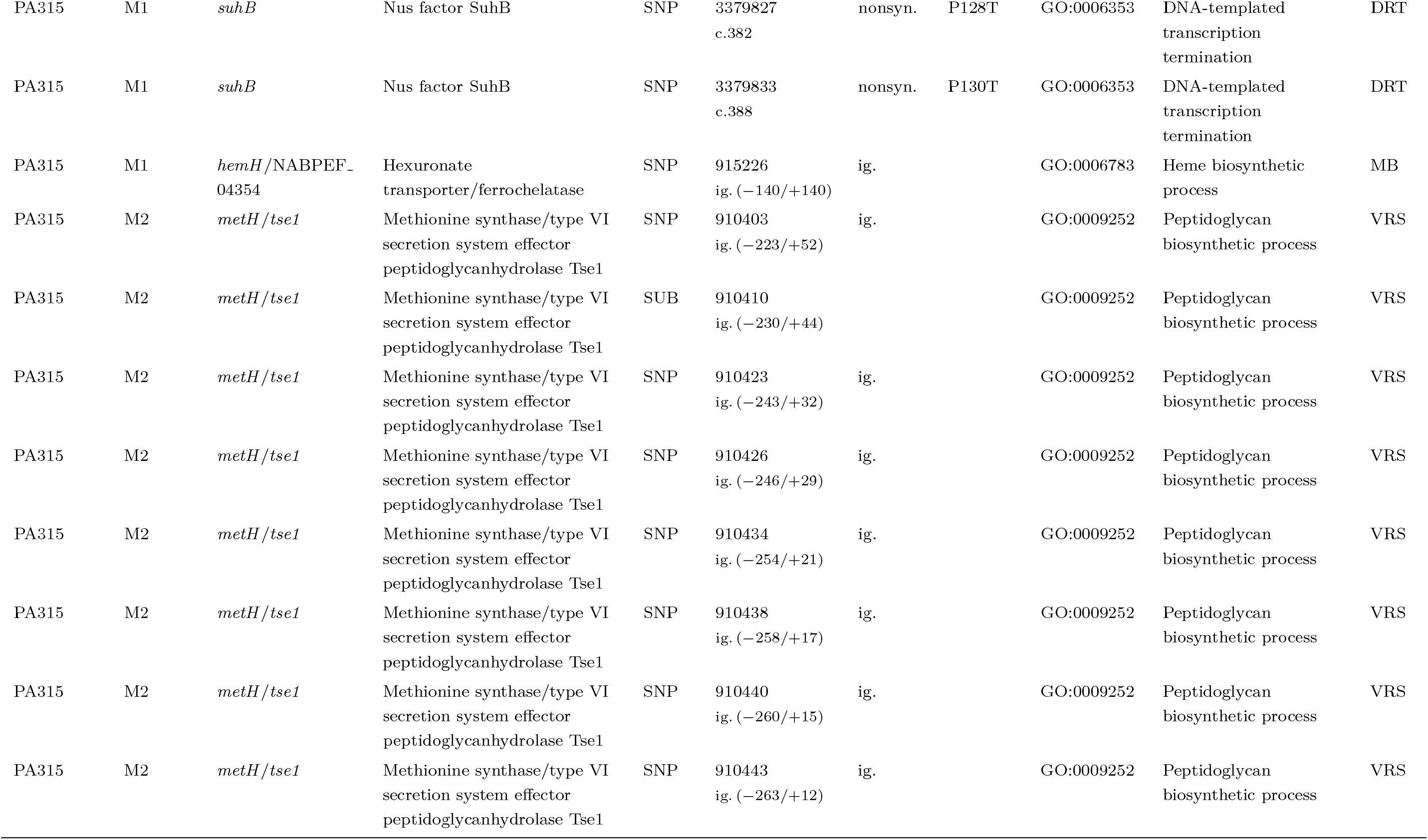
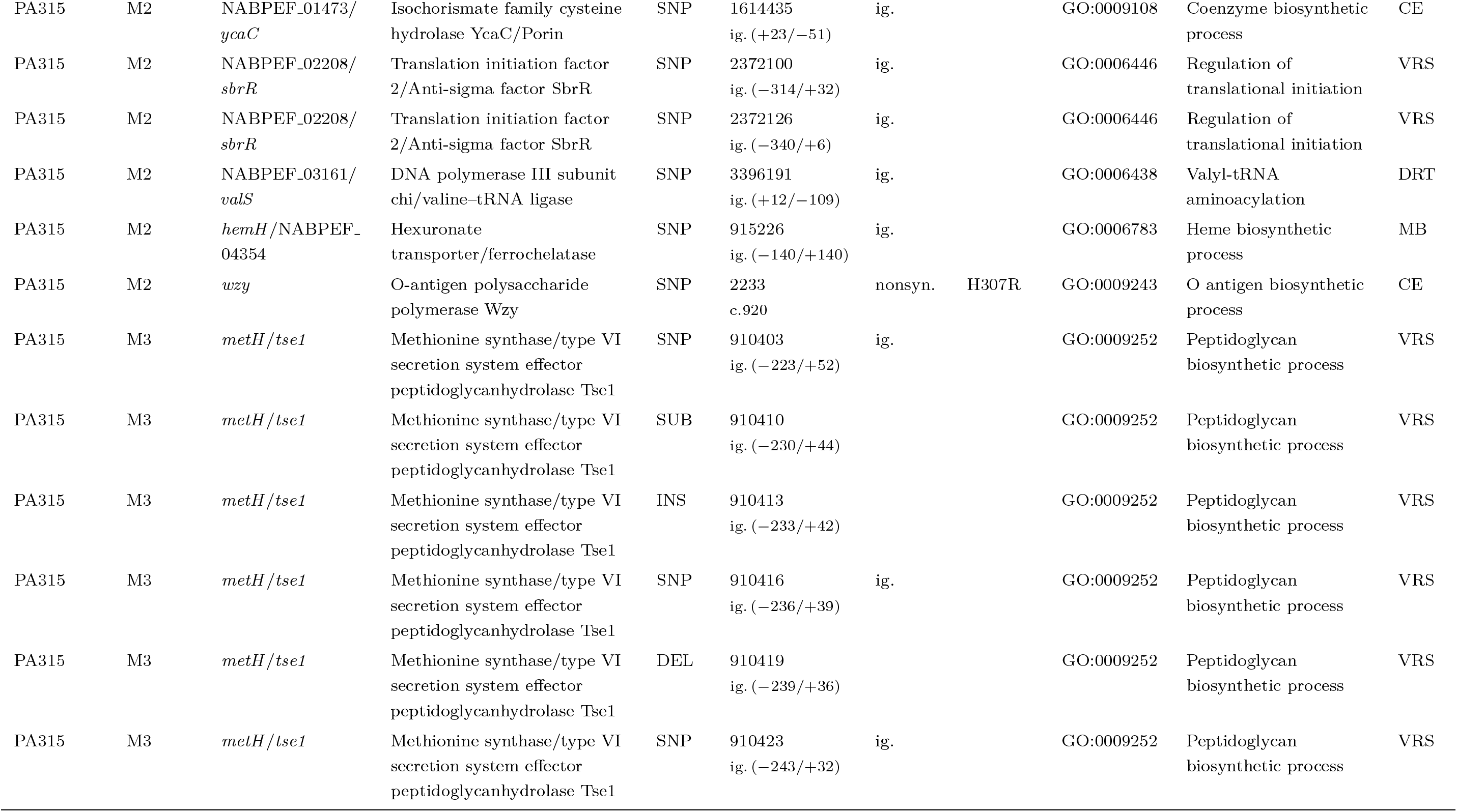
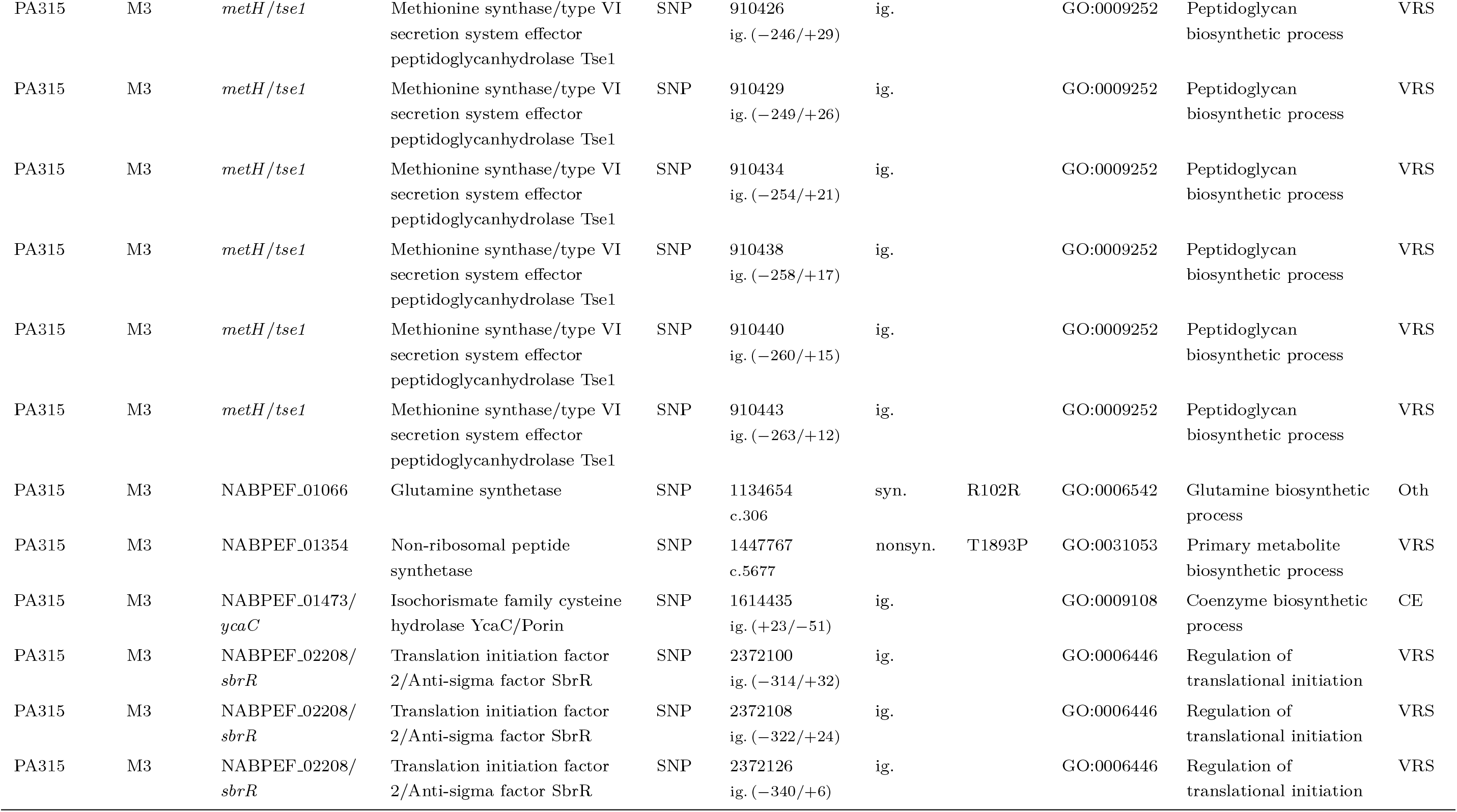
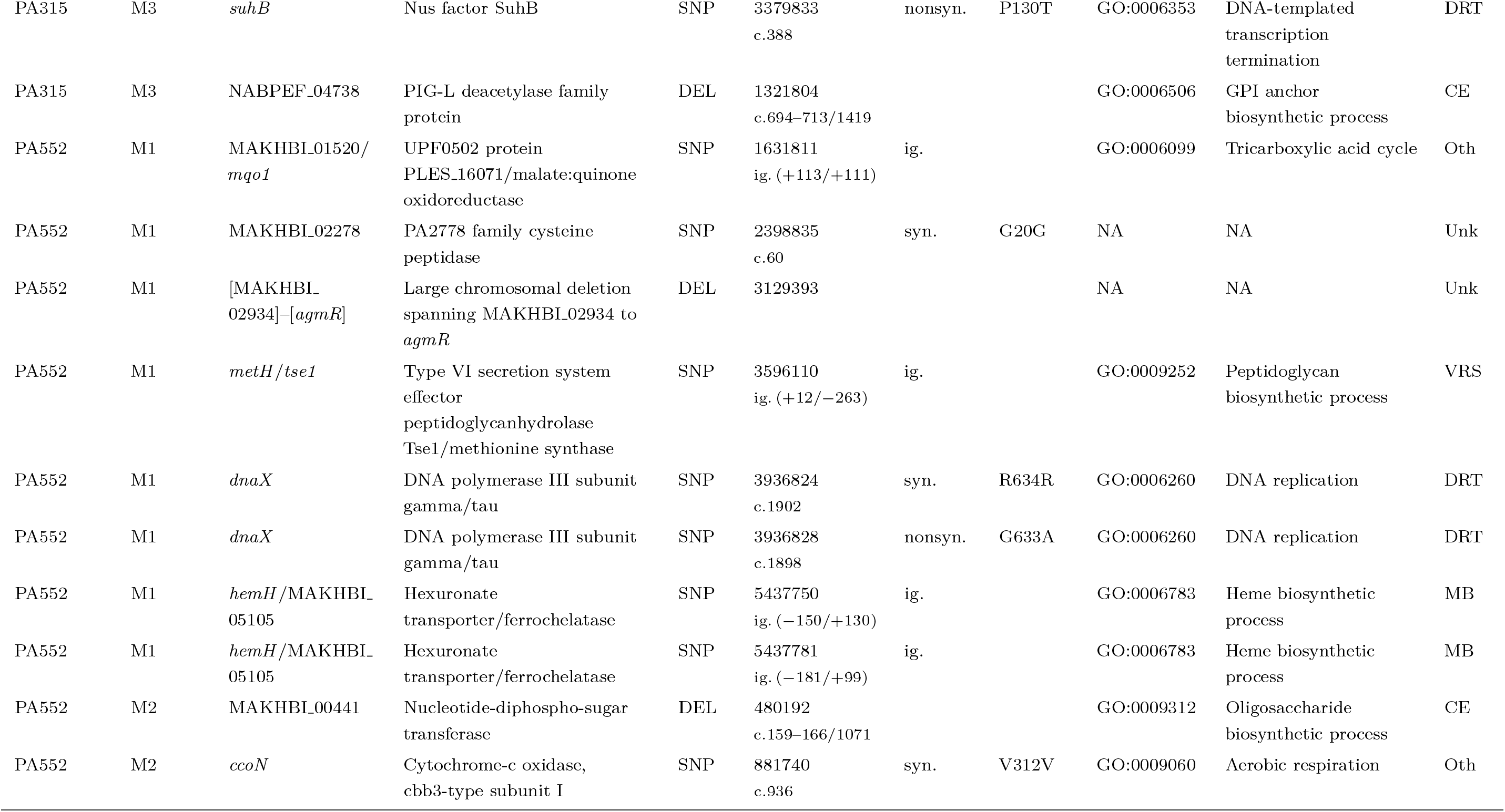
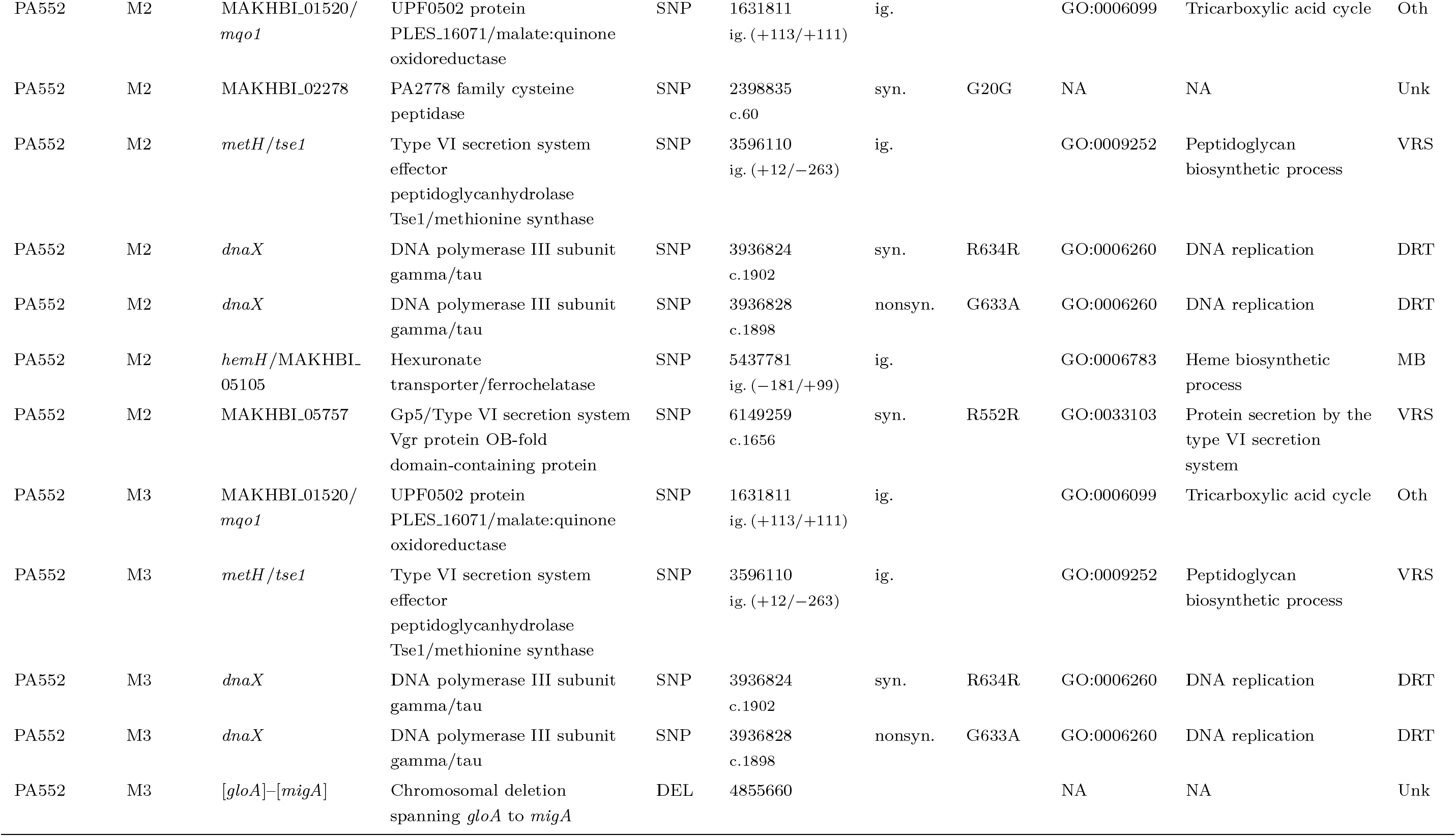
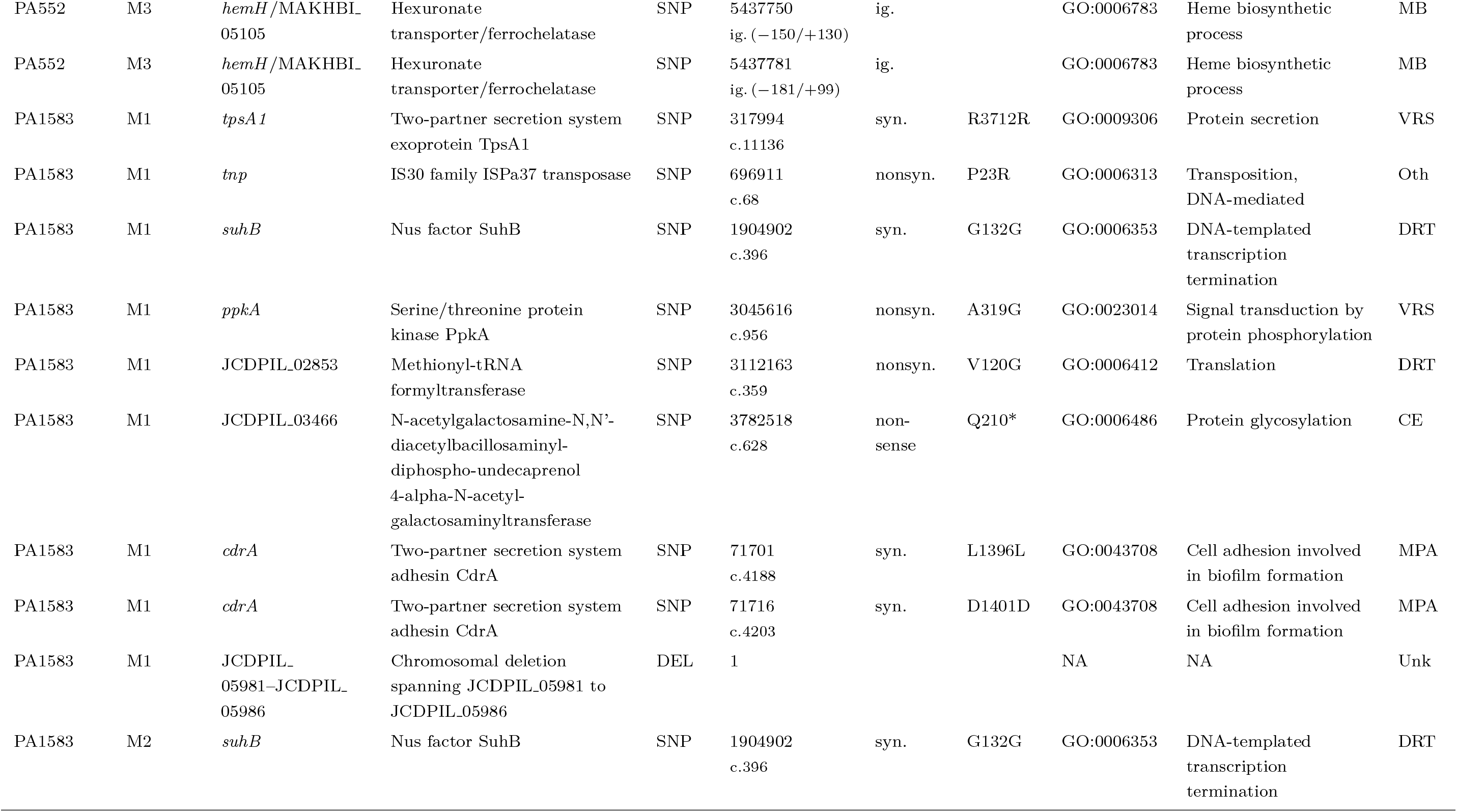
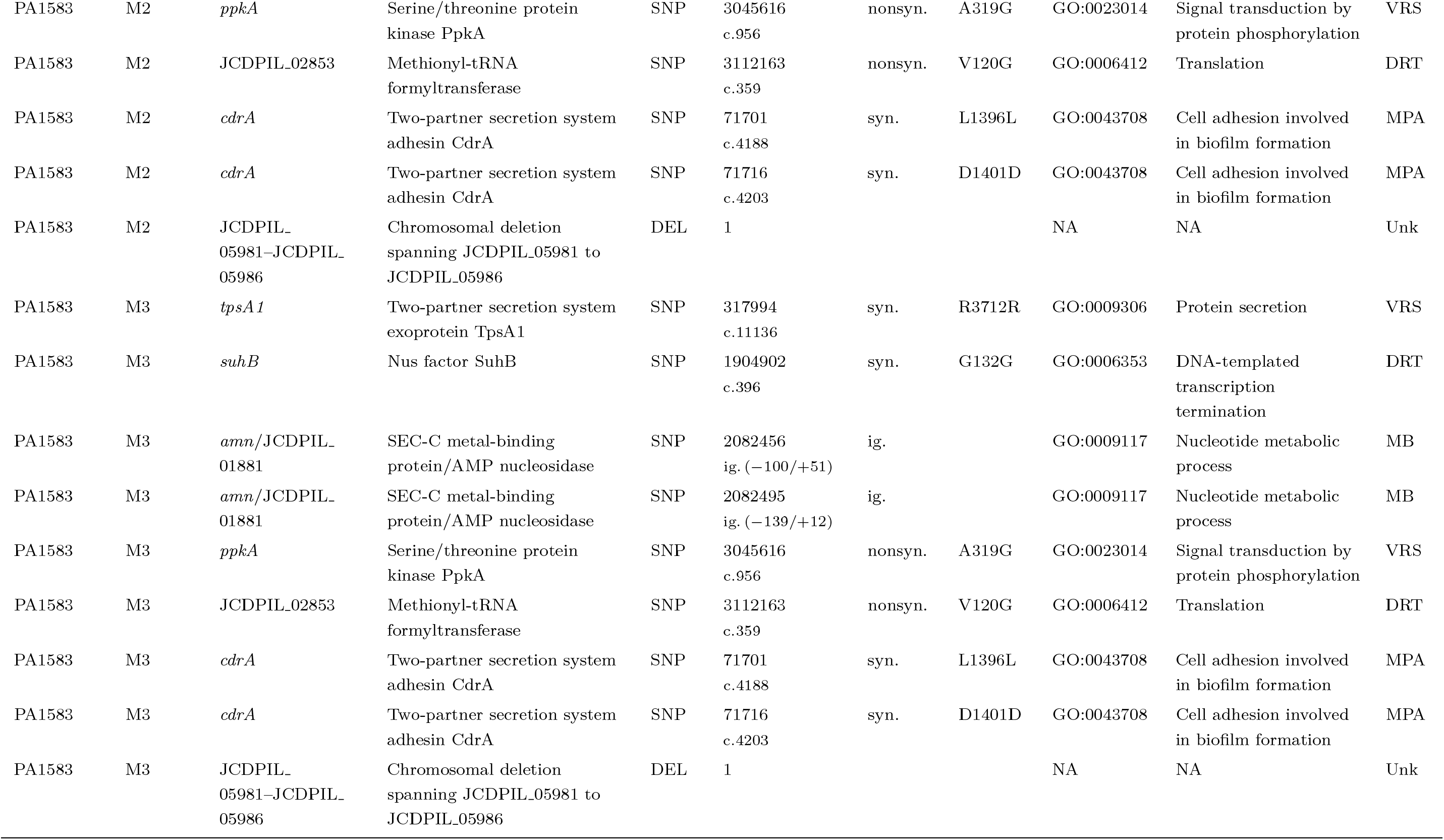
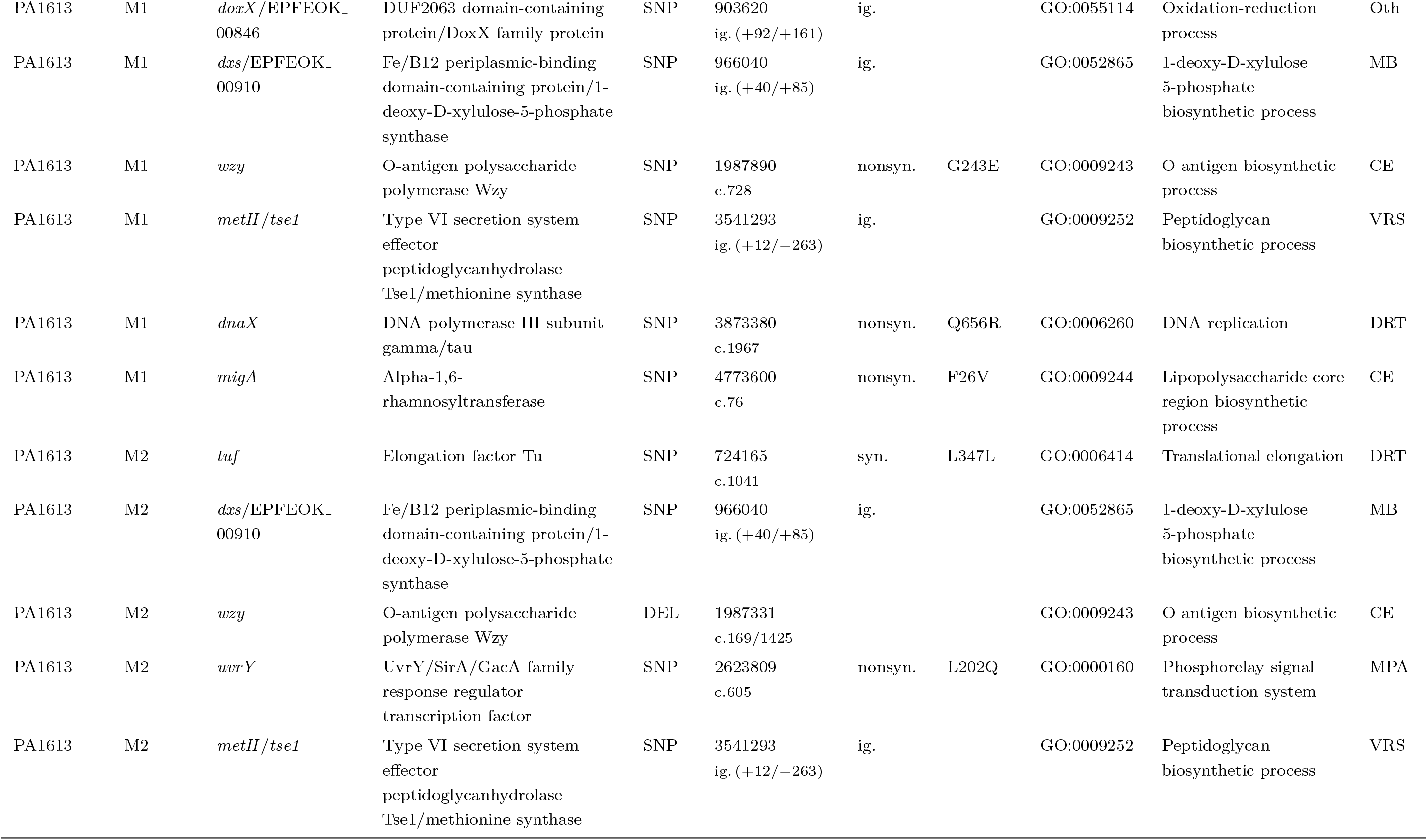
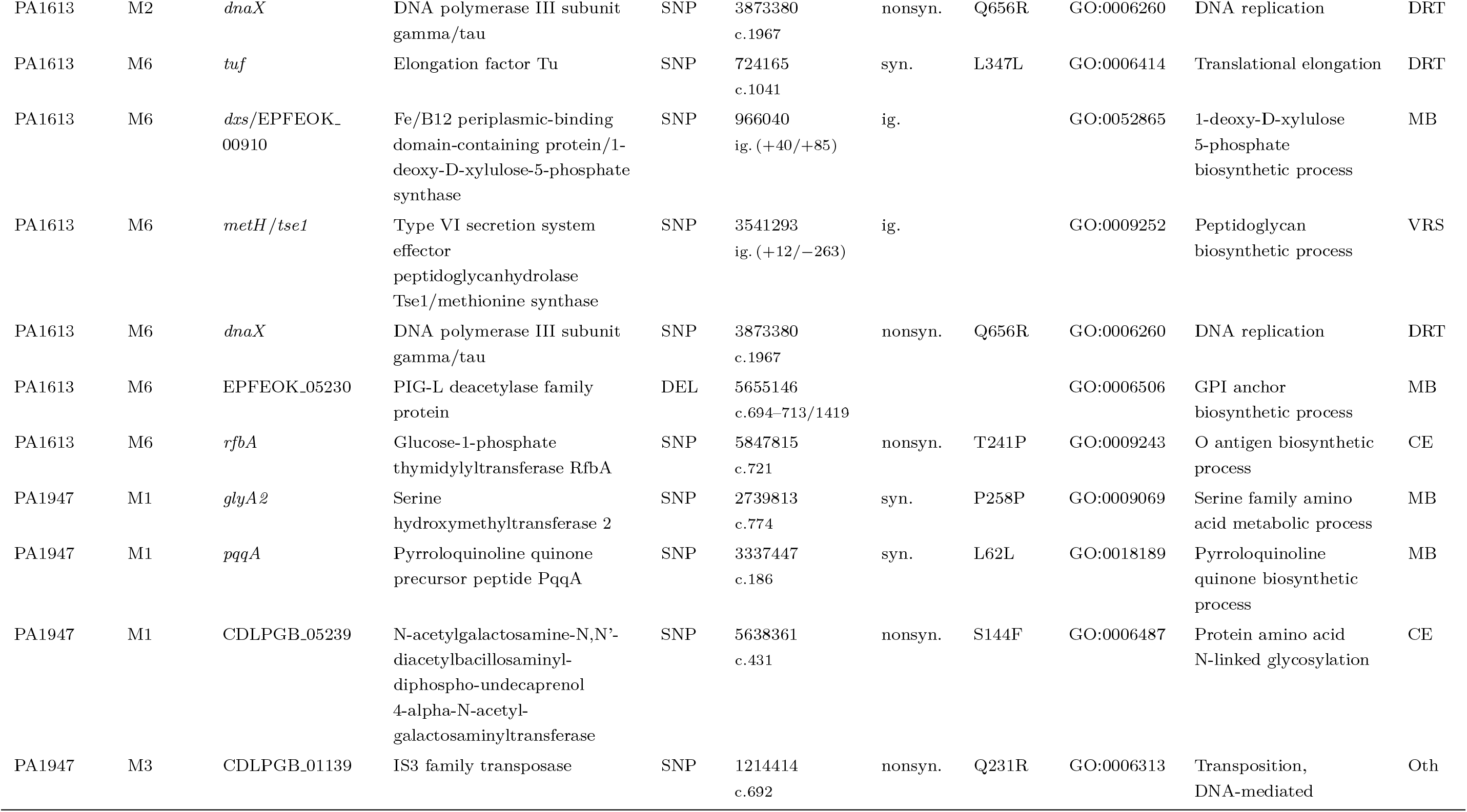
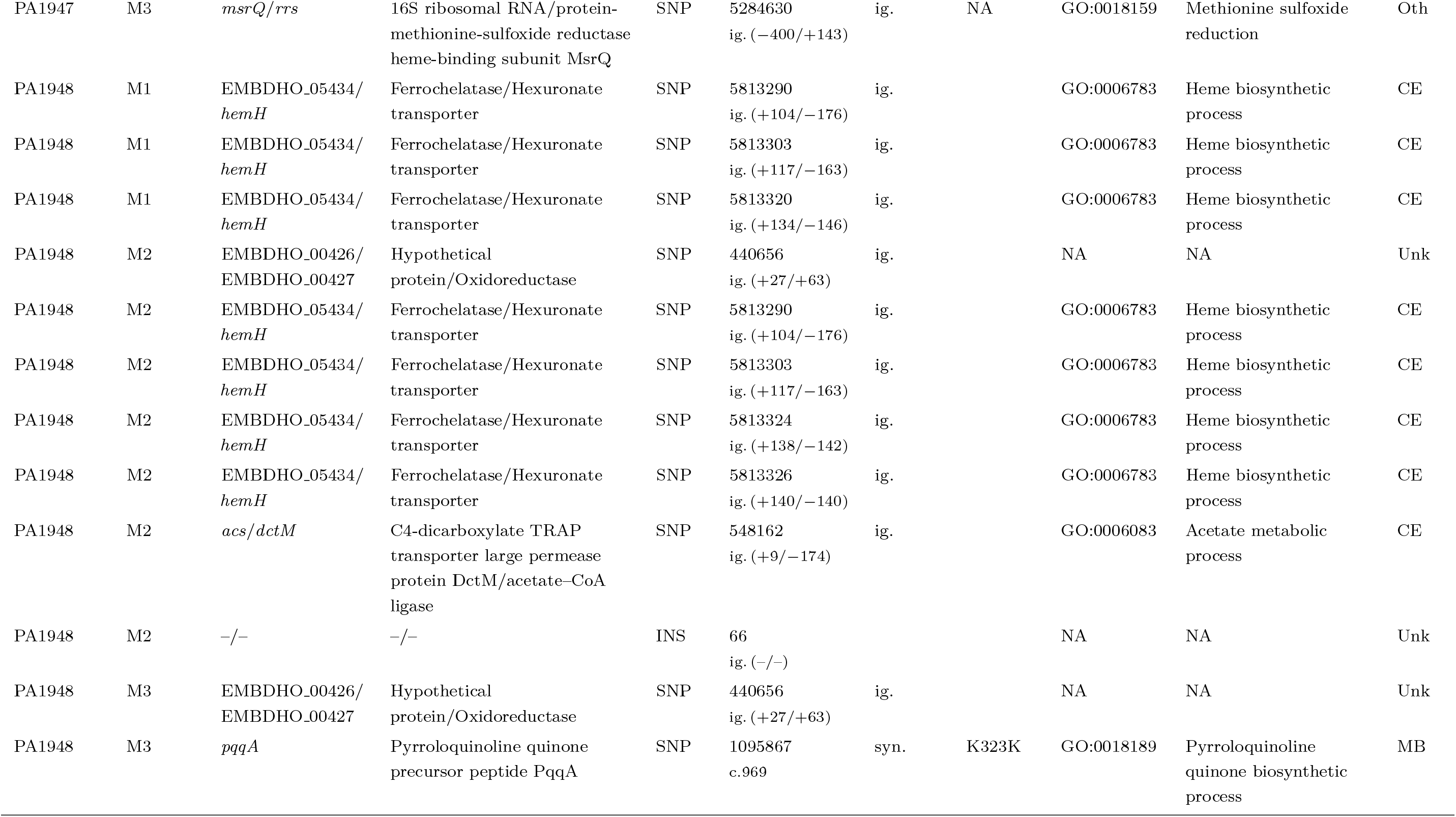
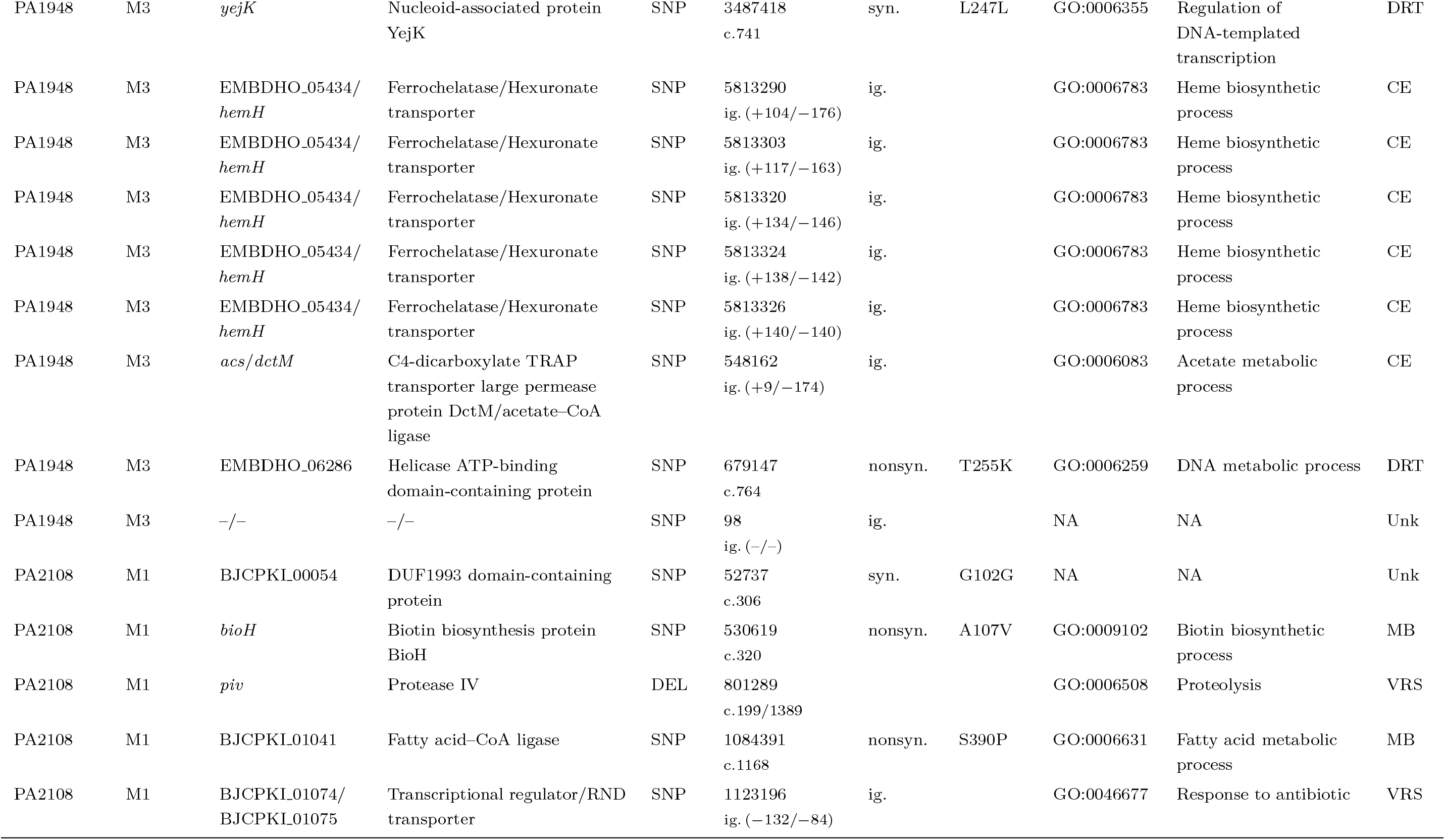
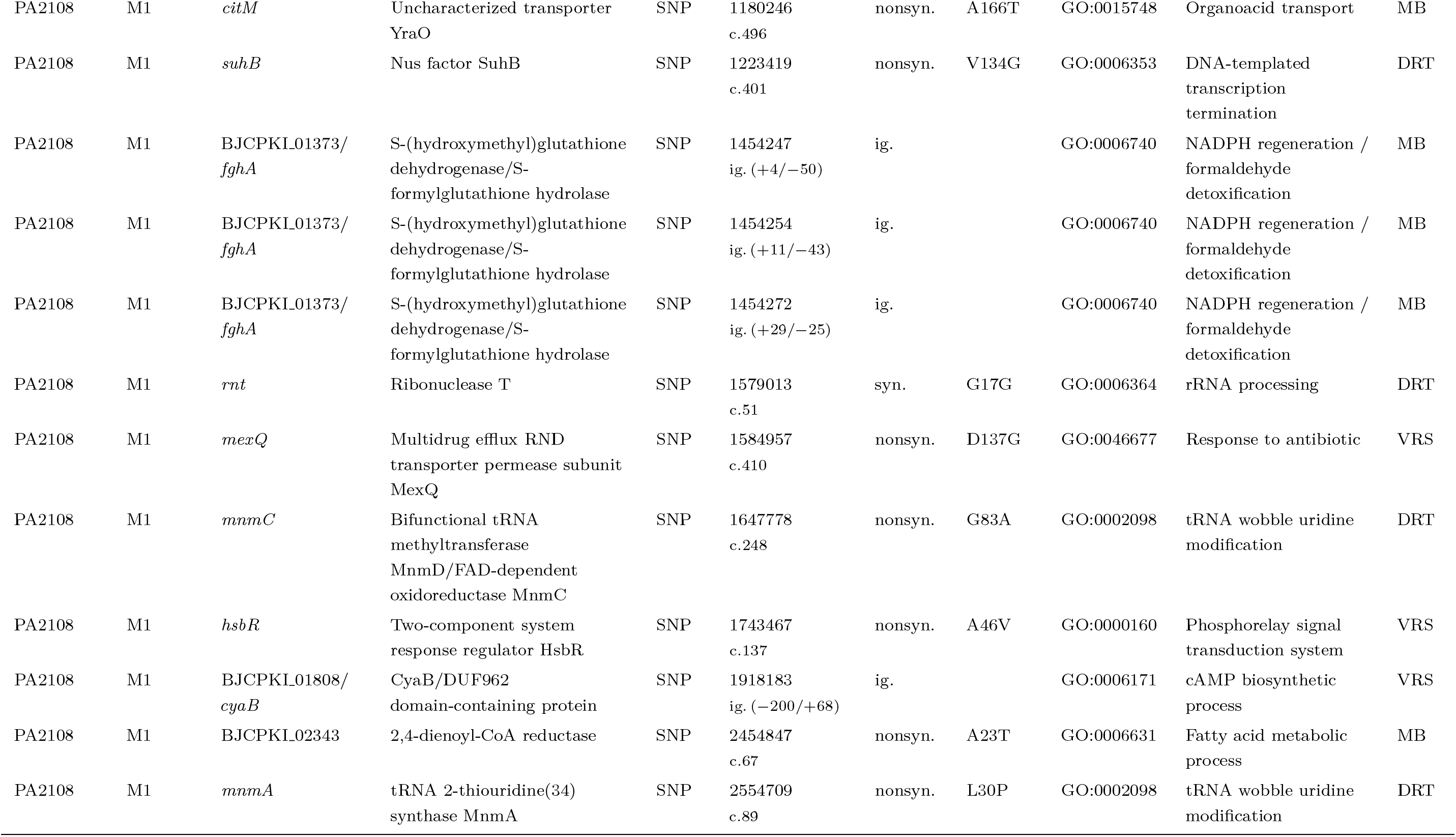
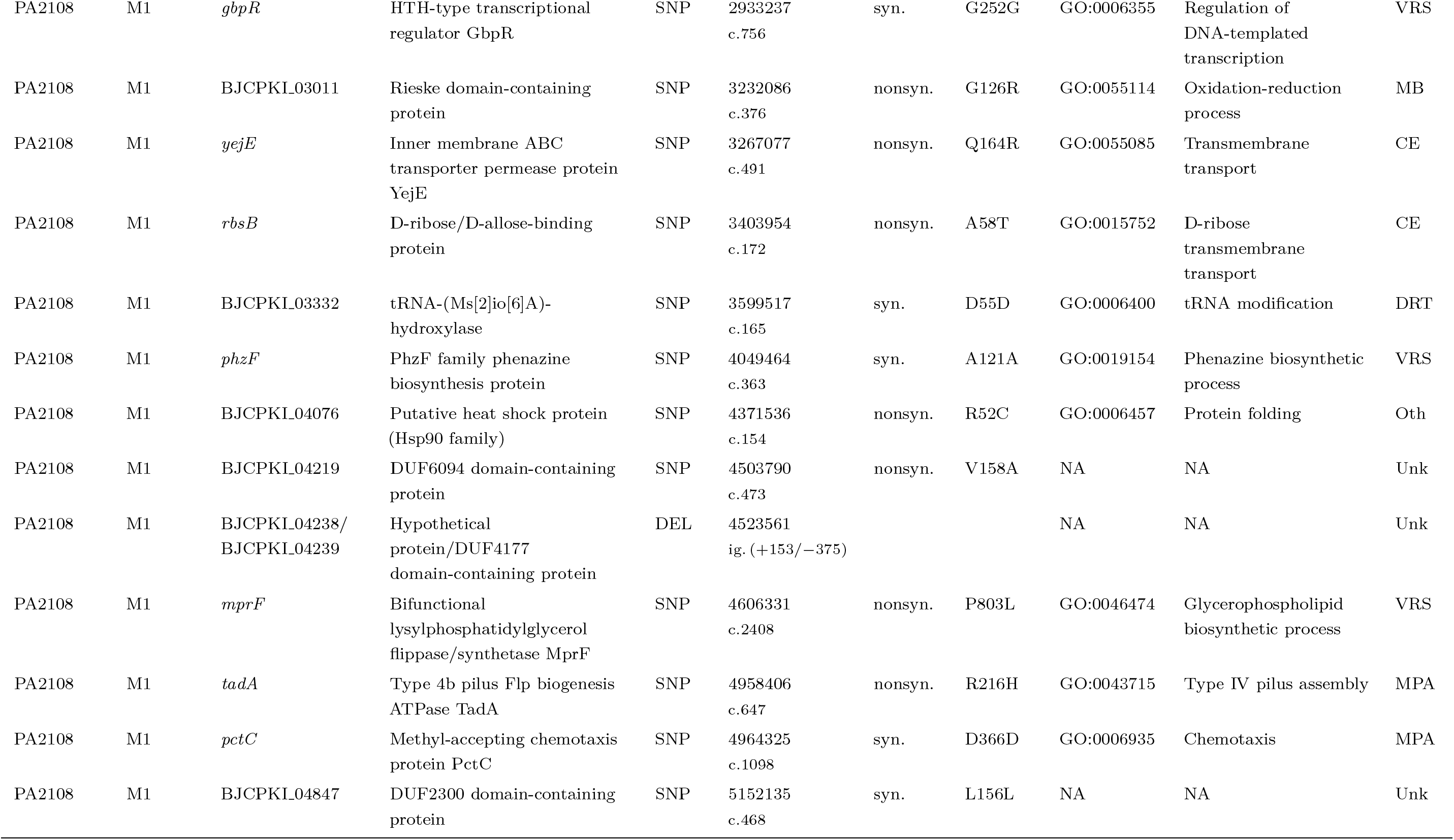
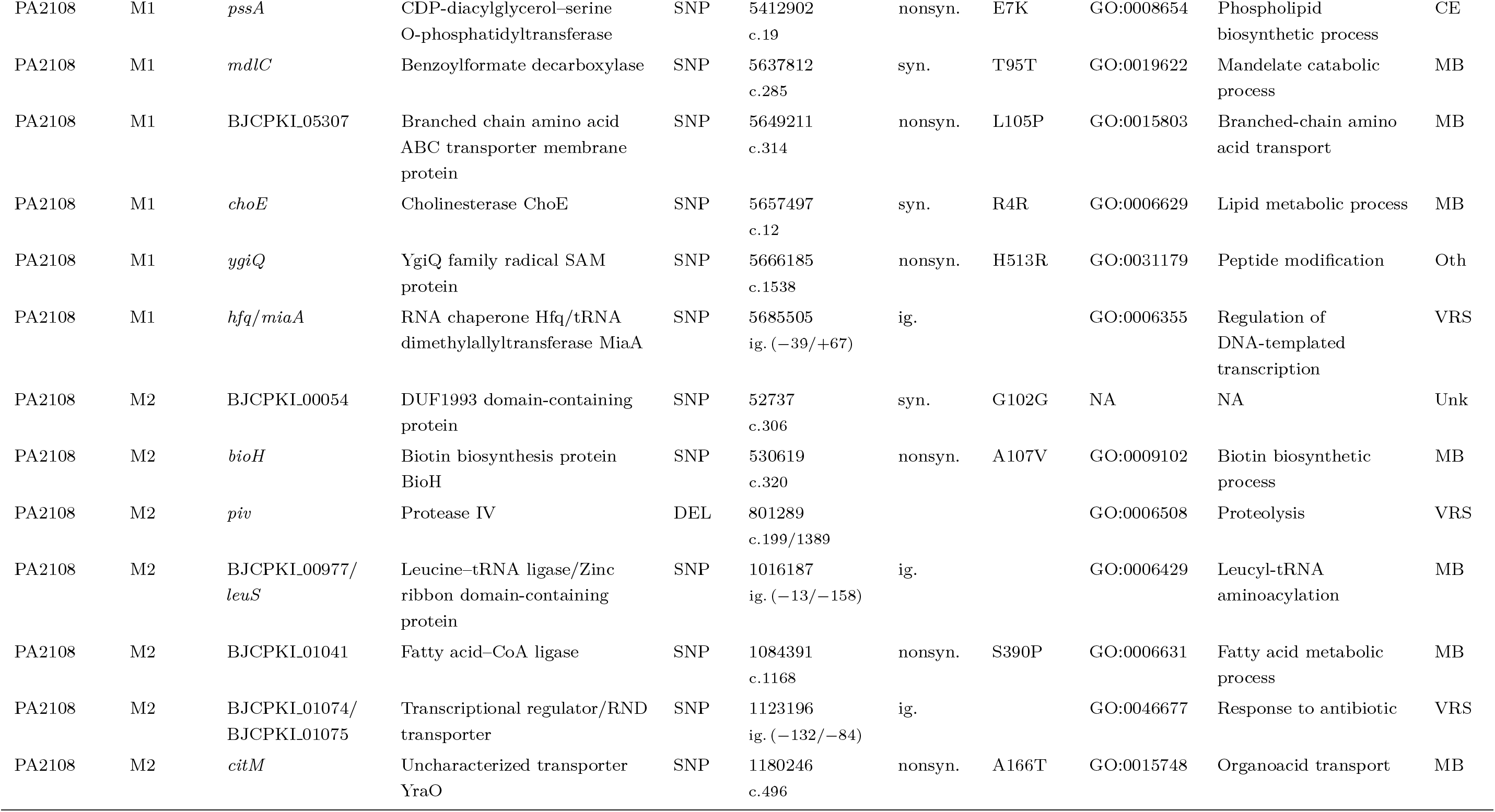
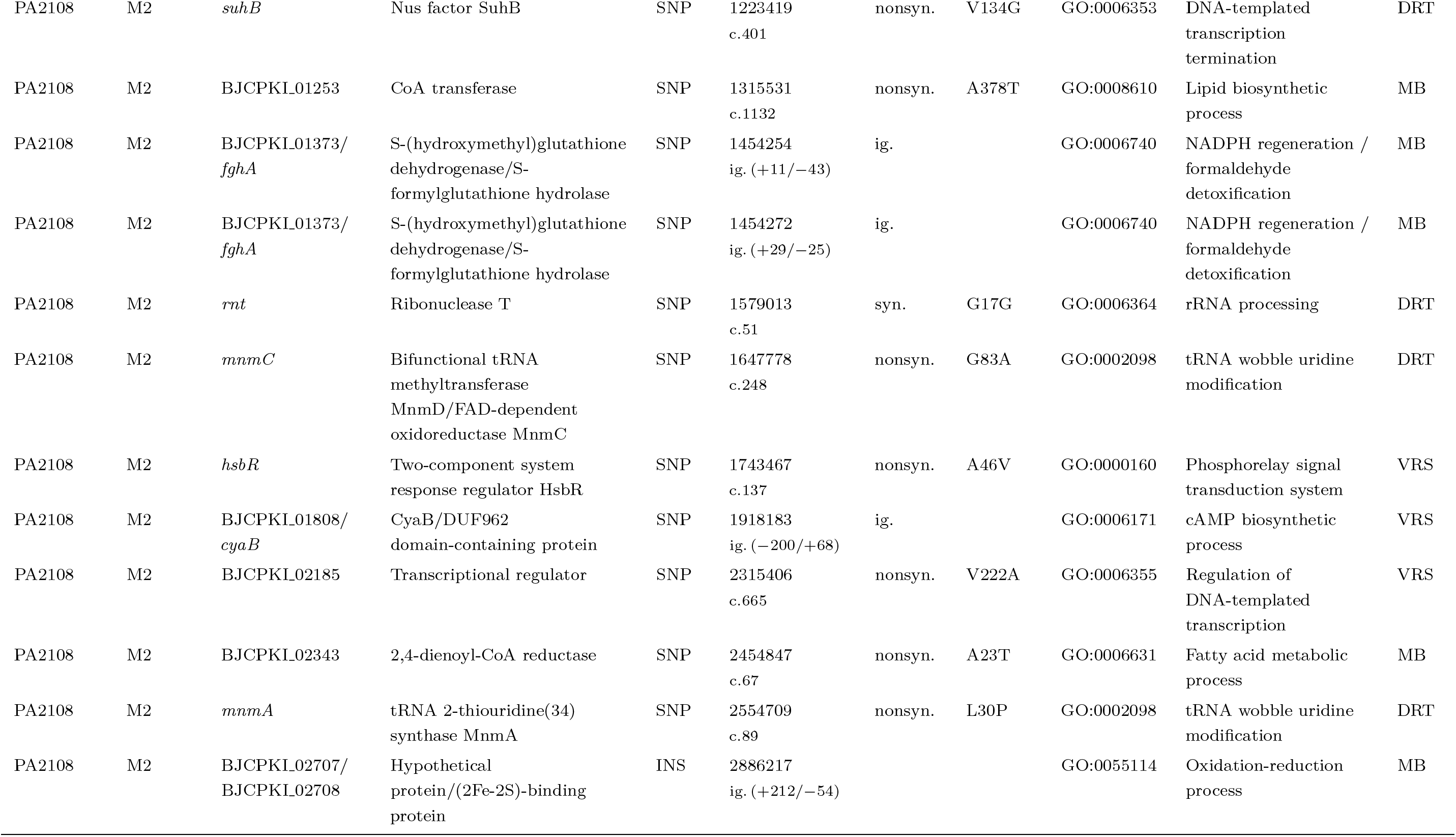
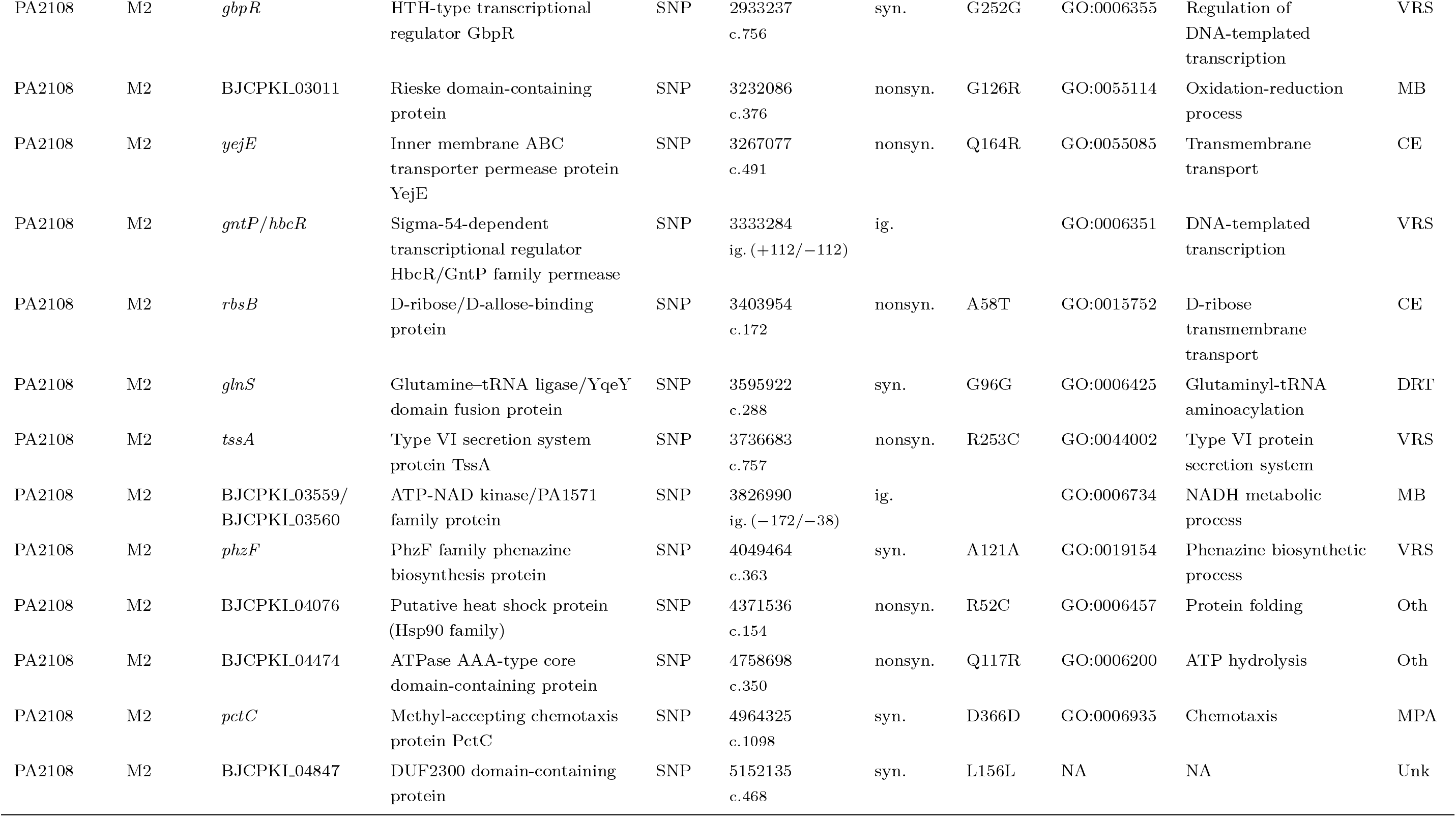
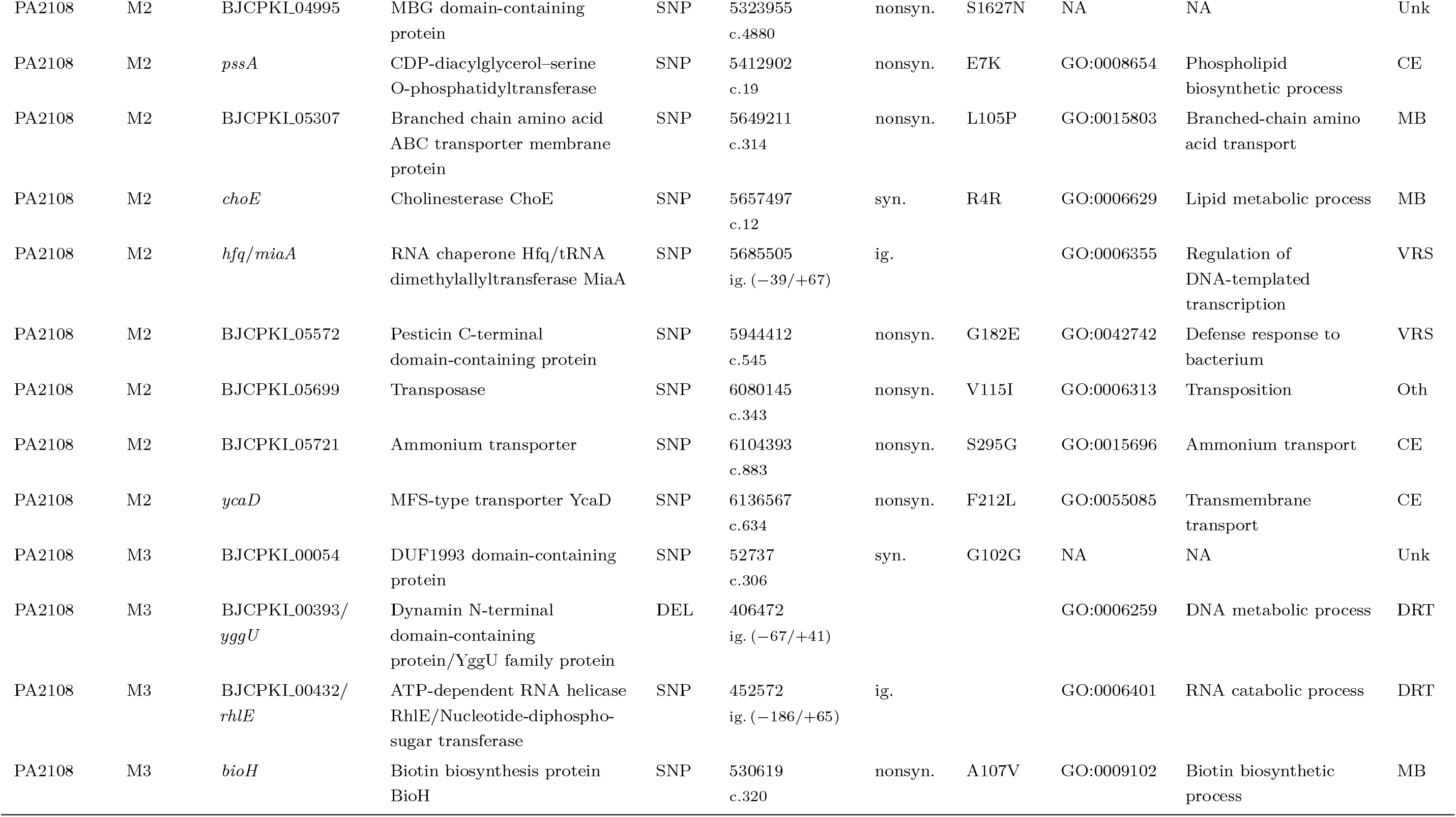
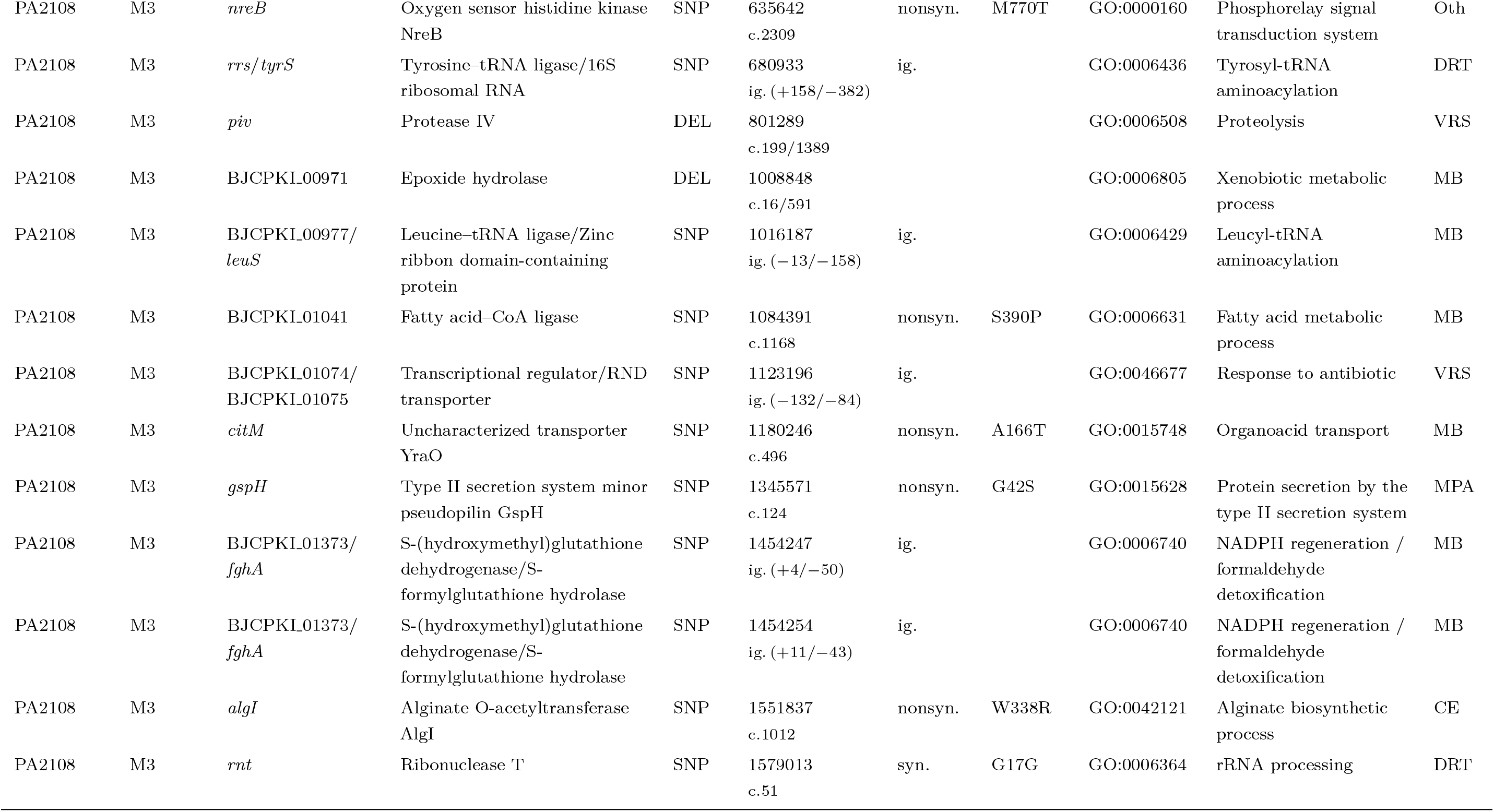
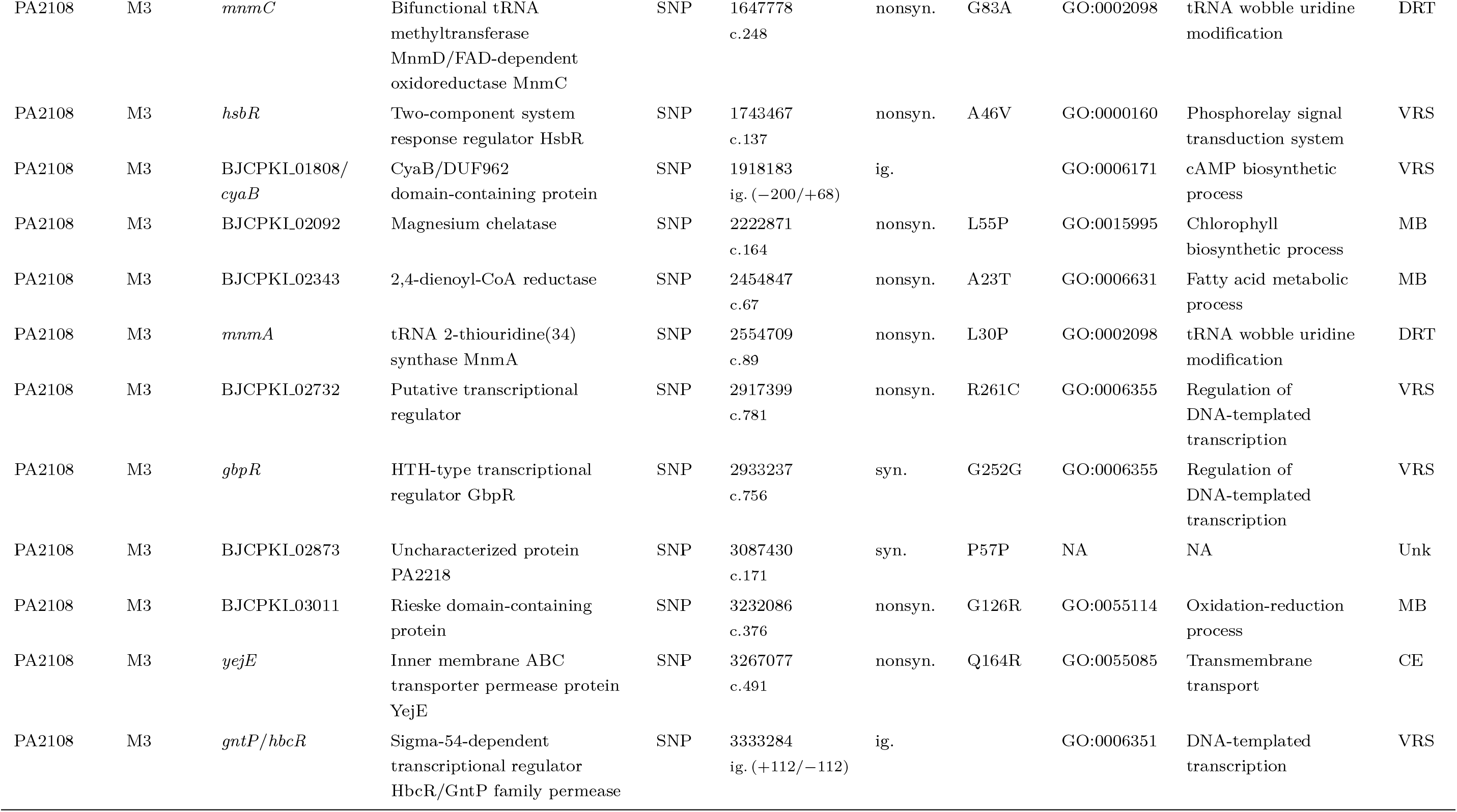
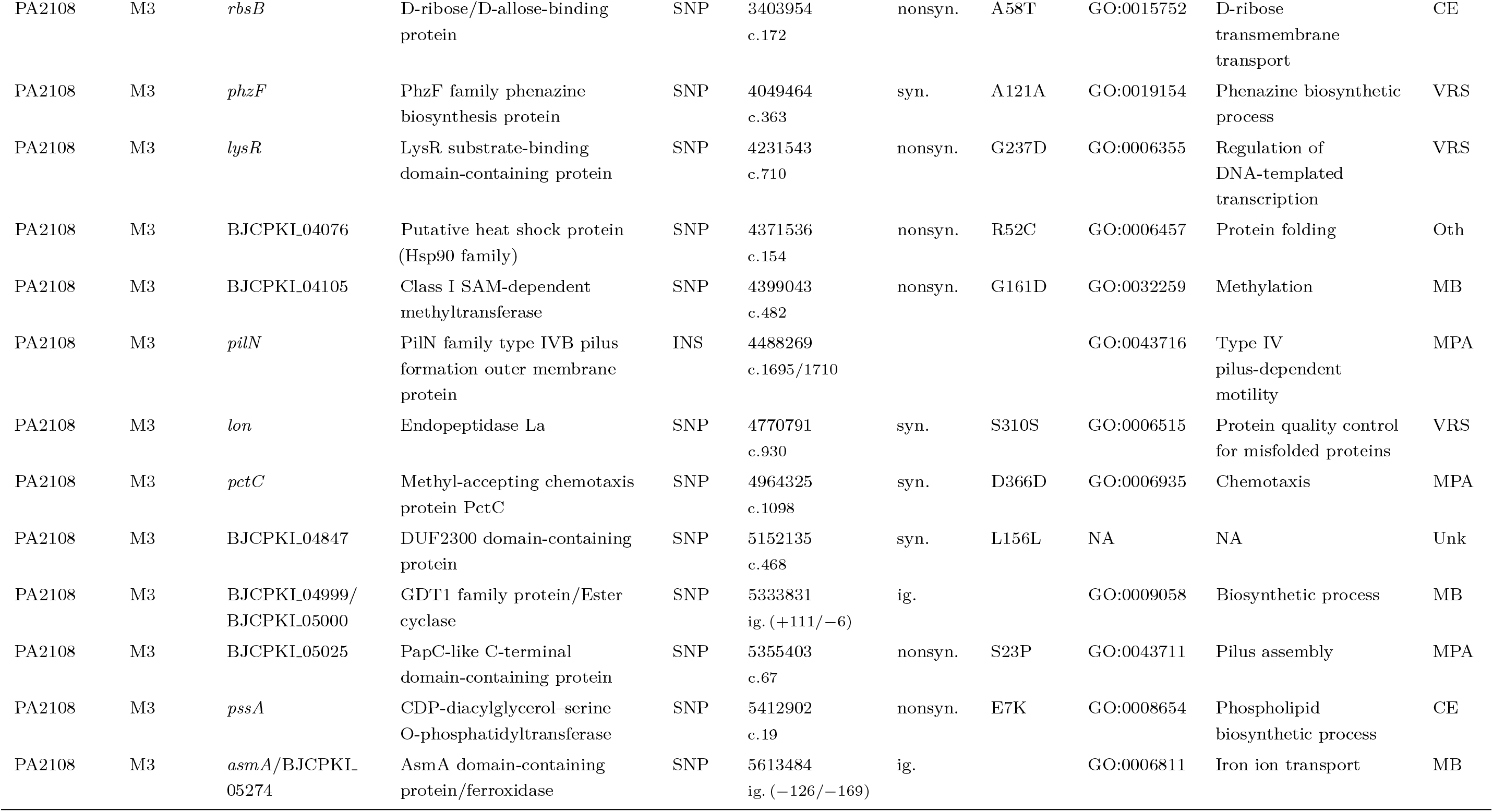
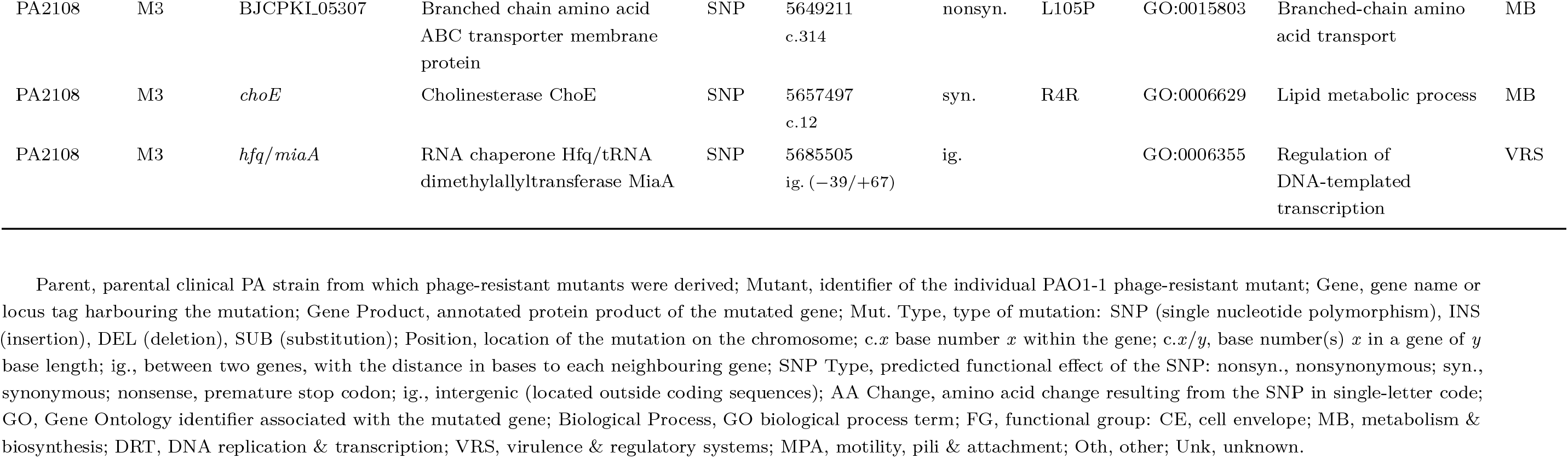
Complete overview of mutations identified in PAO1-1 phage-resistant mutants across parental lines. Parent, parental clinical PA strain from which phage-resistant mutants were derived; Mutant, identifier of the individual PAO1-1 phage-resistant mutant; Gene, gene name or locus tag harbouring the mutation; Gene Product, annotated protein product of the mutated gene; Mut. Type, type of mutation: SNP (single nucleotide polymorphism), INS (insertion), DEL (deletion), SUB (substitution); Position, location of the mutation on the chromosome; c.*x* base number *x* within the gene; c.*x*/*y*, base number(s) *x* in a gene of *y* base length; ig., between two genes, with the distance in bases to each neighbouring gene; SNP Type, predicted functional effect of the SNP: nonsyn., nonsynonymous; syn., synonymous; nonsense, premature stop codon; ig., intergenic (located outside coding sequences); AA Change, amino acid change resulting from the SNP in single-letter code; GO, Gene Ontology identifier associated with the mutated gene; Biological Process, GO biological process term; FG, functional group: CE, cell envelope; MB, metabolism & biosynthesis; DRT, DNA replication & transcription; VRS, virulence & regulatory systems; MPA, motility, pili & attachment; Oth, other; Unk, unknown.

